# Sequence sensitivity of open chromatin reveals constrained regulatory elements underlying human traits

**DOI:** 10.1101/2024.10.31.621195

**Authors:** Kohei Tomizuka, Masaru Koido, Akari Suzuki, Soichiro Yoshino, Nao Tanaka, Yuki Ishikawa, Xiaoxi Liu, Satoshi Koyama, Kazuyoshi Ishigaki, Yasuhiro Murakawa, Immune transcript/enhancer consortium (ITEC), Kazuhiko Yamamoto, Chikashi Terao

**Affiliations:** Laboratory for Statistical and Translational Genetics, RIKEN Center for Integrative Medical Sciences, 1-7-22 Suehiro-cho, Tsurumi-ku, Yokohama City, Kanagawa, 230-0045, Japan; Clinical Research Center, Shizuoka General Hospital, 4 Chome-27-1 Kitaando, Aoi Ward, Shizuoka, 420-8527, Japan; Department of Statistical and Translational Genetics, Center for Genomic Medicine, Fujita Health University, 1-98 Dengakugakubo, Kutsukake-cho,Toyoake, Aichi 470-1192, Japan; Division of Neuropsychiatric Statistics and AI, International Center for Brain Science, Fujita Health University, 1-98 Dengakugakubo, Kutsukake-cho,Toyoake, Aichi 470-1192, JAPAN; Laboratory of Complex Trait Genomics, Department of Computational Biology and Medical Sciences, Graduate School of Frontier Sciences, The University of Tokyo, 4-6-1, Shirokane-dai, Minato-ku, Tokyo, 108-8639, Japan; Laboratory for Autoimmune Diseases, RIKEN Center for Integrative Medical Sciences, 1-7-22 Suehiro-cho, Tsurumi-ku, Yokohama City, Kanagawa, 230-0045, Japan; Laboratory for Cardiovascular Genomics and Informatics, RIKEN Center for Integrative Medical Sciences, 1-7-22 Suehiro-cho, Tsurumi-ku, Yokohama City, Kanagawa, 230-0045, Japan; Laboratory for Human Immunogenetics, RIKEN Center for Integrative Medical Sciences, 1-7-22 Suehiro-cho, Tsurumi-ku, Yokohama City, Kanagawa, 230-0045, Japan; RIKEN-IFOM Joint Laboratory for Cancer Genomics, RIKEN Center for Integrative Medical Sciences, 1-7-22 Suehiro-cho, Tsurumi-ku, Yokohama City, Kanagawa, 230-0045, Japan; Drug Discovery Antibody Platform Unit, RIKEN Center for Integrative Medical Sciences, 1-7-22 Suehiro-cho, Tsurumi-ku, Yokohama City, Kanagawa, 230-0045, Japan; Department of Applied Genetics, The School of Pharmaceutical Sciences, University of Shizuoka, 52-1 Yada, Suruga Ward, Shizuoka, 422-8526, Japan

## Abstract

Open chromatin regions (OCRs) delineate cell-type-specific regulatory landscapes across the genome, yet their functional importance varies considerably. Here, we introduce CAMBUS (Chromatin Accessibility Mutation Burden Score), a machine-learning framework that quantifies the sensitivity of predicted chromatin accessibility to sequence perturbation to prioritize functionally important OCRs. Applying CAMBUS to 29 immune cell types, we identified 66,043 CAMBUS-defined OCRs (cOCRs), without directly incorporating human genetic variation or existing evolutionary constraint metrics. Nevertheless, cOCRs were strongly enriched in independently defined constrained regions (odds ratio (OR)=11.45 (95% confidence interval (CI) 9.33–14.05), P=4.7 × 10^-68^), and active regulatory elements, with enrichment for super-enhancers remaining after adjustment for established constraint metrics and sequence features (OR=6.18 (95% CI 3.80–10.05), P=2.1 × 10^-13^). cOCRs also showed cell-type-specific enrichment for the heritability of immune-mediated diseases and for putative causal variants identified by statistical fine-mapping across European and East Asian ancestries, including rare variants. Integration with statistical fine-mapping further linked 1,533 putative causal variants across 70 complex traits to cell-type-specific cOCRs, revealing plausible regulatory mechanisms underlying complex traits. Together, CAMBUS provides a region-centered framework for identifying regulatory elements whose predicted activity is sensitive to sequence disruption and that are enriched for evolutionary constraint, complementing existing regulatory and constraint annotations for interpreting non-coding genetic variation.

## Introduction

Genome sequences encode the information underlying species identity, evolution, and biological complexity ^1,2^. Yet protein-coding regions account for only a small fraction of vertebrate genomes ^3^, highlighting the importance of understanding the functions of non-coding regions, particularly those with regulatory activity ^4,5^. Active regulatory regions, such as enhancers, control gene expression in a cell-type- and state-specific manner and thereby contribute to diverse cellular functions and phenotypes ^6–9^.

Several genome-wide experimental approaches have been developed to identify candidate regulatory regions, including chromatin immunoprecipitation followed by sequencing (ChIP-seq) for histone modifications, DNase I hypersensitive site sequencing (DNase-seq), and assay for transposase-accessible chromatin using sequencing (ATAC-seq). Among these approaches, ATAC-seq has been widely adopted because of its technical simplicity and robust performance ^10,11^. Compared with histone modification ChIP-seq peaks, ATAC-seq-defined open chromatin regions (OCRs) provide high positional resolution for localizing candidate regulatory elements ^12^. Mapping OCRs across cell types and states has therefore provided important insights into shared and context-specific gene regulatory mechanisms ^13,14^. However, chromatin accessibility reflects the permissiveness of a genomic region to regulatory activity rather than regulatory function itself, and not all OCRs are necessarily active or functionally important regulatory elements ^15,16^.

Distinguishing functionally important regulatory elements among OCRs is particularly relevant to understanding human traits and diseases. Genome-wide association studies (GWAS) have demonstrated that disease-associated variants are enriched in non-coding regions ^17,18^, particularly in regulatory elements ^7,19,20^. Nevertheless, identifying which accessible regions are functionally important and relevant to phenotypic variation remains challenging ^15–17,21,22^. Large-scale epigenomic efforts, including ENCODE Phases 3 and 4 ^12,23^, have integrated chromatin accessibility with other epigenomic features, such as histone modifications, to annotate candidate cis-regulatory elements (cCREs). Chromatin accessibility quantitative trait loci (caQTLs) provide complementary information by identifying genetic variants associated with variation in chromatin accessibility ^24,25^. However, these approaches do not fully resolve which OCRs are critically dependent on their underlying DNA sequence for regulatory function, particularly in a cell-type-specific manner.

Evolutionary constraint provides an orthogonal source of information for assessing functional importance. Genomic regions depleted of genetic variation or sequence substitutions relative to expectation are likely to have been subject to purifying selection, reflecting functional consequences of sequence disruption ^4,26^. Recent large-scale human whole-genome sequencing studies have enabled the development of high-resolution constraint metrics such as Gnocchi ^27^, which accounts for local mutability to identify genomic regions depleted of genetic variation. Cross-species conservation metrics provide complementary evidence of evolutionary constraint over different evolutionary timescales. However, regulatory elements can be highly cell-type-specific and evolutionarily dynamic, and existing constraint metrics do not directly characterize how changes in the underlying DNA sequence affect regulatory activity in a specific cellular context. Thus, systematically identifying cell-type-specific regulatory regions whose function is sensitive to sequence perturbation remains an important challenge.

Sequence-based machine-learning models offer an opportunity to address this challenge. Models trained on genomic sequence can predict tissue- and cell-type-specific chromatin features and regulatory activity ^28–31^ and can estimate the effects of sequence variants without requiring the variants to have been observed in a population. In principle, systematically perturbing the input sequence of such models could quantify the extent to which the predicted regulatory activity of an OCR depends on its underlying DNA sequence. This sequence-to-function approach is conceptually distinct from directly measuring evolutionary constraint: rather than using observed genetic variation or cross-species conservation, it assesses the predicted functional consequences of sequence perturbation. Whether such sequence-derived regulatory sensitivity can identify functionally important OCRs under evolutionary constraint, however, remains unclear.

Here, we developed CAMBUS (Chromatin Accessibility Mutation Burden Score), a framework that systematically quantifies the predicted sensitivity of OCRs to sequence perturbation. CAMBUS uses *in silico* saturated mutagenesis to identify OCRs whose predicted chromatin accessibility is strongly dependent on their underlying DNA sequence (Fig. 1), without directly incorporating human genetic variation or existing evolutionary constraint metrics. We applied CAMBUS across 29 immune cell types and identified OCRs that were strongly enriched for independently defined evolutionary constraint and active regulatory elements. We further evaluated their relevance to complex traits through heritability analyses and statistical fine-mapping, demonstrating enrichment for putative causal variants, including rare variants, across European (EUR) and East Asian (EAS) ancestries (Fig. 1D and 1E). Together, CAMBUS provides a region-centered framework for identifying cell-type-specific regulatory elements that are sensitive to sequence disruption and enriched for evolutionary constraint, thereby facilitating the functional interpretation of non-coding genetic variation.

**Fig. 1.**
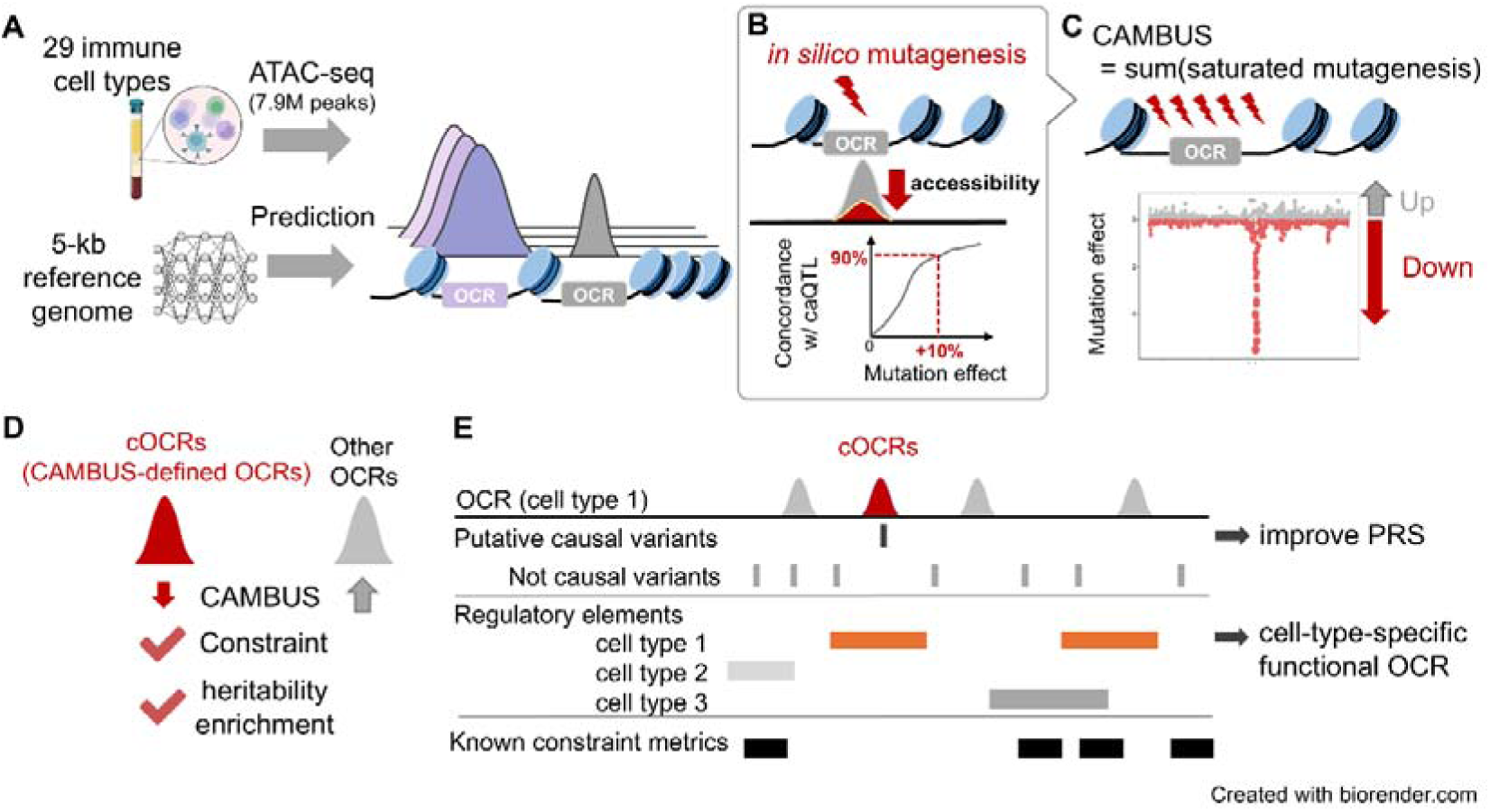
Study design. Overview of the study design. **A,** Dataset comprising 29 cell types was used to validate our approach, and a machine learning model was developed to predict OCRs. **B, C,** Identification of CAMBUS-defined OCRs (cOCRs) based on predicted sensitivity to sequence perturbation. **D,** Characteristics of cOCRs based on CAMBUS. **E,** Characterization of cOCRs and prioritization for putative causal variants.

## Results

### Comprehensive ATAC-seq profiling of 29 immune cell types

We sorted PBMCs (peripheral blood mononuclear cells) and Neutrophils from 55 Japanese individuals (Methods). PBMCs were sorted into 27 immune cell types by cell sorter. We conducted ATAC-seq for PBMC, Neutrophil, and the 27 cell types (hereafter, 29 cell types) in eight subgroups (CD4; CD4^+^ T-cells, CD8; CD8^+^ T-cells, B; B cells, Mono; Monocytes, DC; dendritic cells, NK; Natural Killer cells, Neutrophils, and PBMC) (Supplementary Table 1). The number of cell types in this study is comparable to that in recent scATAC-seq studies using PBMC ^32,33^. Notably, the 29 cell types include minor cell types, such as double-negative B-cells (∼5% in B-cells) ^34^. After stringent quality control, we obtained 7,939,714 OCRs that covered 498 Mb (16.1 %) of the human genome (Methods). The median number and size of OCRs were 252,668 OCRs and 276 bp, respectively. Among the 7,939,714 OCRs, 330,367 (4.2%) were observed in only one cell type, and 437,368 (5.5%) were observed in only one subgroup. Both hierarchical clustering and dimension reduction using principal component analysis (PCA) and uniform manifold approximation and projection (UMAP) showed that these OCRs could clearly distinguish the eight subgroups but not all 29 immune cell types (Extended Data Fig. 1).

### Identifying mutation-sensitive OCRs using CAMBUS

To quantify the sensitivity of OCRs to sequence perturbation, we first developed CAMBUS ML (Chromatin Accessibility Mutation Burden Score machine learning), a sequence-based model that predicts chromatin accessibility across each of the 29 cell types. We then performed *in silico* saturated mutagenesis to quantify the predicted effects of all possible single-nucleotide substitutions within 2 kb of each OCR, irrespective of whether these variants have been observed in human populations (Methods). For each OCR, we aggregated the predicted effects of these sequence perturbations to calculate CAMBUS (Fig. 2A, 2B; Methods). OCRs with significantly negative CAMBUS values were designated CAMBUS-defined OCRs (cOCRs), representing regions in which sequence perturbations were predicted to predominantly reduce chromatin accessibility. We hypothesized that such mutation-sensitive OCRs would be enriched for functionally important regulatory elements under evolutionary constraint. We additionally defined relaxed cOCRs as OCRs within the bottom 5% of the CAMBUS distribution. The median number of cOCRs was 1,584, covering 0.056% of the human genome, while that of relaxed cOCRs was 12,632, covering 0.33%. Notably, read counts from ATAC-seq were not a determinant factor for cOCRs (Extended Data Fig. 2A), reinforcing that our machine-learning approach incorporated information from DNA sequence patterns into OCRs, not from technical fluctuations or noises.

**Fig. 2.**
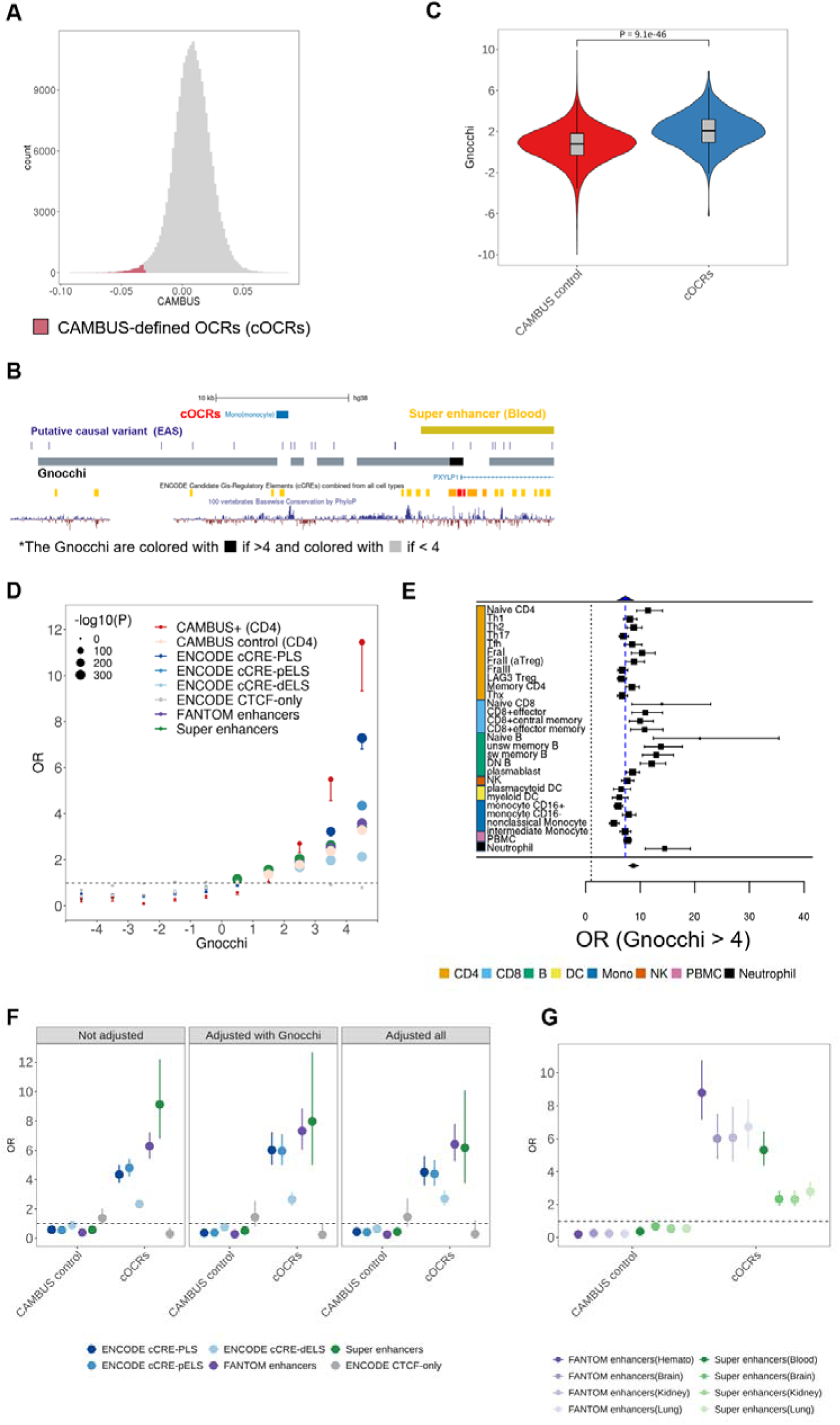
cOCRs are enriched for evolutionary constraint and active regulatory elements. **A,** Distribution of CAMBUS. The red region represents cOCRs with significantly negative CAMBUS values. **B,** Genome browser view at a representative cOCR overlapping a Gnocchi-defined constrained region in monocytes (chr3:141,217,899-141,218,844). The black box represents Gnocchi with Z > 4, and the grey box represents Gnocchi with Z < 4. **C,** Distribution of non-coding Gnocchi in cOCRs and CAMBUS control OCRs in NaiveCD4. The one-sided Wilcoxon-tests were performed between cOCRs and CAMBUS control OCRs. **D, E,** Enrichment (odds ratio; OR) of cOCRs in non-coding Gnocchi (n = 1,843,599), comparing with the known regulatory elements (using the previous study’s results, see URLs) in non-coding Gnocchi (n = 1,843,599). We calculated the OR of cOCRs (n = 1,266) and CAMBUS control OCRs (n = 251,807) for NaiveCD4 using various Gnocchi’s thresholds in (**D**) and OR for each cOCR in 29 cell types and their pooled OR through random-effects meta-analysis using Gnocchi > 4 in (**E**). The blue dashed line represents the OR of ENCODE cCRE-PLS (**E**). **F**, Enrichment of cOCRs for NaiveCD4 (n = 1,266) in the known cCREs (ENCODE cCRE-PLS; n = 40,891, ENCODE cCRE-pELS; n = 172,027, ENCODE cCRE-dELS; n = 789,200, ENCODE CTCF-only; n = 35,839, FANTOM5 enhancers; n = 63,280, Super enhancers; n = 16,472), adjusted for seven constraint metrics (Gnocchi, phyloP46way, phyloP100way, cactus241way, phyloPPrimates, GERP and DR), GC content, and TF motif density. **G,** Enrichment of cOCRs for NaiveCD4 (n = 1,266) in the tissue expressed FANTOM5 enhancers (hematopoietic cells; n = 64,225, Brain; n = 62,978, Kidney; n = 35,177, Lung; n = 26,630), and Super enhancers (Blood; n = 8,055, Brain; n = 9,372, Kidney; n = 4,712, Lung; n = 9,878), adjusted for seven constraint metrics (Gnocchi, phyloP46way, phyloP100way, cactus241way, phyloPPrimates, GERP and DR), GC content, and TF motif density. The dashed line represents no enrichment (OR=1.0; **D, E, F, G**). CAMBUS control in NaiveCD4 (n = 251,807) is used as background in (**F, G**). The error bars represent 95% confidence intervals, with only lower bounds shown in **D** (See Supplementary Table 4 for upper bounds).

We next tested whether the mutational sensitivity captured by CAMBUS was associated with independently defined evolutionary constraint (Methods). As expected, five out of the seven constraint metrics supported the constraint hypothesis for cOCRs across 29 cell types (Methods; Supplementary Table 3). Notably, we found that Gnocchi scores were significantly higher in cOCRs than in CAMBUS control OCRs (one-sided Wilcoxon-test P = 2.5 × 10^-69^ for cOCRs in NaiveCD4). Although cOCRs contained a higher proportion of coding regions than control OCRs (Extended Data Fig. 2B), the same trend of enrichment was observed even when the analysis was restricted to non-coding regions (one-sided Wilcoxon-test P = 9.1 × 10^-46^ for cOCRs in NaiveCD4; Fig. 2C; Extended Data Fig. 2C). Of note, 59,605 (90%) of cOCRs were not prioritized according to the prespecified thresholds of the seven existing constraint metrics (Methods). Nevertheless, the cOCRs showed the strongest constraint defined by Gnocchi (Gnocchi Z≥ 4; odds ratio (OR)=11.45 (95% confidence interval (CI) 9.33–14.05), P=4.7 × 10^-68^ by Fisher’s Exact test) among the compared regulatory elements (Fig. 2D; Supplementary Table 4). Meta-analysis demonstrated the consistent tendency across the 29 cell types (OR=8.71 (95% CI 7.85–9.66), P=3.1 × 10^-365^ from random-effects meta-analysis; Fig. 2E), which was higher than that reported for ENCODE promoter-like regulatory elements (OR = 7.3), the candidate regulatory element category showing the strongest constraint in the previous study ^27^. The relaxed cOCRs also showed the same patterns with slightly reduced magnitude (Extended Data Fig. 2D). Besides, we observed the enrichment of cOCRs in human ancestor quickly evolved regions (HAQER) ^35^ (OR=2.35 (95% CI 1.91–2.89), P=8.3 × 10^-16^). Taken together, cOCRs were strongly enriched for independently defined evolutionary constraint while capturing numerous regions not prioritized by existing constraint metrics.

### cOCRs are enriched for active regulatory elements

As expected, the OCRs obtained in this study were well overlapped with the known candidate regulatory elements; for example, 73.66% of all OCRs in NaiveCD4 were overlapped with super-enhancers ^36^ (the median of 29 cell types is 74.70%) and 4.49% with FANTOM5 enhancers ^6^ (the median of 29 cell types is 5.10%; Supplementary Table 7). Then we explored whether cOCRs could aid in prioritizing functional regulatory elements in comparison with control OCRs (Methods). As a result, cOCRs significantly enriched for the known regulatory elements, especially active enhancers, like super-enhancers (NaiveCD4: OR=9.07 (95% CI 6.80–12.11), Fisher’s exact test P=2.0 × 10^-102^) and FANTOM5 enhancers (NaiveCD4: OR=6.16 (95% CI 5.39–7.03), Fisher’s exact test P=6.3 × 10^-111^) (Supplementary Table 8). Meta-analysis showed a consistent trend across 29 cell types (OR=6.63 (95% CI 5.50–7.99), P = 1.1 × 10^-87^ from random-effects meta-analysis for super-enhancers; Extended Data Fig. 3A, and OR=5.19 (95% CI 4.87–5.54), P=4.9 × 10^-547^ from random-effects meta-analysis for FANTOM5 enhancers; Extended Data Fig. 3B). The cOCRs exhibited higher GC content and Transcription Factor (TF) motif density, consistent with their enrichment in regulatory sequences (one-sided Wilcoxon-paired test P=9.2 × 10^-6^, 2.5 × 10^-4^ respectively; Extended Data Fig. 3C). Remarkably, even after further adjusting for these sequence features in addition to the known seven constraint metrics, we still observed substantial enrichment for regulatory elements in cOCRs across the 29 cell types (e.g., super-enhancers in NaiveCD4: OR=6.18 (95% CI 3.80-10.05), P=2.1 × 10^-13^; Fig. 2F; Supplementary Table 9). The same trend of enrichment was observed even when the analysis was restricted to non-coding regions across the 29 cell types (e.g., super-enhancers in NaiveCD4: OR=10.67 (95% CI 5.02-22.66), P=7.3 × 10^-10^; Extended Data Fig. 3D; Supplementary Table 9). Reflecting the characteristics of the analyzed cell types (blood and immune cell types), a comparison of enrichment across four tissues (blood/hematopoietic, brain, kidney, and lung) revealed the strongest prioritization of blood’s regulatory elements using logistic regression models with the same adjustments (CD4 as a representative: super-enhancers; OR=5.32 (95% CI 4.42–6.41), P=1.3 × 10^-69^; FANTOM5 enhancers; OR=8.80 (95% CI 7.22–10.72), P=5.9 × 10^-103^; Fig. 2G; Supplementary Table 10). This pattern was consistently observed across 29 cell types (Supplementary Table 11). We also confirmed that the relaxed cOCRs exhibited the same tendency (Extended Data Fig. 3E; Supplementary Table 12).

A previous study ^37^ demonstrated tissue-specific enrichment of archaic introgressed variants in enhancers. Consistent with the evolutionary relevance of regulatory elements, cOCRs were also enriched for archaic introgressed sequences ^38,39^ (OR=1.43 (95% CI 1.35–1.51), P=4.8 × 10^-32^ from random-effects meta-analysis; Extended Data Fig. 3F and Supplementary Table 5). Together, these results demonstrate that cOCRs are strongly enriched for active regulatory elements, particularly tissue-relevant enhancers, even after accounting for established constraint metrics and sequence features.

### cOCRs are enriched for the heritability of immune-mediated traits

Given the established contribution of regulatory variation to complex-trait heritability, we next tested whether cOCRs captured trait-associated genetic variation beyond that explained by broader OCR annotation. We computed heritability enrichments in cOCRs using stratified LD score regression (S-LDSC) ^40^ conditioning on known functional annotations and all OCRs from this study (Methods). We used GWAS summary statistics for 33 complex traits, including 14 immune-mediated traits (e.g., multiple sclerosis (MS), rheumatoid arthritis (RA), and Atopic dermatitis (AD)), from European (EUR) and East Asian (EAS) populations (Supplementary Table 13).

As a result, cOCRs demonstrated substantial heritability enrichment in a wide variety of immune-mediated diseases (FDR < 5%; Fig. 3A; Supplementary Table 14). Particularly, we found plausible pairs of immune traits and cell types supported by the strong enrichment of heritability: MS and T_h_17 ^41^, and systemic lupus erythematosus (SLE) and B cells ^42,43^ (Extended Data Fig. 4A). Other plausible pairs included pairs of RA with CD4 or CD8^+^ T cells, and AD with CD4 (particularly Th2 ^44^), or CD8^+^ T cells. Remarkably, comparable heritability enrichments were not observed for the other constraint annotations matched for annotation size (Fig. 3B).

**Fig. 3.**
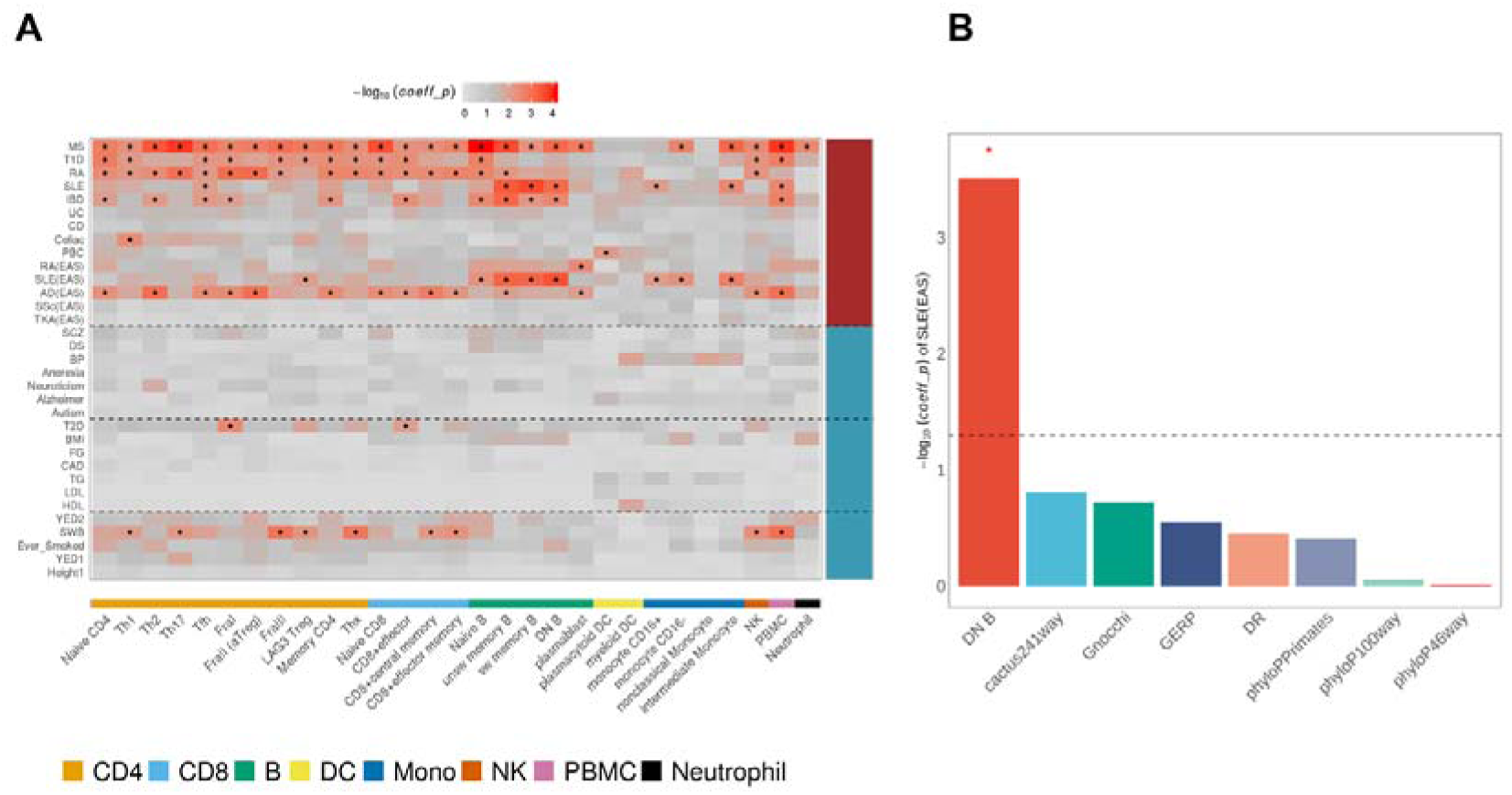
Heritability enrichment for immune-mediated diseases. **A,** The heritability enrichment of various GWAS traits (in total, 33 GWAS for EUR and EAS ancestries; Supplementary Table 13) in cOCRs for each of 29 cell types. The heatmap colors indicate the -log10-scale p-value of the coefficient for cOCRs annotation in S-LDSC (coefficient P). **B,** The -log10-scale coefficient P of cOCRs in double-negative B-cells (DN B) and seven known constraint metrics for SLE GWAS in EAS ancestry. ‘*’ indicates significance (qvalue < 0.05) after FDR correction (**A, B**).

Based on these results, we hypothesized that polygenic risk scores (PRS) prioritized by cOCRs could enhance their predictive ability. To evaluate this hypothesis, we focused on cOCRs in double-negative B cells (DN B), which showed significant enrichment for the heritability of SLE in EAS ancestry (Extended Data Fig. 4A) and were also known for their functional associations with SLE ^45^. We constructed the PRS based on variants overlapping cOCRs and evaluated the additional performance of the conventional PRS by five-fold cross-validation using 137,620 individuals (Methods). A fixed-effect meta-analysis demonstrated that the cOCR-prioritized variants provided significant predictive information beyond the conventional PRS (P=3.12 × 10^-6^; Supplementary Table 15; Methods). These suggest that cOCRs can help to characterize the polygenic and cell-type-specific genetic architecture of complex traits and may provide complementary information for genetic risk prediction by prioritizing cell-type-specific regulatory regions relevant to the traits.

### Putative causal variants were enriched in cOCRs

To further assess whether the region-level prioritization by cOCRs could inform causal variant identification, we examined whether variants within cOCRs were preferentially enriched among known putative causal variants for complex traits. As expected, the median regional length of cOCRs (845 bp) was much shorter than that of 95% credible sets (3.0M bp for EUR ancestry and 1.1M bp for EAS ancestry), which may facilitate the prioritization of putative causal variants. We observed cOCRs showed significant enrichment in the pathogenic regulatory variants cataloged in the Human Gene Mutation Database (HGMD) ^46^ (OR=2.70 (95% CI 2.43–2.99), P=1.7 × 10^-77^; from random-effects meta-analysis; Extended Data Fig. 3G; Supplementary Table 6). This is in line with overlap between cOCRs and super-enhancers or FANTOM5 enhancers, both of which are known to be enriched for putative causal variants ^47^. We further analyzed the large-scale statistical fine-mapping studies in 94 traits in EUR ancestry ^48^ and 62 traits in EAS ancestry ^49^. The cOCRs were significantly enriched for the putative causal variants in 95% credible sets (median OR:3.85 and 3.94 in EUR and EAS, respectively, Extended Data Fig. 5A). In line with a data source of OCRs, cOCRs exhibited significantly higher enrichment for the putative causal variants of hematopoietic traits, especially leukocyte-related traits (Methods, Supplementary Table 21) than non-hematopoietic traits (two-sided *t*-test P=1.0 × 10^-11^ in EUR ancestry and 1.4 × 10^-2^ in EAS ancestry for leukocyte-related traits; Extended Data Fig. 5B). As an example, cOCRs in type 1 helper T-cells demonstrated stronger enrichment for variants with high posterior inclusion probability (PIP) (>0.9) for hematopoietic traits in EAS ancestry (OR=7.62 (95% CI 3.60–16.18), P=1.2 × 10^-7^ from logistic regression; Fig. 4A), and this enrichment was further increased when restricted to the leukocyte-related traits (OR=11.0 (95% CI 4.04–29.96), P=2.7 × 10^-6^; Fig. 4A). Meta-analysis showed that this trend of leukocyte-related traits was consistent across 29 cell types (OR=8.99 (95% CI 7.85–10.29), P=8.8 × 10^-221^ for EUR ancestry; OR=19.24 (95% CI 16.03 – 23.09), P=3.9 × 10^-221^ for EAS ancestry; Fig. 4B). Taken together, these results indicate that cOCRs preferentially overlap fine-mapped putative causal variants and may therefore facilitate their functional prioritization.

**Fig. 4.**
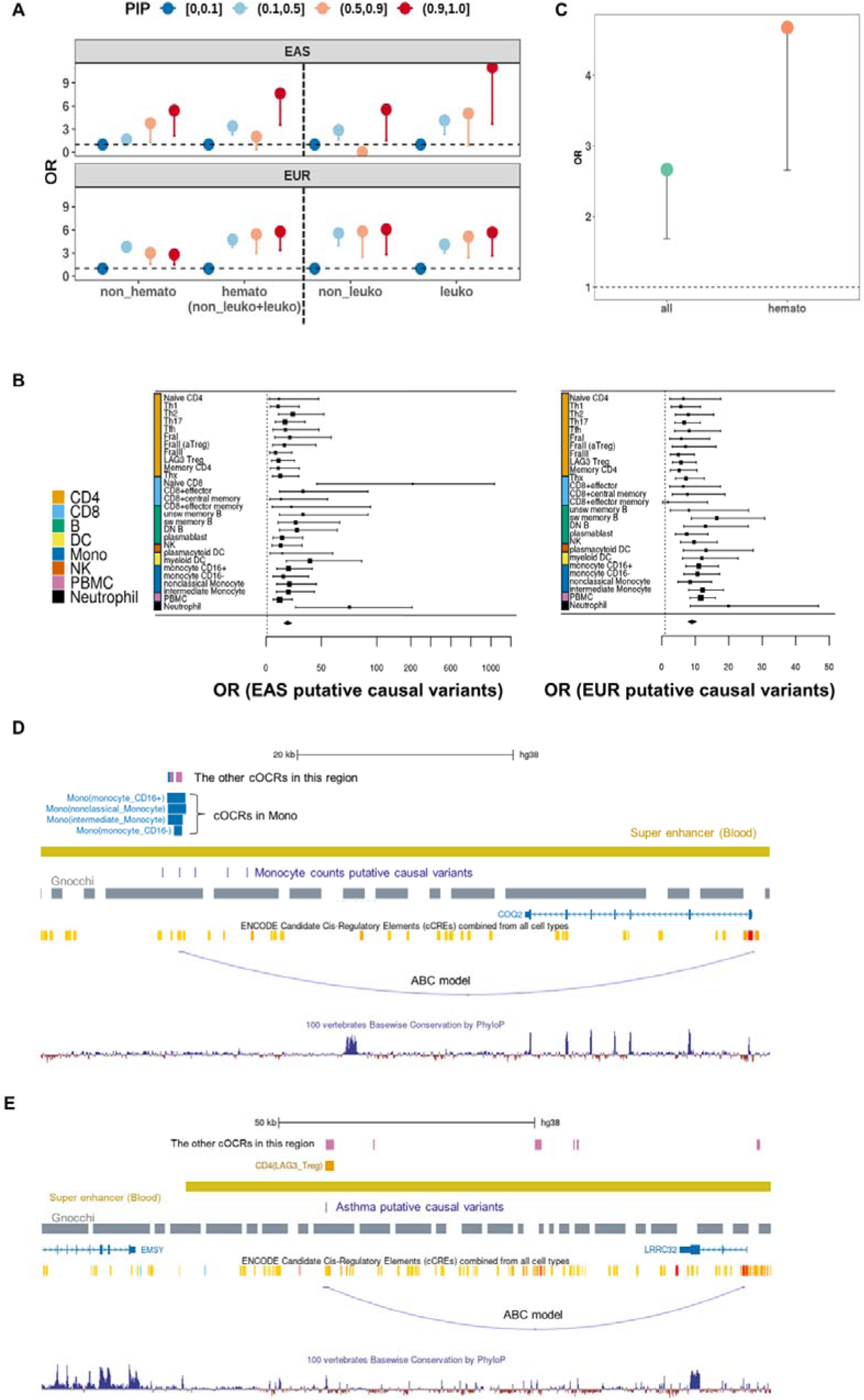
Prioritization of putative causal variants using cOCRs. **A,** Enrichment of putative causal variants in leukocyte (leuko), non-leukocytes (non_leuko), hemato (leuko + non_leuko), and non-hematopoietic (non_hemato) traits in cOCRs for Th1. Putative causal variants were obtained from statistical fine-mapping analysis of many complex traits’ GWAS results for EAS and EUR populations ^48,49^. **B,** Forest plot showing the enrichment of cOCRs across 29 cell types and the pooled OR in highly confident putative causal variants (PIP > 0.9). Variants with PIP <= 0.1 were used as the background for OR calculation. **C,** Enrichment of cOCRs for any of 29 cell types in putative causal rare variants (maf < 0.01 in EAS), evaluated by comparing the proportion of cOCRs overlapping with putative causal rare variants with PIP > 0.1. The left dot represents the OR calculated using all the putative causal variants, while the right dot represents the OR based only on the putative causal variants for hematopoietic phenotypes. The dashed line represents no enrichment (OR=1.0). **D,E,** Representative examples of variants associated with monocyte counts prioritized with cOCRs in Mono (**D**) and asthma prioritized with cOCRs in regulatory T-cells (**E**) in putative causal variants in EUR ancestry and surrounding genomic annotations. The navy boxes represent putative causal variants in this credible set. The blue dashed line represents an enhancer-gene connection to the nearest protein-coding gene predicted by the ABC model for CD14^+^ monocytes (**D**) and HUVEC (**E**). The gene with the highest ABC score is also shown in (**D**). All the other cOCRs around putative causal variants (rs13122305 (**D**), rs55646091 (**E**)) are also shown in the figures (The purple box represents PBMC).

Notably, even for rare variants (minor allele frequency (MAF) < 0.01), the putative causal variants of hematopoietic traits were significantly enriched in cOCRs (Methods; OR=4.68 (95% CI 2.66–7.99), two-sided Fisher’s exact test P=2.2 × 10^-6^; Fig. 4C). This suggested that cOCRs can potentially facilitate the prioritization of rare putative causal variants that are difficult to resolve through conventional statistical fine-mapping alone ^22^.

We found plausible examples of putative causal variants that were not covered by known constraint regions (Gnocchi) but were identified by cOCRs, providing plausible biological interpretations of GWAS signals. First, rs13122305 was the only variant overlapping cOCRs in monocytes among seven candidate causal variants in a credible set for monocyte count (Fig. 4D). We found that the Activity-by-Contact (ABC) model’s prediction (ABC enhancer) in monocytes only overlapped with rs13122305 and that *COQ2*, whose anti-inflammatory functions via regulating gene expression were reported ^50,51^, was supported by the ABC model with the second-highest ABC score (0.025) among the eight target genes. Second, among the three candidate causal variants in a credible set in the *FCGR* region for Albumin/Globulin ratio (AG), only rs2487451 overlapped with cOCRs. Non-classical monocytes, known to display antigen presenting properties ^52^, were the only cell type whose OCR overlapped with the candidate causal variant. The nearest gene was *FCGR3B*, encoding a receptor (FcγRIIIB) that binds to the Fc region of IgG (Immunoglobulin) antibodies, which is known to be expressed in non-classical monocytes ^53^. Third, rs55646091, a putative causal variant for asthma (PIP = 0.99), overlapped with cOCRs of regulatory T-cells (T-reg) (Fig. 4E). The nearest gene of this variant was *LRRC32* (leucine-rich repeat-containing 32), which regulates TGF-beta. *LRRC32* was potentially involved in asthma ^54^ and its encoding protein was known as a surface marker for T-reg. Furthermore, this variant-gene pair was supported by the ABC model and *LRRC32* displayed the highest ABC score (ABC score = 0.041 that is much higher than the threshold corresponding to 70% recall and 59% precision) ^55^ among the nine genes connected with the regulatory element overlapping with the OCR and the variant. Fourth, we found that rs8009224 was the only one that overlapped with cOCRs of plasmablast (Extended Data Fig. 6A) among the seven putative causal variants for hemoglobin A1c levels. This is an intronic variant of *MIDEAS* (Mitotic Deacetylase Associated SANT Domain Protein), which is low frequency in both 1KG EUR (MAF = 3.4%) and EAS (MAF = 1.2%) and also associated with mean corpuscular hemoglobin concentration ^56^. The overlapping regulatory element was connected to the 16 genes by the ABC enhancer in B-cell, and the putative target gene with the second highest ABC score is *MIDEAS* (ABC score = 0.097, much higher than the proposed threshold corresponding to 70% recall and 59% precision). Lastly, we demonstrated that cOCRs in T-reg could prioritize only one putative causal variant (rs9814991) from numerous variants (fifty-seven) in the credible set for eosinophil count (Extended Data Fig. 6B). These indicate that cOCRs, together with statistical fine-mapping, enable the prioritization of regulatory regions likely to harbor putative causal variants of complex traits in a cell type-specific manner.

## Discussion

In this study, we developed CAMBUS to quantify the extent to which predicted chromatin accessibility depends on the underlying DNA sequence. Importantly, cOCRs were defined solely on the basis of predicted sensitivity to sequence perturbation, without directly incorporating evolutionary constraint metrics or human genetic variation. Nevertheless, cOCRs were strongly enriched for independently defined evolutionary constraint and active regulatory elements, while also capturing high-resolution, cell-type-specific regulatory regions not readily prioritized by existing regulatory or constraint annotations. These findings suggest that sequence sensitivity of regulatory activity provides information complementary to conventional measures of evolutionary constraint.

Sequence-to-function models have shown that DNA sequence patterns can predict chromatin features and estimate the effects of non-coding variants on regulatory outputs ^31^. MPRA-based saturation mutagenesis of mammalian enhancers has further shown that activity-reducing mutations are depleted during evolution, consistent with functional constraint on sequence features required for regulatory activity ^57^. Population-genetic analyses have similarly demonstrated signatures of purifying selection in functional genomic elements, including cis-regulatory elements ^58–60^. Together, these observations provide a mechanistic basis for the relationship observed here between sequence-dependent regulatory sensitivity and evolutionary constraint.

An alternative potential approach using ATAC-seq (or DNase-seq) could prioritize regions with high accessibility (i.e., regions with many read counts). However, previous studies have suggested no relationship between accessibility and functionality or constraints ^61–63^. Consistently, accessibility was not a determinant factor in defining cOCRs. Thus, leveraging surrounding DNA sequence patterns through machine learning offers a unique and effective strategy for identifying biologically important OCRs.

Established constraint metrics differed in their ability to capture the evolutionary constraint observed in cOCRs (Supplementary Table 3), with GERP and phyloP100way showing no significant difference between cOCRs and CAMBUS control OCRs. These differences likely reflect, at least in part, differences in the species included, evolutionary timescales, alignment strategies, and statistical definitions of constraint across metrics. phyloP100way is derived from alignments between humans and a broad range of vertebrate species, many of which are phylogenetically distant from humans. Therefore, these metrics may have limited sensitivity to lineage-specific or rapidly evolving regulatory constraints in humans ^64^, such as immune-cell-specific enhancers ^65^. In contrast, human-variation-based metrics and primate- or mammalian-scale conservation scores (Gnocchi, DR, phyloPPrimates, phyloP46way, cactus241way) may be more closely aligned with the regulatory constraint captured by cOCRs. Thus, these metric-dependent differences suggest that cOCRs capture regulatory elements enriched for evolutionary constraint that is not fully represented by conventional cross-species conservation metrics ^5^. cOCRs showed enrichment for active and functional regulatory elements, particularly tissue-specific active enhancers, even after adjustment for established constraint metrics (Gnocchi, phyloP46way, phyloP100way, cactus241way, phyloPPrimates, GERP, and DR) and sequence features (GC content, TF motif density). Rather than replacing existing evolutionary metrics, cOCRs provide complementary information for identifying functionally constrained regulatory elements. Beyond their enrichment for evolutionary constraint and active regulatory elements, cOCRs captured significant heritability for multiple complex traits and preferentially overlapped fine-mapped putative causal variants, particularly for immune-related traits. These findings demonstrate the potential of cOCRs to connect trait-associated genetic variation with cell-type-specific regulatory mechanisms.

Notably, cOCRs also effectively prioritize rare putative causal variants for complex traits. Previous population-scale sequencing studies showed that rare-variant excess is greater at evolutionarily conserved or constrained sites ^66,67^, consistent with the enrichment of constrained regulatory regions among cOCRs. In GWAS using SNP arrays, low statistical power and low imputation accuracy for rare variants make it difficult to detect rare variant associations and identify rare putative causal variants through statistical fine-mapping. Despite these challenges, cOCRs can facilitate the rare variant prioritization that might otherwise be missed.

Eosinophilic asthma is a subtype characterized by elevated levels of eosinophils ^68^. In this analysis, we identified a variant (rs55646091) associated with asthma linked to increased eosinophil counts (PIP = 0.03). This variant exhibited positive effect sizes in both traits (0.20 for asthma, 0.072 for eosinophil counts), suggesting that carriers of this variant demonstrate elevated eosinophil levels, which contribute to inflammation and the onset of asthma ^69^. Notably, this variant overlaps specifically with the OCR in T-reg cells among 29 blood cell types analyzed. The nearest gene of this variant is *LRRC32*, which regulates TGF-beta, a known surface marker of T-reg cells, which has been previously implicated in immune regulation ^70^. Together, these observations suggest a plausible regulatory mechanism linking rs55646091, T-reg cell function, eosinophilic inflammation, and asthma susceptibility. Although these data do not establish a causal pathway, they illustrate how integration of cOCRs with fine-mapping can generate cell-type-specific hypotheses for the biological interpretation of GWAS signals.

In this study, we focused on various immune cell types; however, our approach is broadly applicable to other tissues and cell types. CAMBUS is designed to characterize regulatory constraints at the level of OCRs rather than to predict those of individual variants. To achieve this, CAMBUS systemically performs *in silico* saturated mutagenesis surrounding each OCR and summarizes their predicted effects at the region level. Computational efficiency is therefore an important consideration for applying CAMBUS across millions of regulatory regions. Our relatively simple CNN-based model achieved predictive performance comparable to that of the more recent Sei- ^71^ and ChromBPNet-based models ^72^, while enabling substantially faster *in silico* mutagenesis, providing a practical balance between predictive performance and computational efficiency for large-scale CAMBUS calculation (Supplementary Note). By contrast, substantially large sequence models would be far more computationally demanding; for example, applying Evo 2 ^73^ to the same task is estimated to require more than 200 years on a single H100 GPU (Supplementary Note).

Recent work has developed conceptually related frameworks that integrate predicted regulatory effects with evolutionary conservation to prioritize individual non-coding variants, such as FLARE ^74^. In contrast, CAMBUS addresses a distinct but complementary problem. Rather than prioritizing individual variants, CAMBUS uses *in silico* saturated mutagenesis to identify regulatory OCRs whose regulatory activity is strongly dependent on their underlying DNA sequence, without relying on known variants or existing constraint metrics as training targets. This region-centered framework enables systematic prioritization of functionally important regulatory elements, including evolutionarily constrained regions that are not captured by established constraint metrics. As GWAS results continue to accumulate, integrating cOCRs with these expanding resources has the potential to uncover further insights into the regulatory mechanisms underlying complex traits. Moreover, this method can be applicable to OCRs identified at the single-cell level, further extending its applicability.

Taken together, CAMBUS provides a computationally efficient framework for identifying cell-type-specific regulatory regions that are sensitive to sequence perturbation and enriched for evolutionary constraint. By prioritizing functionally important regulatory elements rather than individual variants, cOCRs provide complementary information for interpreting non-coding genetic variants and understanding the regulatory architecture of complex traits.

## Methods

### Sample and library preparation

We collected peripheral blood (PB) cells from 55 healthy Japanese volunteers and sorted them into 29 cell types. PBMCs were isolated by density gradient separation with Ficoll-Paque (GE healthcare) immediately after blood drawing. Erythrocytes were lysed with a potassium ammonium chloride buffer, and non-specific binding was blocked with Fc-gamma receptor antibodies. PBMCs were stained with cell surface specific antibodies (Supplementary Table 22). 27 immune cell populations were sorted by FACS (BDAria III) using the gating strategy (Supplementary Table 1; Extended Data Fig. 7). In addition, neutrophils were isolated by MACSxpress Neutrophil Isolation Kits human (Miltenyi Biotec) from 4 × 10^6^ whole blood cells. Sorted cells, isolated PBMC, and Neutrophils were stored at −80°C in CytoStor-10 (Charles River Laboratories Cell Solutions). Libraries for ATAC-seq were prepared using a modified protocol of Omni-ATAC ^75^. Genomic DNA from whole blood was isolated using QIAamp DNA Blood Mini Kits, respectively. ATAC-seq was performed using 150-bp paired-end sequencing on Illumina NovaSeq6000. In total, we obtained a median of 13.34M (+/- 7.56M) reads per sample. This study was approved by the Ethics Committees of RIKEN. Written informed consent was obtained from each volunteer.

### Mapping ATAC-seq data

ATAC-seq data were processed following the ENCODE ATAC-seq pipeline (version 1.8.0) as follows. Raw reads were trimmed using cutadapt (version 2.5) and aligned to the human genome assembly GRCh38 analysis set using bowtie2 (version 2.3.4.3) for each sample. Multimapping reads were processed according to the ENCODE pipeline, and aligned reads were filtered using samtools (version 1.9) to retain properly paired primary alignments. Duplicate reads were identified using MarkDuplicates (Picard version 2.20.7-SNAPSHOT) and subsequently removed. Reads mapped to the mitochondrial genome were also excluded. After quality control, a median of 5.39 million reads per sample (s.d., 3.28 million) were retained.

### ATAC-seq peak calling

For each of the 29 immune cell types, quality-controlled BAM files from individual samples were merged using samtools merge. ATAC-seq peaks, representing OCRs, were then identified using MACS2 (version 2.1.1) following a previously described approach ^25^, with the parameters (-p 0.01 --shift -75 --extsize 150 --nomodel). Across the 29 cell types, we obtained a median of 252,668 (s.d., 66,703) OCRs with a median peak width of 276bp (s.d., 359 bp) (FDR<0.05). Using the alignment file for each cell type, we calculated the read counts from the first mates for each OCR with bedtools (version 2.30.0)’s coverage function.

### Quality control for ATAC-seq dataset

We first assessed the quality of ATAC-seq data based on Pearson correlations of chromatin accessibility between samples within each cell type. For this analysis, we used a common set of 614,516 peaks identified across all samples using the same peak-calling parameters described above; these peaks were used exclusively for quality control. Samples with Pearson correlations of ≤0.65 were excluded, with the threshold determined by manual inspection of the correlation distributions. Additional outlier samples showing low correlations with other samples of the same cell type were excluded by visual inspection (Supplementary Table 16). Samples with a mapping rate of <50% were also excluded. After quality control, 1,386 samples were retained, with a median of 48 donors per cell type (Supplementary Table 1).

### Hierarchical clustering and dimension reduction

We first selected the 5,000 most accessible OCRs from each cell type based on TPM values and merged the selected OCRs across all 29 cell types, yielding 145,000 OCRs. We then constructed a binary matrix (0, 1) indicating whether each 300-bp genomic bin overlapped an OCR (1) or not (0) in each cell type. Hierarchical clustering was performed on this binary matrix using Ward’s method with Euclidean distance. PCA was performed on the binary matrix using the binaryPCA function in the scBFA package (version 1.16.0) ^76^ in R with default parameters. UMAP was performed using the top five principal components with the umap function from the scikit-learn package (version 0.18.1) in Python with default parameters.

### Strategy for the CAMBUS ML model

The CAMBUS ML model was based on the DeepSEA architecture, a deep CNN model ^31^. We used the Selene library (v0.4.7) to train the CAMBUS ML model. The model takes a 5-kb reference sequence as input, which is five times longer than the 1-kb sequence used in the original DeepSEA model, to capture broader genomic sequence contexts. For each cell type, the model outputs a sigmoid-transformed score representing the predicted probability that the central 200-bp region of the input sequence overlaps an OCR (see Supplementary Table 17 for the configuration details). We referred to this predicted probability for chromatin accessibility as the accessibility probability.

We merged 7,939,714 OCRs across 29 cell types using the ‘bedtools merge’ function, yielding 1,349,934 non-overlapping unique input sequences. According to Selene’s default settings, we used chromosomes 8 and 9 for testing, chromosomes 6 and 7 for validation, and the other autosomal chromosomes for training data. The details of Selene’s parameters are described in Supplementary Table 17.

### *In silico* saturated mutagenesis using the trained CAMBUS ML model

For all nucleotide positions within 1kb of the center of each OCR, we used the trained CAMBUS ML model to predict the effects for all possible single-nucleotide substitutions. This comprehensive approach, inspired by the previous study of transcript expression levels ^28^, provides an unbiased analysis of mutational effects across the entire sequence, rather than focusing on only specific known polymorphisms. To quantify the mutation effect, we calculated the difference in log fold change scores between the reference and alternative alleles (putting reference alleles as denominators) for each site and averaged them across all bases. In total, we obtained 505 million mutation effects (median across the 29 cell types).

### Identification of CAMBUS-defined OCRs

We modeled the distribution of the mean predicted mutation effects (*in silico* saturated mutagenesis) within 2 kb of each OCR (CAMBUS) using a Gaussian mixture model. We defined OCRs with significantly negative CAMBUS values as CAMBUS-defined OCRs (cOCRs), representing regions in which sequence perturbations were predicted to predominantly reduce chromatin accessibility, whereas OCRs with neither significantly negative nor positive CAMBUS values were used as CAMBUS control OCRs. Local FDR was calculated using the locfdr function in the locfdr R package (version 1.1-8) for each cell type. This analysis identified a median of 1,584 cOCRs across the 29 cell types at a local FDR probability < 0.5, the defined threshold in the previous study ^28^, together with a median of 248,749 CAMBUS control OCRs. We additionally defined relaxed cOCRs as OCRs within the bottom 5% of the CAMBUS distribution for each cell type (Supplementary Table 19).

### Enrichment of cOCRs in constraint regions

We evaluated whether cOCRs are enriched in constraint regions of the genome using Gnocchi. We used the 1,984,900 QCed Gnocchi (see URLs) and calculated the OR following the methodology described in the previous paper ^27^ for 1,843,559 non-coding 1-kb windows. In brief, Gnocchi windows were stratified into the following score bins: (-∞, -4), [−4,−3), [−3,−2), [−2,−1), [−1,−0), [0,1), [1,2), [2,3), [3,4), [4,∞). For each bin, we counted the number of 1-kb windows overlapping cOCRs and calculated ORs relative to all 1-kb windows. ORs were calculated by comparing these overlaps with all 1kb windows. Fisher’s exact test was conducted to estimate the statistical significance (p-value). The OR of the known regulatory elements for each bin was obtained from the gnomAD download site (see URLs) and used for the comparison.

To assess the overlap between cOCRs and existing constraint metrics, we examined seven constraint metrics (see URLs): (1) The Gnocchi, estimated by comparing observed allele counts in diverse human populations with the estimated counts in each 1kb region of the human genome ^27^, (2) phyloP, defined by phylogenetic P-values (phyloP) scores derived from conservation across various species: phyloP46way ^77^, phyloP100way, cactus241way ^78^, based on alignments of 46, 100, and 241 species, respectively, (3) Primate-specific PhastCons (referred to as phyloPPrimates in the source dataset), defined by phylogenetic hidden Markov model (phylo-HMM) based on an alignment of 43 primate genomes ^78^, (4) Depletion Rank (DR), estimated by comparing observed allele counts in EUR populations with the estimated counts in each 500bp region of the human genome ^79^, (5) Genomic Evolutionary Rate Profiling (GERP), defined by rejected substitution scores (RS) that quantify evolutionary constraint using alignments of diverse mammalian genomes ^80^. The phyloP46way coordinates were converted to hg38/GRCh38 using liftOver as described previously. For Gnocchi and DR, the overlapping regions were considered to be overlapped if they had more than 50% overlap with each other.

To determine whether cOCRs were prioritized by existing constraint metrics, we employed the following approach: First, we calculated the mean values for each of the seven constraint metrics overlapping for each cOCR. Second, cOCR were considered not prioritized by existing constraint metrics if all seven mean values failed to meet their recommended thresholds; Gnocchi > 4 ^27^, phyloP100way with score > 3.0075 (FDR5%, following the previous approach ^81^), phyloPPrimates with score > 0.961 ^81^, cactus241way > 2.27 ^81^, DR < 0.01 ^79^, GERP > 2 (a RS score threshold of 2 provides high sensitivity while still strongly enriching for truly constrained sites, as described in the UCSC Genome Browser documentation at 2025/7/22) ^82^). For phyloP46way, we used the top 10% score (0.533) as a threshold since even the maximum score (0.655) did not reach nominal statistical significance (P = 0.221), reflecting the limited statistical power associated with the relatively small number of species included in the alignment ^77^.

### Calculation of GC content and TF motif density in cOCRs

For each OCR, GC content and TF motif density were calculated based on the hg38 reference genome. GC content was estimated using addGCBias in the chromVAR package (v.1.24.0) as the fraction of G and C bases in the reference sequence corresponding to each OCR. TF motif density was calculated using the motifmatchr package (v.1.24.0) with human JASPAR CORE motifs obtained from JASPAR2022. Motif matches were evaluated for each OCR, and TF motif density was defined as the number of distinct matched TF motifs per kilobase.

### Enrichment of cOCRs in the archaic genome-derived sequences

To assess the enrichment of cOCRs in HAQERs ^35^, we first merged cOCRs across the 29 cell types to generate a non-redundant set of cOCRs (merged cOCRs). We then determined whether each merged cOCR overlapped a HAQER and estimated the OR relative to all OCRs using logistic regression.

We also evaluated whether cOCRs are enriched in the archaic genome. For each cOCR, we determined its overlap with Neanderthal-derived sequences observed in the Japanese population ^38^ and with the Neanderthal- and Denisovan-derived sequences observed in 1000 Genomes populations ^39^. ORs were estimated relative to all OCRs using logistic regression.

### Enrichment of cOCRs in regulatory elements

We evaluated the functional enrichment of cOCRs in known regulatory elements, including ENCODE cCREs, FANTOM5 enhancers, and super-enhancers (see URLs). For each cOCR, we counted overlapping annotations for each known regulatory element and calculated the OR relative to CAMBUS control OCRs, and tested it using Fisher’s exact test, as described in the previous section.

For tissue-expressed FANTOM5 enhancers, we obtained count per million (CPM) data for 249,599 enhancers across 347 tissue/cell types in hg19 coordinates (see URLs) and selected enhancers expressed in hematopoietic_cell (N = 163), brain (N = 85), kidney (N = 16), and lung (N = 8) based on a threshold of CPM ≥ 0.1 in at least 10% of the corresponding samples. As described previously, the positions were mapped to the hg38/GRCh38 coordinates using liftOver.

For tissue-expressed super-enhancers, we obtained super-enhancer annotations for blood (N = 115), brain (N = 203), kidney (N = 34), and lung (N = 119) enhancers in hg38 coordinates (see URLs) and merged these enhancers by tissue with ’bedtools merge’ function.

To account for potential confounding by evolutionary constraint and sequence characteristics, adjusted ORs were estimated using logistic regression. The models included the mean values of seven constraint metrics overlapping each cOCR or CAMBUS control OCR (Gnocchi, phyloP46way, phyloP100way, cactus241way, phyloPPrimates, GERP, and DR), together with GC content and TF motif density, as covariates.

For all analyses, two regions were considered overlapping when at least 50% of either region overlapped the other.

### Heritability enrichment analysis

To estimate heritability enrichment in each OCR, we conducted stratified-linkage disequilibrium score regression (S-LDSC) with LDSC (ver.1.0.1) using the baseline model (version 1.2; 53 categories). The p-value of the coefficient for cOCR (or relaxed cOCR) was obtained by adjusting for the 53 categories in the baseline model and all OCRs in each cell type. To avoid false positives ^83^, we used the relaxed cOCRs (the median proportion of single nucleotide polymorphism (SNPs) across the 29 cell types was 0.5%) for primary analyses instead of the cOCRs (the median was 0.07%). Multiple testing correction across all trait-cell type pairs was performed to estimate FDR using the Storey method ^84^. The summary statistics were obtained from the Alkes Price Lab (See URLs and our previous study ^9^), and five GWAS results in EAS ancestry: systemic lupus erythematosus ^85^, rheumatoid arthritis ^86^, Systemic sclerosis ^87^, Atopic dermatitis ^88^, and Takayasu arteritis ^89^. For this analysis, cOCRs’ positions were mapped to the hg19 coordinates using the UCSC liftOver tool with hg38ToHg19.over.chain file to be consistent with the summary statistics. After mapping to hg19, regions mapped to different chromosomes were excluded from the analysis. For the other constraint metrics, we extracted the strongly constrained region: Gnocchi Z > 4, phyloP46way score > 0.655, phyloP100way score > 9, phyloPPrimates score > 0.999, cactus241way score > 8.8, GERP RS > 5, DR rank <= 0.005. Thresholds for the other constraint metrics were selected to match the proportion of SNPs annotated by Gnocchi (1.18%), thereby ensuring comparable annotation sizes across constraint metrics.

### PRS analysis

The PRSs were constructed using 1,178 cases and 136,442 controls (137,620 individuals in total) and a traditional clumping-and-thresholding approach implemented in PLINK v1.9, based on summary statistics from an SLE GWAS of EAS ancestry ^85^. The 137,620 samples analyzed in PRS were not included in the sumstats. To select variants for PRS construction, we applied 20 GWAS P-value thresholds: 1, 0.9, 0.8, 0.7, 0.6, 0.5, 0.4, 0.3, 0.2, 0.1, 0.05, 0.01, 0.005, 5.0 × 10^-4^, 5.0 × 10^-5^, 1.0 × 10^-5^, 5.0 × 10^-6^, 1.0 × 10^-7^, 5.0 × 10^-7^ and 5.0 × 10^-8^. For each P-value threshold, linkage disequilibrium (LD) clumping was performed using nine r² thresholds (0.1, 0.2, 0.3, 0.4, 0.5, 0.6, 0.7, 0.8, 0.9) within a 250-kb window, resulting in 180 candidate PRS models.

We used five-fold cross-validation to select the optimal PRS parameters. The dataset was divided into five folds, with four folds used for training and the remaining fold used for testing in each iteration. All model selection was performed within the training folds and evaluated in the held-out fold. For each training set, PRSs were calculated for all 180 combinations of P-value and r² thresholds using the EAS SLE GWAS summary statistics. The AUC was calculated using the pROC package (version 1.18.4), and the combination of P-value and r² thresholds yielding the highest AUC was selected.

We then took the two variant sets; (a) variants filtered with the best parameters, and (b) all variants overlapping with DN B cOCRs as prioritized variants. To mitigate overfitting due to LD structures, we removed variants with LD>0.7 with variants in set (b). We then calculated the PRS for both sets (a) and (b) for each of the five test sets, and estimated the effect size of PRS from set (b), controlling for the PRS from set (a) using a logistic regression model. Finally, we performed a fixed-effects meta-analysis to evaluate the additional predictive contribution of the cOCR-prioritized variants.

### Fine-mapping enrichment analysis

We downloaded the UKB fine-mapped variants of 94 complex diseases and traits from the Finucane lab (<u>see URLs</u>) and obtained BBJ fine-mapped variants of 62 quantitative traits from a recent study ^49^. For the UKB dataset, we used PIPs estimated with SuSiE. The positions of those datasets were mapped to the hg38/GRCh38 coordinate using liftOver as described previously. In total, we obtained 830,654 trait-variant pairs for UKB and 544,745 trait-variant pairs for BBJ. The number of overlapped variants for each trait in 29 cell types of cOCRs was counted, and the OR was estimated using variants with PIP less than 0.1 and all OCRs in each of the 29 cell types as the background, and the P-value was calculated from the logistic regression model. Bonferroni correction was applied separately according to the number of phenotypes tested, using a significance threshold of 0.05 divided by the corresponding number of phenotypes. Among traits showing significant enrichment, median ORs were calculated separately for UKB and BBJ (Extended Data Fig. 5A, B; Supplementary Table 20).

Traits were classified as hematopoietic or non-hematopoietic, with hematopoietic traits further classified as leukocyte or non-leukocyte traits. Trait classifications and the numbers of putative causal variants in each category are provided in Supplementary Table 21. We then evaluated the enrichment of fine-mapped variants for leukocyte, non-leukocyte hematopoietic, and non-hematopoietic traits in cOCRs from each of the 29 cell types. We calculated the OR using the method described above.

For the rare-variant enrichment analysis, we merged cOCRs across the 29 cell types to generate a non-redundant set of cOCRs (merged cOCRs). We restricted the analysis to rare variants with MAF < 0.01 and classified them according to PIP (PIP ≤ 0.1 as background) and overlap with merged cOCRs. The OR was calculated, and one-sided P-value was assessed using Fisher’s exact test. Analyses were performed for all traits and separately for hematopoietic traits.

### Catalog cell-type-specific putative causal variants

We first selected putative causal variants from credible sets in which only one variant overlapped a cOCR. We then identified variants that overlapped cOCRs from only one cell type. This yielded 186 and 62 cell-type-specific putative causal variants in the EUR and EAS datasets, respectively.

We next extended the definition of cell-type specificity to the subgroup level and identified variants that overlapped cOCRs from only one immune cell subgroup (CD4, CD8, B, Mono, DC, NK, or Neutrophil; Supplementary Table 1). Combining them, we obtained a total of 1,210 and 323 trait-associated cell-type-specific putative causal variants in the EUR and EAS datasets, respectively.

### Statistical analysis

When standard floating-point arithmetic in R provided insufficient numerical precision for extremely small P values, high-precision arithmetic implemented in the Rmpfr package (version 0.9.2) was used to calculate P values from z-scores. For meta-analyses, pooled odds ratios (ORs) were generally estimated using a random-effects model implemented in the metafor R package (version 2.0-0). For the PRS analysis, effect estimates for the cOCR-prioritized PRS across the five test folds were combined using a fixed-effects meta-analysis implemented in METAL (version released on 5 May 2020).

## Data availability

The CAMBUS scores for all analyzed open chromatin regions across 29 immune cell types will be made publicly available upon publication.

## Code availability

Code used in this analysis is available at https://doi.org/10.5281/zenodo.18466265

## URLs

Gnocchi is at https://gnomad.broadinstitute.org/downloads#v3-genomic-constraint. Enrichment for Gnocchi with the known cCREs is at gs://gnomad-nc-constraint-v31-paper/fig_tables/constraint_z_genome_1kb.annot.txt. ENCODE cCREs V3 is at https://screen.encodeproject.org/. FANTOM5 enhancer is at https://zenodo.org/record/556775/. Super enhancer is at https://bio.liclab.net/sedb/download.php. phylop46way is at https://hgdownload.cse.ucsc.edu/goldenPath/hg19/phyloP46way/primates<u>. phyloP100way</u> is at https://hgdownload.cse.ucsc.edu/goldenpath/hg38/phyloP100way/hg38.phyloP100way.bw phyloPPrimates is at https://cgl.gi.ucsc.edu/data/cactus/zoonomia-2021-track-hub/hg38/phyloPPrimates.bigWig. cactus241way is at https://hgdownload.soe.ucsc.edu/goldenPath/hg38/cactus241way/cactus241way.phyloP.bw. GERP is at ftp.ensembl.org/pub/current_compara/conservation_scores/91_mammals.gerp_conservation_score/gerp_conservation_scores.homo_sapiens.GRCh38.bw. DR is at https://static-content.springer.com/esm/art%3A10.1038%2Fs41586-022-04965-x/MediaObjects/41586_2022_4965_MOESM3_ESM.gz.GWAS summary statistics are at https://alkesgroup.broadinstitute.org/sumstats_formatted, and https://alkesgroup.broadinstitute.org/UKBB/. ABC predictions are at https://mitra.stanford.edu/engreitz/oak/public/Nasser2021/AllPredictions.AvgHiC.ABC0.015.minus150.ForABCPaperV3.txt.gz. UKB fine-mapped variants are at https://www.finucanelab.org/data. FANTOM5 tissue-expressed data is at https://zenodo.org/records/5348471. HAQERs is at https://www.vertgenlab.org/haqer2022.html.

## Acknowledgment

We would like to thank all the participants in the study. We deeply thank all the members of the Laboratory for Statistical and Translational Genetics, RIKEN Center for Integrative Medical Sciences for the technical support. The super-computing resource was provided by Human Genome Center, the Univ. of Tokyo (http://sc.hgc.jp/shirokane.html). We acknowledge the use of Biorender for aspects of the figures. This work was supported by Japan Agency for Medical Research and Development (AMED) grants 21ek0109555, 21tm0424220, 21ck0106642, 23ek0410114, and 23tm0424225. Japan Society for the Promotion of Science (JSPS) KAKENHI grant JP20H00462 and JP23K14164, Takeda Hosho Grants for Research in Medicine, the Medical Research Support Project of the Shizuoka Prefectural Hospital Organization and JSPS program for Forming Japan’s Peak Research Universities (J-PEAKS) JPJS00420240019.

## Author contributions

K.T., M.K., and C.T. conceived the study design. A.S. performed ATAC-sequencing and DNA genotyping. K.T. and N.T. performed whole genome imputation. K.T. analyzed this study with the help of S.Y., N.T., Y.I., X.L., S.K., and M.K. M.K. and C.T. supervised this study. S.Y., N.T., S.K., K.I., Y.M., Immune transcript/enhancer consortium, and K.Y. provided helpful comments and discussion. K.T., M.K., A.S., and C.T. wrote the manuscript.

## Ethics declarations

Competing interests

The authors declare no competing financial interests.

## Extended Data Figure

**Extended Data Fig. 1.**
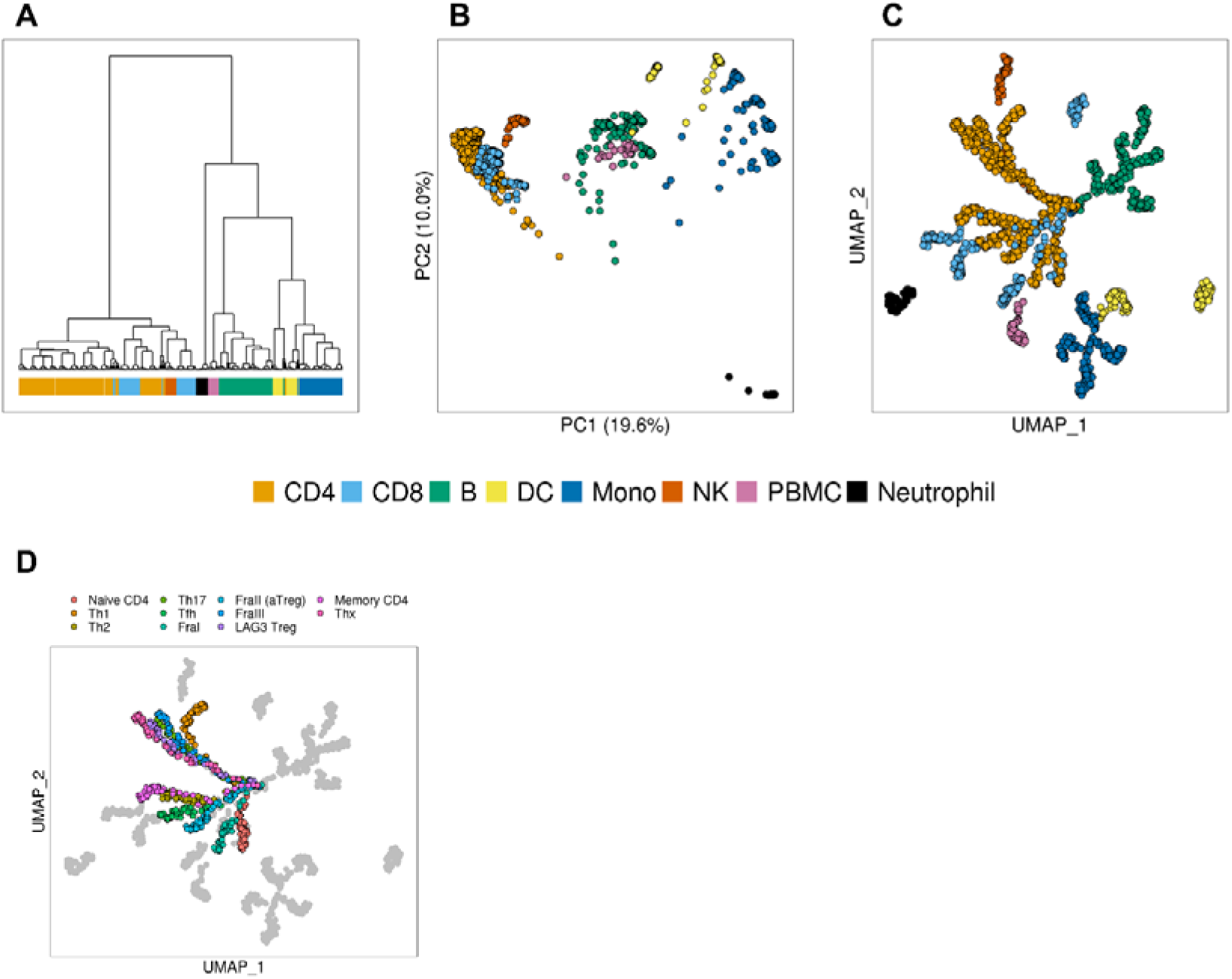
Clustering analysis and dimension reduction using PCA and UMAP. A-C, (**A**) Hierarchical clustering, (**B**) PCA, and (**C**) UMAP of 1,386 samples. The top 5,000 most accessible OCRs were used for these analyses. The colors indicate the classification of samples into eight subgroups across 29 cell types. **D,** UMAP of 1,386 samples (**C**) colored by eleven types of CD4; other subgroups are colored with grey.

**Extended Data Fig. 2.**
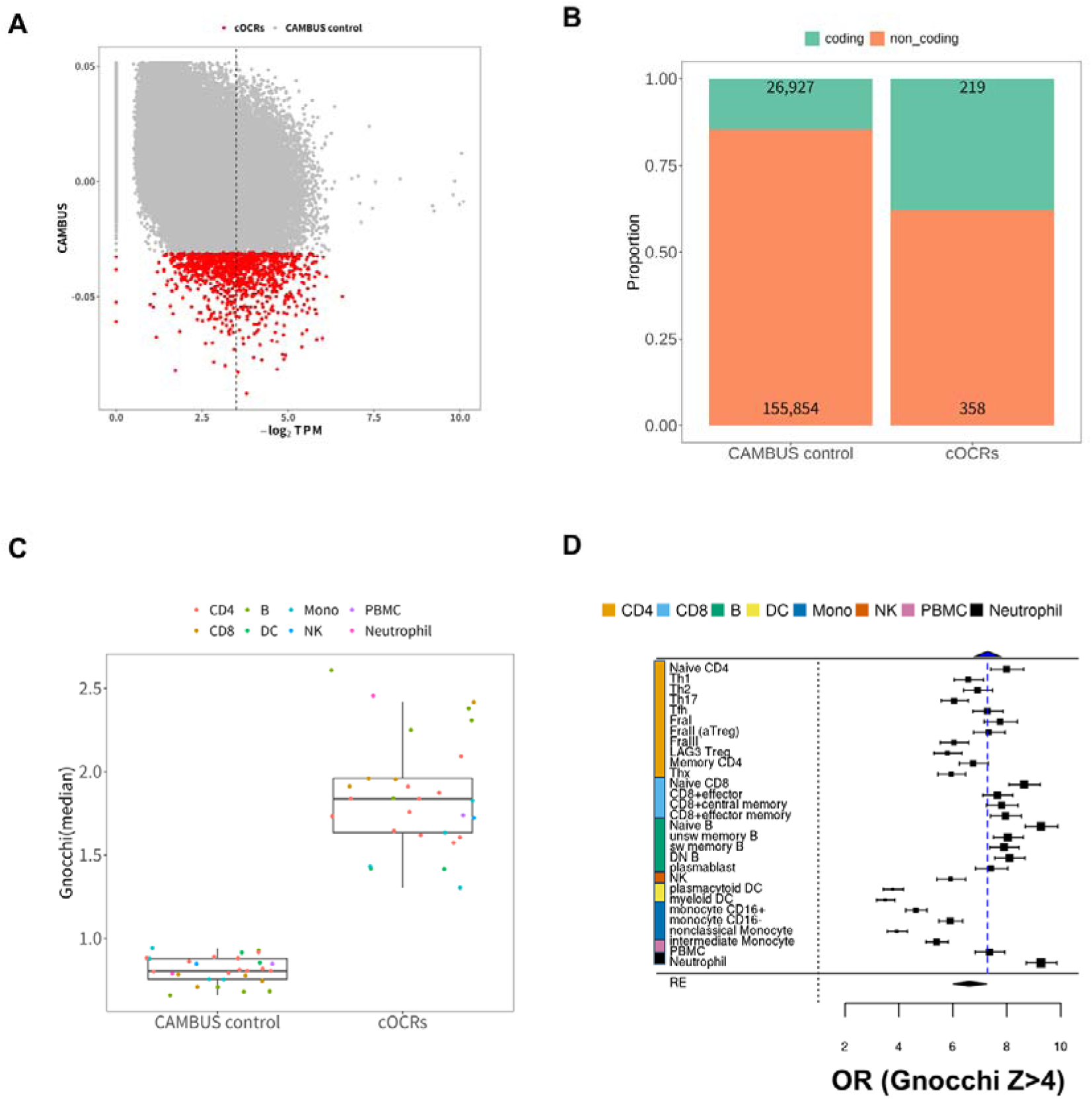
cOCRs are enriched for evolutionary constraint. **A,** Scatter plot of median TPM and CAMBUS in Naïve CD4. The vertical dashed line represents the top 5% of the median TPM. **B,** The number of OCRs in coding and non-coding regions for each cOCRs and CAMBUS control OCRs. The regions of coding and non-coding are referred to previous study ^27^. **C,** The distribution of non-coding Gnocchi (median) for each cOCRs and CAMBUS control OCRs in each of the 29 cell types. **D,** OR of relaxed cOCRs across 29 cell types in Gnocchi> 4 and pooled OR estimated through random-effects meta-analysis. The blue dashed line represents the OR of ENCODE promoter-like elements. The dashed line represents no enrichment (OR=1.0)

**Extended Data Fig. 3.**
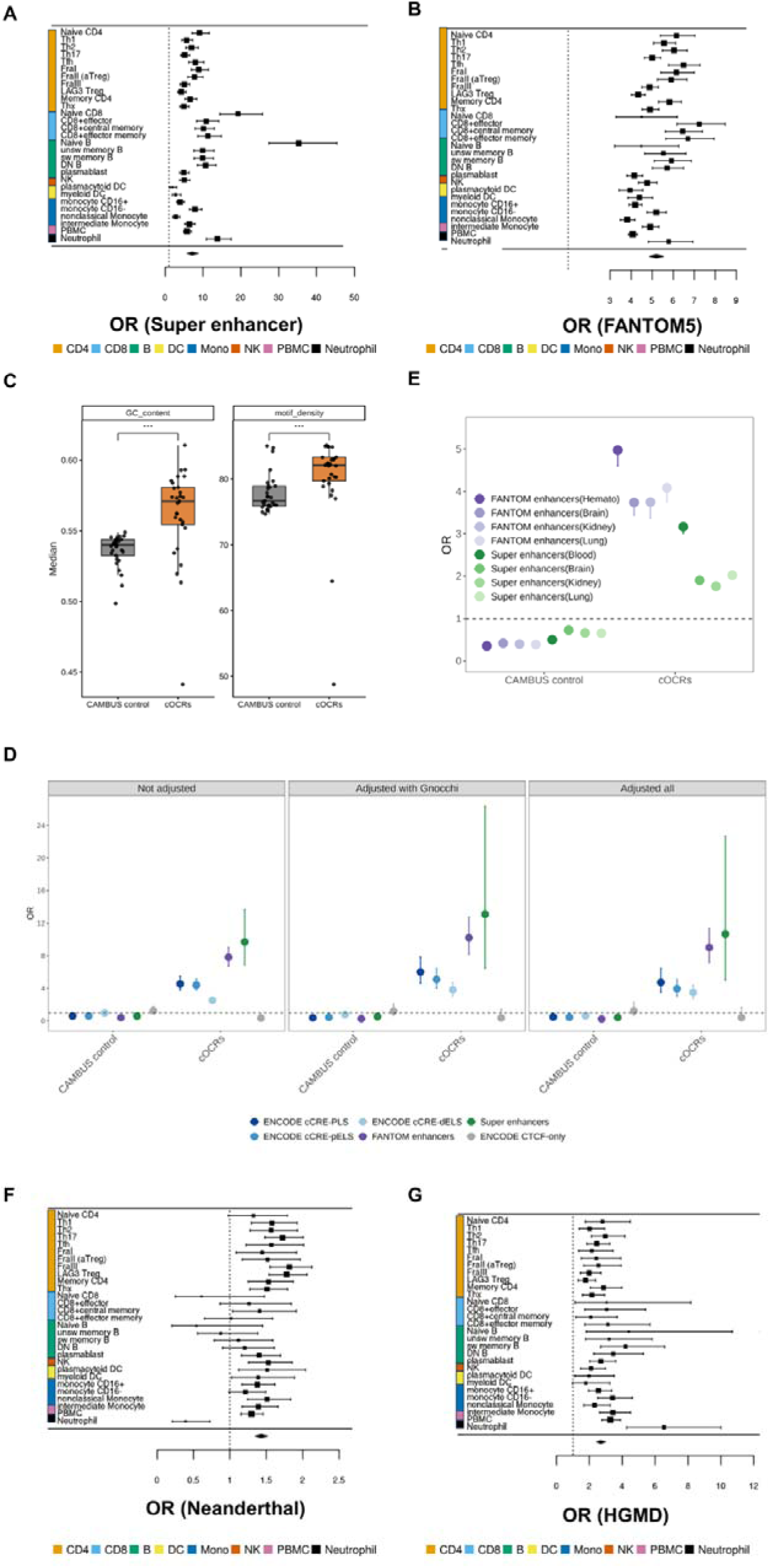
cOCRs are enriched for active regulatory elements. **A, B,** pooled OR of the cOCRs in super-enhancers **(A),** FANTOM5 enhancers **(B). C,** Median GC content (left panel) and TF motif density (right panel) in cOCR and CAMBUS control OCR across 29 cell types. One-sided Wilcoxon-test were performed between cOCR and CAMBUS control OCR with significance levels denoted as P < 0.001 (‘***’). **D,** Enrichment of non-coding cOCRs for NaiveCD4 (n = 865) in the known cCREs (ENCODE cCRE-PLS; n = 40,891, ENCODE cCRE-pELS; n = 172,027, ENCODE cCRE-dELS; n = 789,200, ENCODE CTCF-only; n = 35,839, FANTOM5 enhancers; n = 63,280, Super enhancers; n = 16,472), adjusted for seven constraint metrics (Gnocchi, phyloP46way, phyloP100way, cactus241way, phyloPPrimates, GERP and DR), GC content, and TF motif density. **E,** ORs of relaxed cOCRs (NaiveCD4; n = 12,736) in cell-type-specific regulatory elements: the tissue expressed FANTOM5 enhancers (hematopoietic cells; n = 64,225, Brain; n = 62,978, Kidney; n = 35,177, Lung; n = 26,630), and super-enhancers (Blood; n = 8,055, Brain; n = 9,372, Kidney; n = 4,712, Lung; n = 9,878). **F, G,** pooled OR of the cOCRs in Neanderthal genome sequences derived from samples of EAS ancestry (**F**), pathogenic regulatory variants (**G**). The dashed line represents no enrichment (OR=1.0; A, B, D-G).

**Extended Data Fig. 4.**
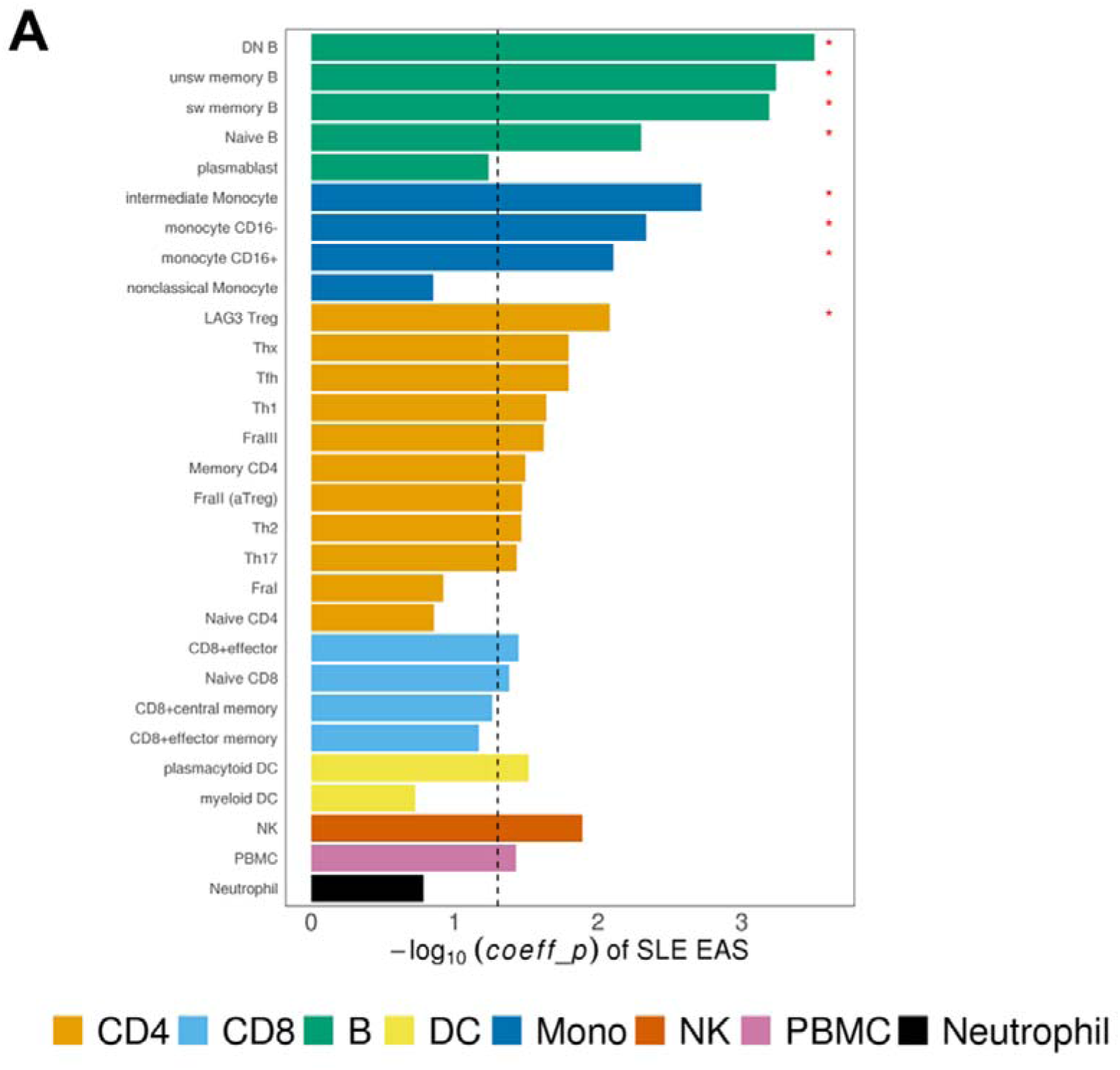
Heritability enrichment for immune-mediated diseases. **A,** The -log10-scale coefficient P of cOCRs in 29 cell types for SLE GWAS in EAS ancestry. The dashed line indicates a coefficient P = 0.05. ‘*’ indicates significance (qvalue < 0.05) after FDR correction.

**Extended Data Fig. 5.**
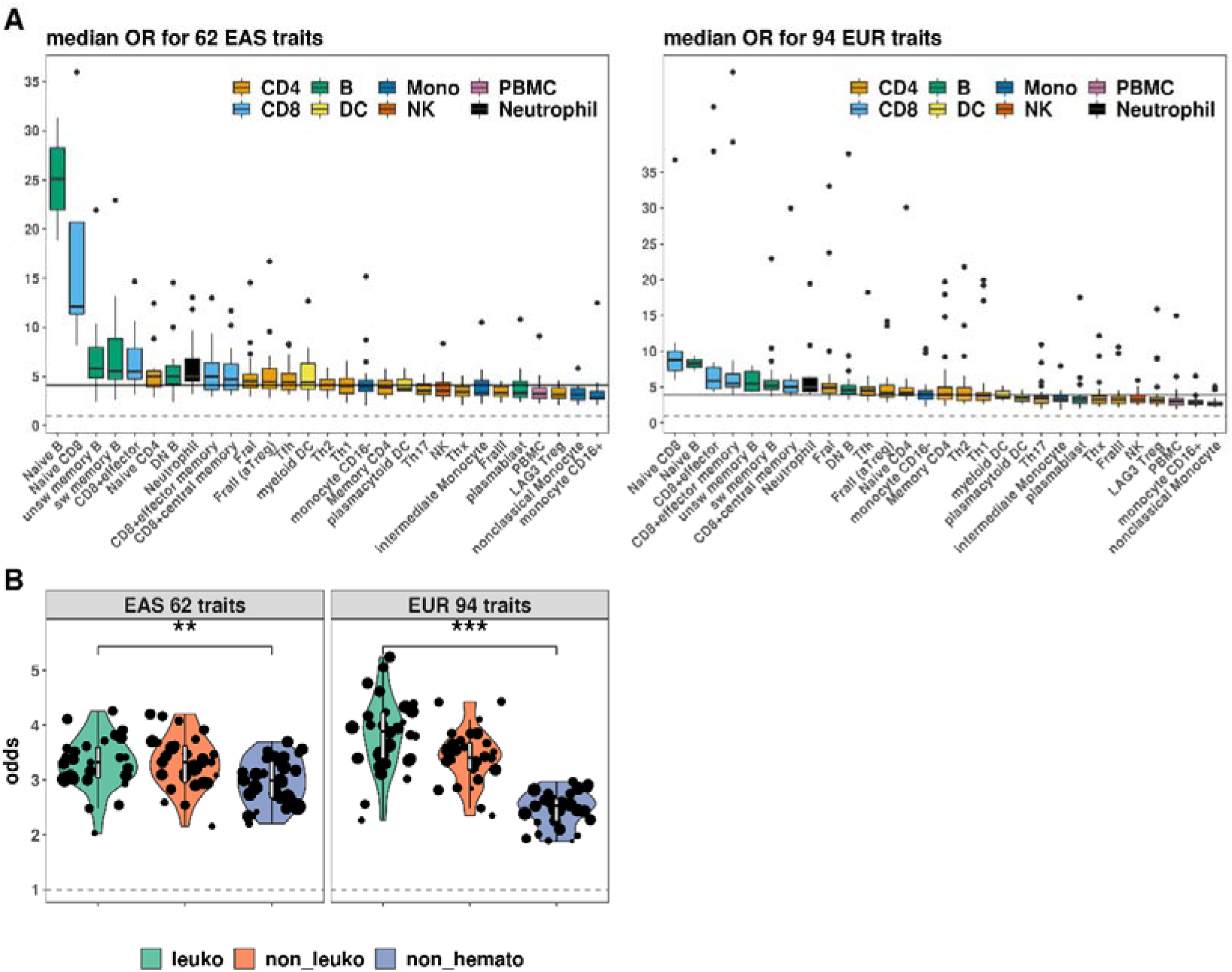
Prioritization of putative causal variants using cOCRs. **A,** Median OR of all variants in 95% credible sets in cOCRs across 29 cell types. We used putative causal variants from statistical fine-mapping studies in EUR and EAS ancestries ^48,49^. The solid line represents the median OR across 29 cell types. **B,** Enrichment of the putative causal variants in leukocyte (leuko), non-leukocytes (non_leuko), and non-hematopoietic (non_hemato) traits in cOCRs across 29 cell types. Only cell types with P < 0.05 are shown (27 cell types in BBJ leuko; 28 in UKB non_hemato; 29 in BBJ non_leuko, BBJ non_hemato, UKB leuko, and UKB non_leuko), and dots indicate -log10-scale p-value. t-tests were performed between leuko and non_hemato traits, with significance levels denoted as P <0.05 (‘*’) and P < 0.001 (‘***’).

**Extended Data Fig. 6.**
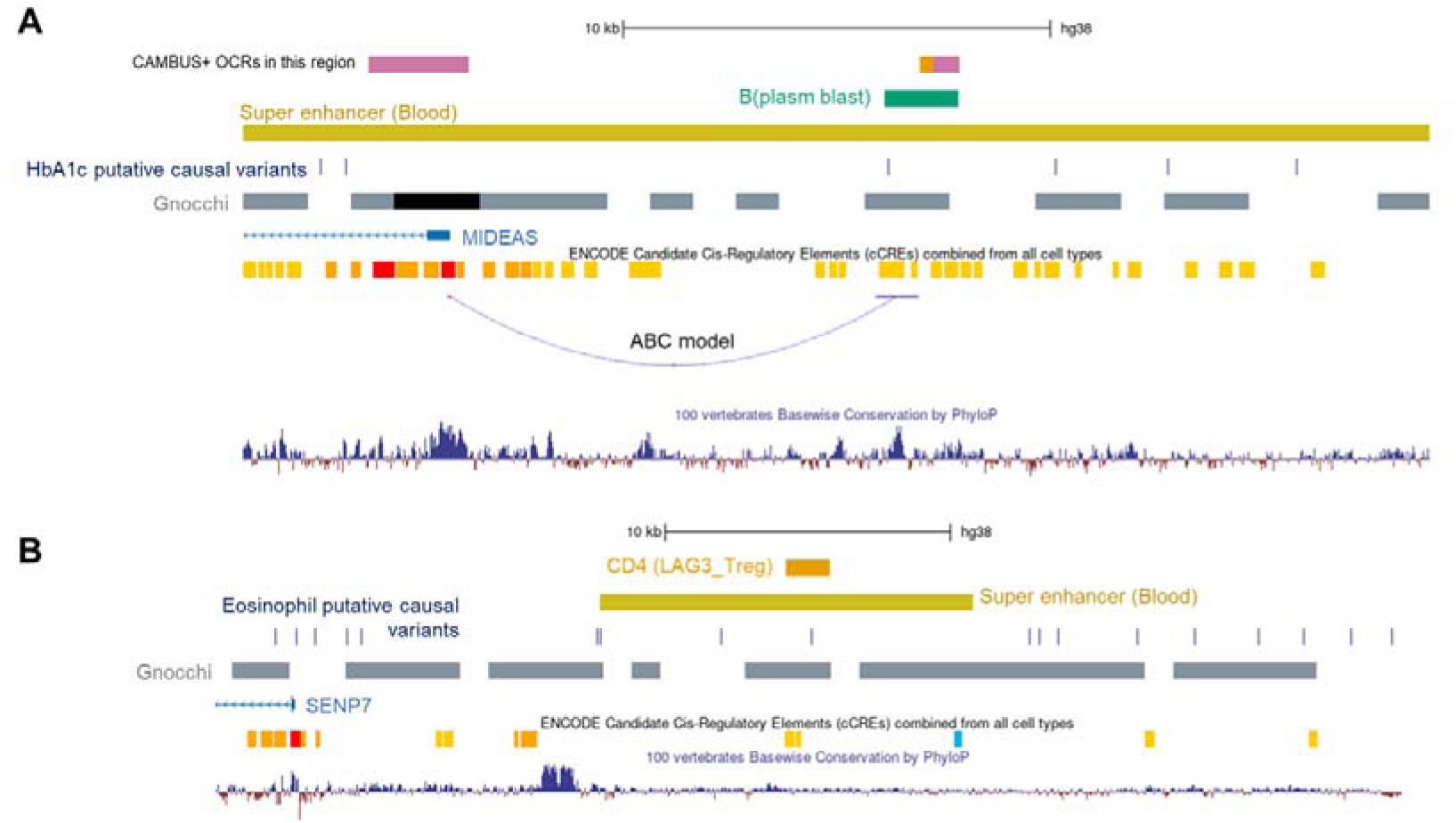
Prioritization of putative causal variants using cOCRs. **A,B,** representative examples of variants associated with Hemoglobin A1c level prioritized with cOCRs in plasmablast in putative causal variants in EUR ancestry (**A**) and eosinophils prioritized with cOCRs in regulatory T-cell (**B**) and their surrounding genomic annotations. The navy boxes represent putative causal variants in the credible set. The blue dashed line represents an enhancer-gene connection to the nearest protein-coding gene predicted by the ABC model in B-cells (**A**). All the other cOCRs around putative causal variants (rs8009224 (**A**)) are also shown in the figures (The purple box represents PBMC).

**Extended Data Fig. 7.**
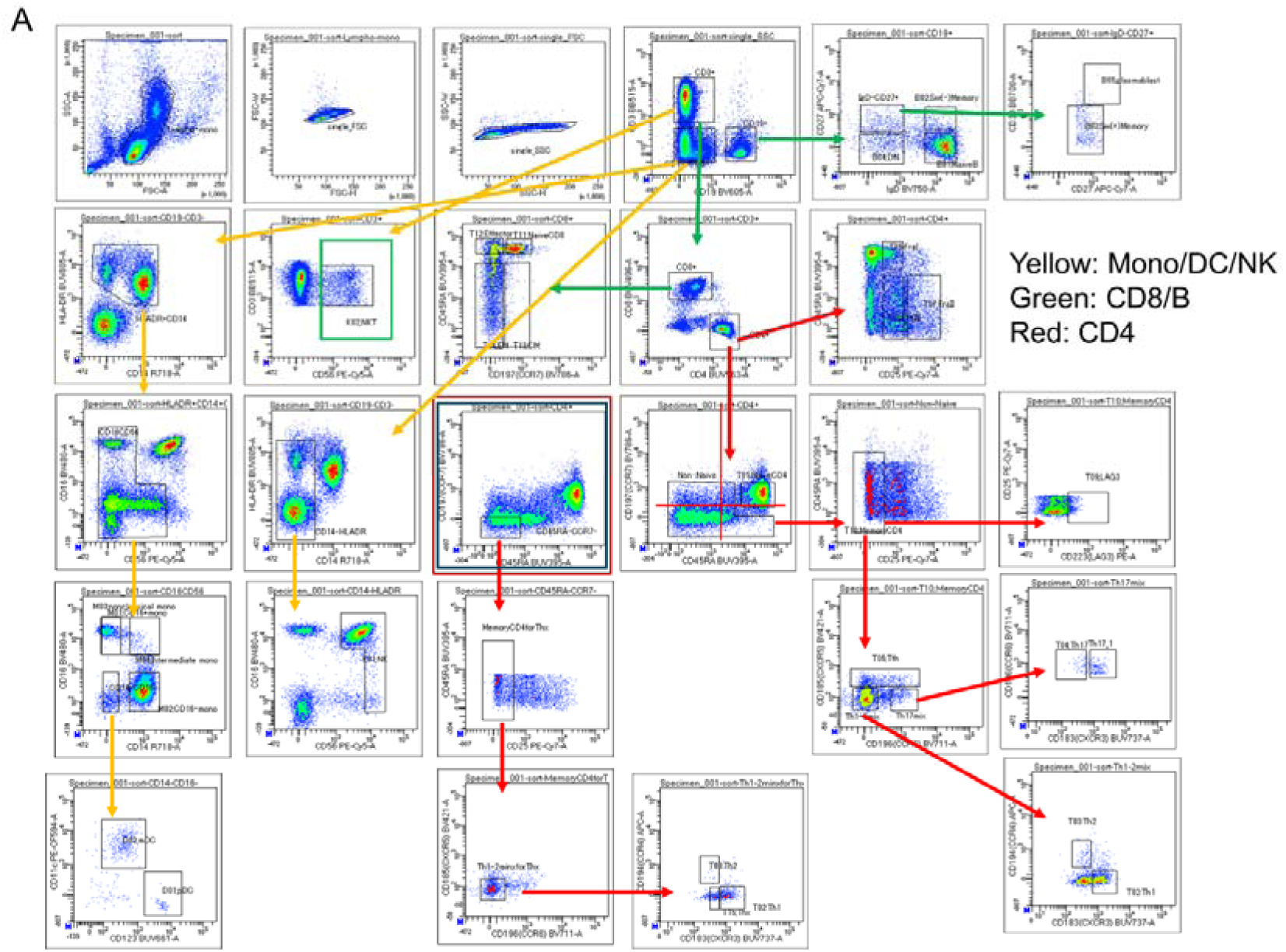
Sorting strategy for immune cell subsets. **A,** Representative FACS plots describing gating strategies for 27 cell types. See Supplementary Table 22 for flow cytometry antibodies used in this study.

