## Supplementary Note for "Sequence sensitivity of open chromatin reveals constrained regulatory elements underlying human traits"

#### Contents

Supplementary Notes

Supplementary Methods

Supplementary Figures

Supplementary Tables

#### Supplementary Notes

##### Cell-type-specific OCR predictions from DNA sequences

We first examined whether CAMBUS ML (Chromatin Accessibility Mutation Burden Score machine learning) models can predict the OCRs of each of 29 cell types from DNA sequence patterns. For this purpose, we used a deep convolutional neural network to predict whether OCRs overlap within the central 200bp of 5kb input DNA sequences (Methods). We observed a notably high mean area under the receiver operating characteristic curve (AUROC) of 0.87, with a minimum of 0.84 (neutrophils), to predict OCRs across 29 cell types in the testing dataset (Supplementary Fig. 1A). We observed that cell-type-specific OCRs were more challenging to predict; however, even for the OCRs observed in only one cell type, the mean AUROC was 0.74 (Supplementary Fig. 1B), suggesting CAMBUS ML models could effectively capture cell-type-specific genetic architecture underlying OCRs. Notably, this level of prediction accuracy has been enough for successful downstream analyses (such as *in* *silico* mutagenesis for GWAS variants) in the previous studies: for example, AUROC per cell = 0.762 [1] and mean AUROC for enhancer RNAs = 0.69 [2].

We also assessed the accuracy of *in silico* mutagenesis on the cell-type-specific OCRs in the 29 cell types. The direction concordance between the *in silico* mutation effects and the estimated effect sizes from caQTL increased at a higher threshold of the mutation effects, as shown in previous studies [2,3]. Particularly when variants were selected based on a threshold where accessibility probability changed by more than 10%, indicating high confidence of *in silico* mutagenesis, the direction concordance with caQTL effect sizes (Supplementary Methods) reached 90% for the six major cell types (CD4, CD8, B, Mono, DC, and NK) (Supplementary Fig. 1C; Supplementary Table 2). We also confirmed the good concordance using the independent caQTL study of lymphoblastoid cell lines (LCL; B cell lineage; Supplementary Fig. 1D, Fig. 2A) [4]. As expected, the concordance dropped when we compared *in silico* mutation effects in non-B cell types with the caQTL study (Supplementary Fig. 2B).

We further extended this validation to another independent caQTL study [5] comprising five major immune cell types (T cells, B cells, monocytes, NK cells, and PBMCs) to assess their concordance across immune cell types beyond B cells. Remarkably, we observed highly concordant directions between mutation effects and caQTL effect sizes in matched cell types (Supplementary Fig. 3A). In line with this observation, the concordance rate tended to increase as the precision of both mutation effects and caQTL effect sizes improved (Supplementary Fig. 3B).

Thus, CAMBUS ML models incorporating surrounding DNA sequence patterns could accurately predict OCRs within the 29 cell types, including cell-type-specific ones, and mutation effects on these OCRs in a cell-type-specific manner.

**Evaluation of alternative sequence-based model architectures**

We compared the DeepSEA-based CAMBUS ML model used throughout the main analysis (hereafter, the DeepSEA-based model) with a CAMBUS ML model based on Sei [6] (hereafter, the Sei-based model) to evaluate whether a more recent sequence-based regulatory prediction architecture could improve CAMBUS performance (Supplementary Method). To assess both predictive accuracy and computational cost for large-scale *in silico* saturated mutagenesis, we trained both architectures under a practical computational budget of up to two weeks. At two weeks, the Sei-based model was still being trained and had reached 288,000 training steps, whereas the DeepSEA-based model had already converged and was approximately 22-fold faster at the same number of training steps. We then compared the two models in terms of test-set prediction accuracy and concordance between the effects predicted for the B-cell lineages by *in silico* mutagenesis and caQTL effect directions for LCLs [4]. The DeepSEA-based model showed slightly higher test-set prediction accuracy, with a mean AUROC of 0.868 versus 0.854 for the Sei-based model. It also showed generally higher direction concordance across B-cell lineages, particularly for variants with large absolute *in silico* mutation effects (e.g., 0.87 for the DeepSEA-based model versus 0.81 for the Sei-based model in Naive B-cells at a >10% change in predicted accessibility probability; Supplementary Fig. 4A).

We further benchmarked the DeepSEA-based architecture against ChromBPNet [7], one of the state-of-the-art models for chromatin accessibility prediction. Because ChromBPNet requires BAM files for model training, and the BAM files used in the main analyses could not be reused for this benchmarking purpose owing to their data-use restrictions, we performed the comparison using ENCODE K562 ATAC-seq data (Supplementary Method). We trained each architecture with its default parameters (Supplementary Table 23). We did not report AUROC between the two models as a standard evaluation metric here, because they are trained for different prediction tasks (DeepSEA predicts OCR presence, whereas ChromBPNet predicts quantitative base-resolution accessibility profiles and total counts). Instead, we performed *in silico* mutagenesis and compared their direction concordance with effect estimates from the independent caQTL dataset described above. Overall, the direction concordance was almost comparable between the two models and increased as the analysis was restricted to variants with larger absolute predicted mutation effects. Notably, among the top-ranked variants, concordance was slightly better for DeepSEA-based model than for ChromBPNet (0.94 versus 0.79 for the top 100 variants, respectively; Supplementary Method; Supplementary Fig. 5A). Furthermore, on a single NVIDIA L4 GPU under their respective standard configurations (Supplementary Method), DeepSEA-based model required substantially less time for *in silico* mutagenesis than ChromBPNet (approximately 400 sec versus 1 hour, respectively), whereas their training times were comparable (approximately 17 hours for both models). Taken together, these results support the use of the DeepSEA-based architecture as the primary model for CAMBUS, given its comparable predictive performance for the purpose and substantially lower computational cost.

#### Supplementary Method

##### Validation of the CAMBUS ML model with *in silico* mutagenesis

We compared the *in silico* mutation effects predicted by the CAMBUS ML model with effect sizes from the caQTL study. The samples for caQTL (n=74, including additional independent samples to increase statistical power) were genotyped using the Infinium ASA-24+v1.0 kit, and whole-genome imputation was conducted after stringent quality control (Supplementary Table 18). The caQTLs were mapped using RASQUAL (v1.1) with default parameters. The permutation to empirically obtain the null distribution for each region was performed with default parameters, resulting in a median of 654 lead caQTLs with FDR $< 0.1$, which we compared against the *in silico* mutation effects. Additionally, we calculated the *in silico* mutation effects for 297,308 autosomal lead SNVs in the previous study's caQTL (see URLs). To further refine the comparison, we filtered the target caQTLs within the peak. In total, we used 155,989 caQTL peak pairs to compare with the mutation effect.

For additional validation, we compared the mutation effects with caQTL effect sizes identified from single-cell ATAC-seq in immune cells. We extracted 6,901 lead caQTL variants (FDR<10%; see URLs), including 2,128 in T cells, 666 in B cells, 841 in NK cells, 3,507 in monocytes, and 4,662 in PBMCs. For each cell type, the precomputed mutation effects from the corresponding cell-type-specific CAMBUS ML model were compared with the effect sizes of lead caQTL variants in matched OCR peaks, in the same manner as above. In total, we used caQTLs from T cells (n = 2,090; matched to naive CD4 T cells), B cells (n = 666; naive B cells), NK cells (n = 834; NK cells), monocytes (n = 3,411; CD16+ monocytes), and PBMCs (n = 4,520; PBMCs) to evaluate concordance with the predicted mutation effects.

All the positions of caQTLs in the previous studies were mapped to the hg38/GRCh38 coordinates using the liftOver tool with the hg19ToHg38.over.chain file to be consistent with our study.

**Estimation of computational time for *in silico* mutagenesis using Evo 2**

To calculate CUMBUS across millions of OCRs, we performed *in silico* mutagenesis for 15,893 M variants using CAMBUS ML, which required about two months of computation on five Quadro GV100 with 32GB memory (NVIDIA) and three Tesla P100 with 16GB memory (NVIDIA). To estimate the computational demand of performing the same analysis using Evo 2 (an open source, one of the state-of-the-art DNA foundation model, trained on a representative snapshot of genomes spanning all observed evolution) [8] instead of CAMBUS ML, we installed Evo 2 on an H100 GPU with 96GB of memory (NVIDIA) following the instructions in its repository (https://github.com/ArcInstitute/evo2/blob/a796302818055b9710a6a2c4d7882a6243363fdd/README.md).

Using the evo2_7b model (Evo2 with 7 billion parameters), we observed that processing 1,000 variants required approximately 450 seconds. Based on this empirical runtime, we extrapolated that processing 15,893 M variants would take approximately more than 200 years using an H100 GPU. Thus, to calculate CAMBUS across millions of OCRs obtained from ATAC-seq, our strategy of using CAMBUS-ML, primarily based on computationally efficient convolutional networks, is a practical and scalable choice.

**Comparison of a Sei-based prediction model**

We compared Sei with DeepSEA, the model architecture used in CAMBUS-ML, by training a Sei-based model on the same OCR dataset using the Selene library (v0.6.0). Although recently developed large-scale AI models, such as Evo2, may provide powerful sequence representations, their computational requirements make genome-wide saturated *in silico* mutagenesis impractical for the scale required in this study. We therefore chose Sei as a recent model architecture that was feasible to train and apply under our computational setting. We installed the latest version of Sei on 2026/4/6 following the instructions in its repository (https://github.com/FunctionLab/sei-framework). Both the DeepSEA-based and the Sei-based models were trained on four NVIDIA L4 GPUs with 24GB memory to ensure matched computational conditions. The DeepSEA-based model completed training within three days, whereas the Sei-based model had not completed training after two weeks. Considering the trade-off between model performance and computational cost, we terminated Sei training after two weeks and used this checkpoint for comparison. The details of Selene's parameters are described in Supplementary Table 17.

**Comparison of a ChromBPNet-based prediction model**

To benchmark the predictive accuracy and computational efficiency of the DeepSEA-based model used in CAMBUS-ML against a state-of-the-art chromatin accessibility prediction model, we compared DeepSEA with ChromBPNet using the ENCODE K562 dataset described in the ChromBPNet tutorial (see URLs). Because the BAM files used in the main CAMBUS analyses could not be reused for this benchmarking purpose owing to their data-use restrictions, we instead used the publicly available ENCODE K562 dataset, which also allowed both models to be evaluated using the standardized dataset and workflow provided in the ChromBPNet tutorial. We installed the latest version of ChromBPNet on 2026/8/20 following the instructions in its repository (https://github.com/kundajelab/chrombpnet). Both models were trained on a single NVIDIA L4 GPU using their default parameters to assess their performance under standard configurations (Supplementary Table 23). ChromBPNet was trained following the tutorial protocol (https://github.com/kundajelab/chrombpnet/wiki/Tutorial). To ensure consistency in the training data, DeepSEA was trained using the same 269,695 filtered peaks generated by the ChromBPNet pipeline.

We then calculated mutation effects using the 155,993 autosomal lead variants located within the peak extracted from a previous caQTL study. Variants were ranked separately by the absolute magnitude of the predicted mutation effect sizes for each model since the scales of predicted mutation effects differ between DeepSEA and ChromBPNet. We then selected the top 100, 200, 500, 1,000, 2,000, 5,000, 10,000, and 20,000 variants from each model, and the concordance between the predicted mutation-effect direction and the experimentally observed caQTL effect direction was evaluated.

#### URLs

caQTL from ATAC-seq in LCL is at https://doi.org/10.5281/zenodo.1405945. caQTL from scATAC-seq in immune cell types is at https://zenodo.org/records/7375095. Bam file and peak data from ENCODE ATAC-seq in K562 is at https://www.encodeproject.org/files/ENCFF077FBI/@@download/ENCFF077FBI.bam, https://www.encodeproject.org/files/ENCFF128WZG/@@download/ENCFF128WZG.bam, https://www.encodeproject.org/files/ENCFF534DCE/@@download/ENCFF534DCE.bam, https://www.encodeproject.org/files/ENCFF333TAT/@@download/ENCFF333TAT.bed.gz.

#### Supplementary Figures

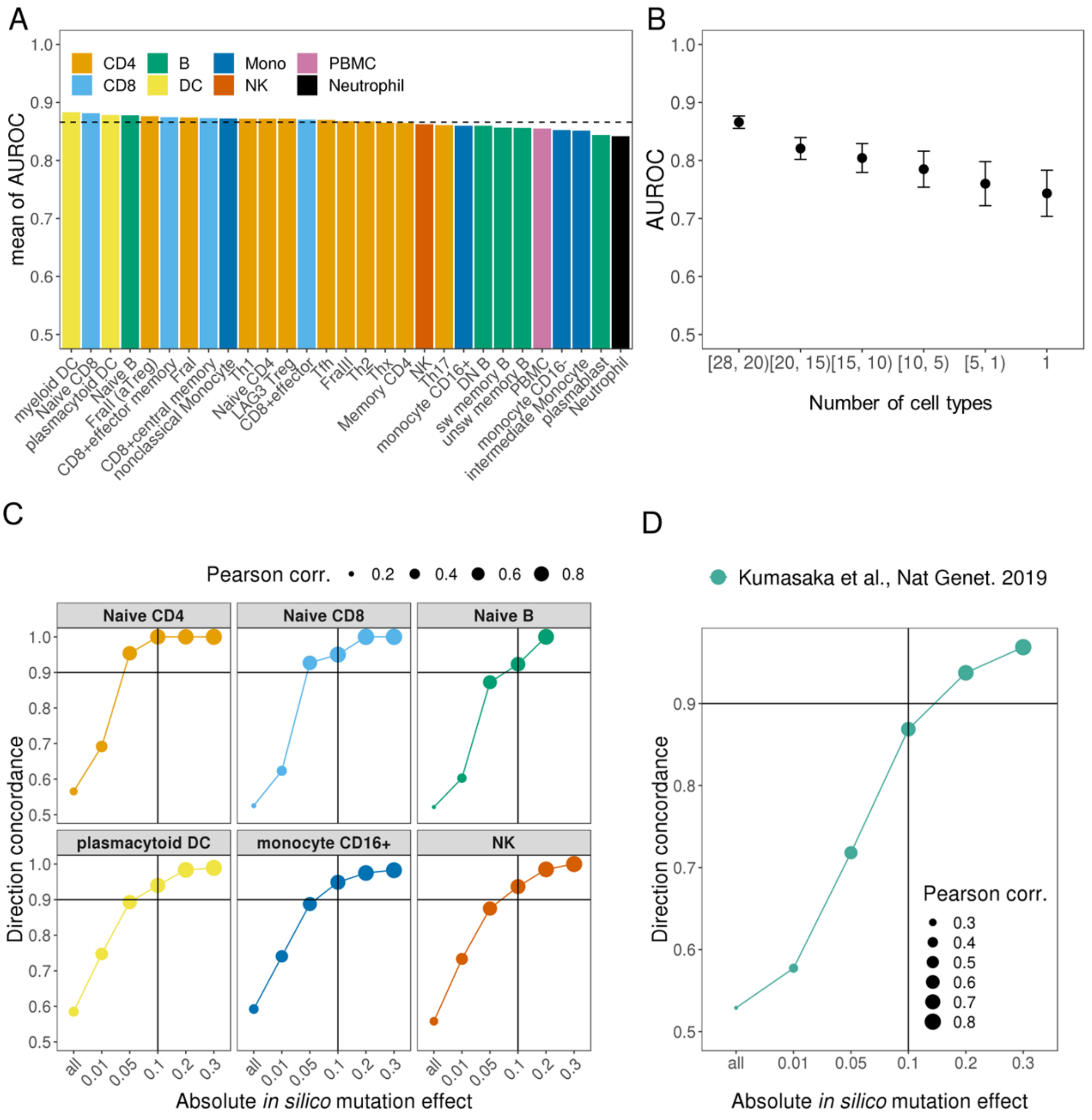


**Supplementary Fig. 1 | Accuracy of CAMBUS ML model and its mutation effect. A,** Predictive accuracy of the CAMBUS ML models across 29 cell types. The dashed line represents the mean AUROC across the 29 cell types (0.87). **B,** Specificity of CAMBUS ML models with respect to OCR sharing. The x-axis shows the number of cell types sharing OCRs. For example, the leftmost category represents OCRs shared among 20 to 28 cell types, and the rightmost category indicates OCRs unique to a single cell type. **C,** Direction concordance between *in silico* mutation effect from CAMBUS ML and effect size from our caQTL study. **D,** Direction concordance between *in silico* mutation effect from the CAMBUS ML for Naïve B and effect size from the independent LCL caQTL study. The Pearson correlation coefficient (Pearson corr.) was calculated to assess the relationship between *in silico* mutation effects and eQTL effect sizes.

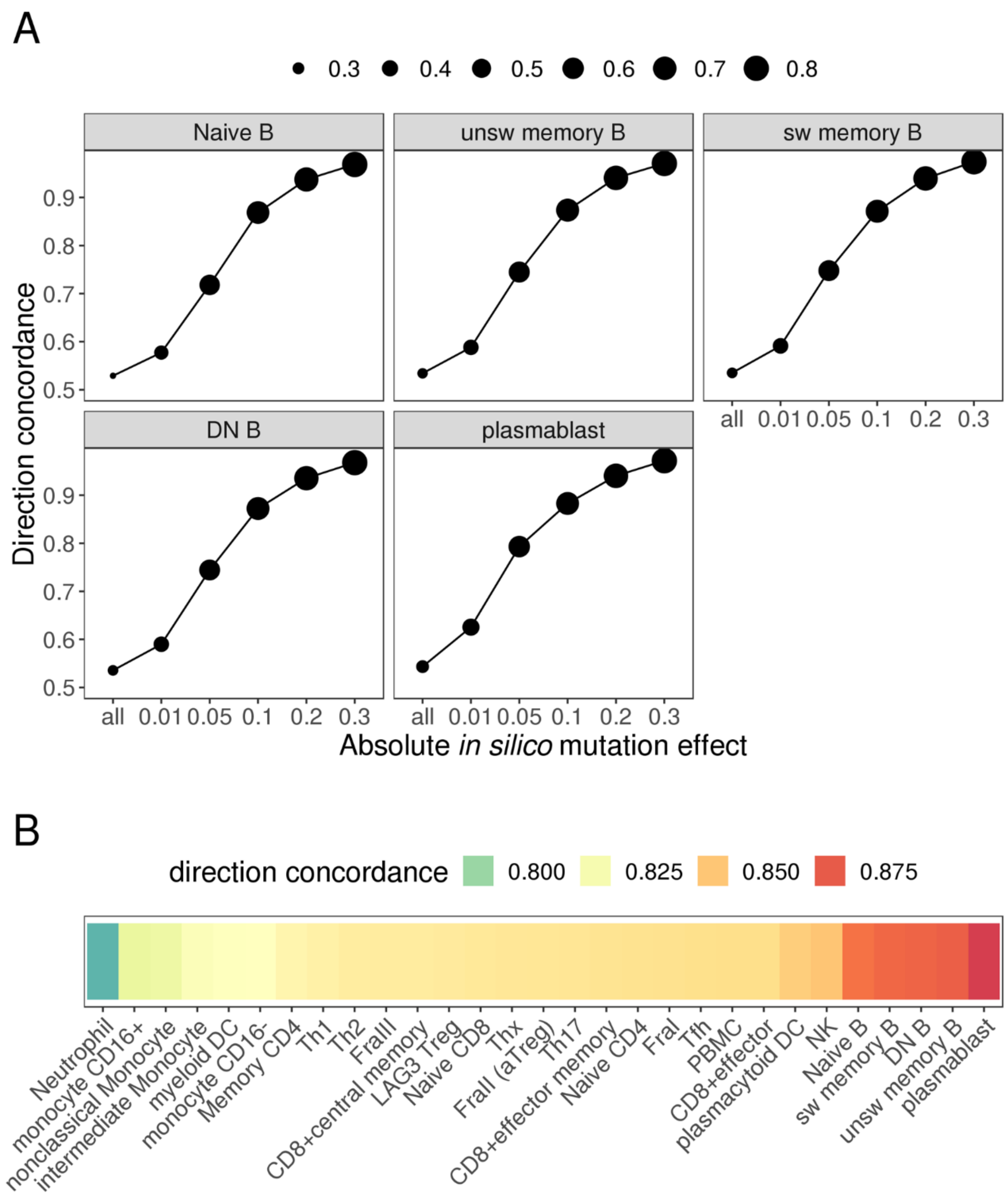


**Supplementary Fig. 2 | Accuracy of *in silico* mutation effects. A,** Comparison of direction concordance between *in silico* mutation effects predicted by the CAMBUS ML models (across five B-cell types) and effect sizes from an independent LCL caQTL study. The

Pearson correlation coefficient (Pearson corr.) was calculated to assess the relationship between *in silico* mutation effects and eQTL effect sizes. The x-axis represents a threshold of *in silico* mutation effects to calculate direction concordance, where 'all' indicates no threshold was applied. **B,** Comparison of direction concordance between *in silico* mutation effects from the CAMBUS ML models and effect size from the LCL caQTL study. Absolute *in silico* mutation effects greater than > 0.1 are used (see Supplementary Fig. 1C).

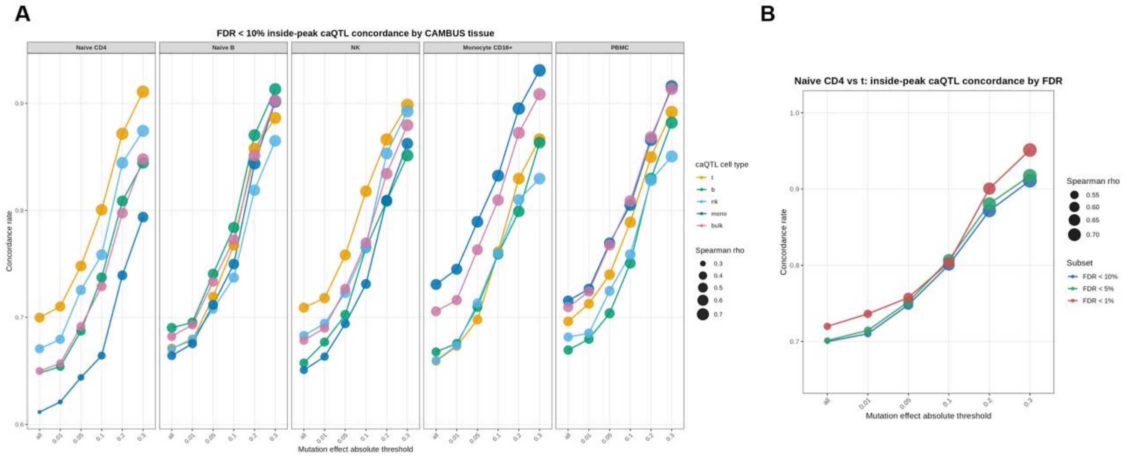


**Supplementary Fig. 3 | Accuracy of *in silico* mutation effect compared with single-cell caQTL effect sizes. A,** Directional concordance between *in silico* mutation effects predicted by the CAMBUS ML models across five cell types matching those analyzed in the caQTL study and effect sizes from an independent single-cell caQTL study of five major immune cell types. **B,** Concordance between Naive CD4 CAMBUS mutation effects and T cell caQTL effect sizes across caQTL significance thresholds (FDR<10%, <5%, <1%).

A

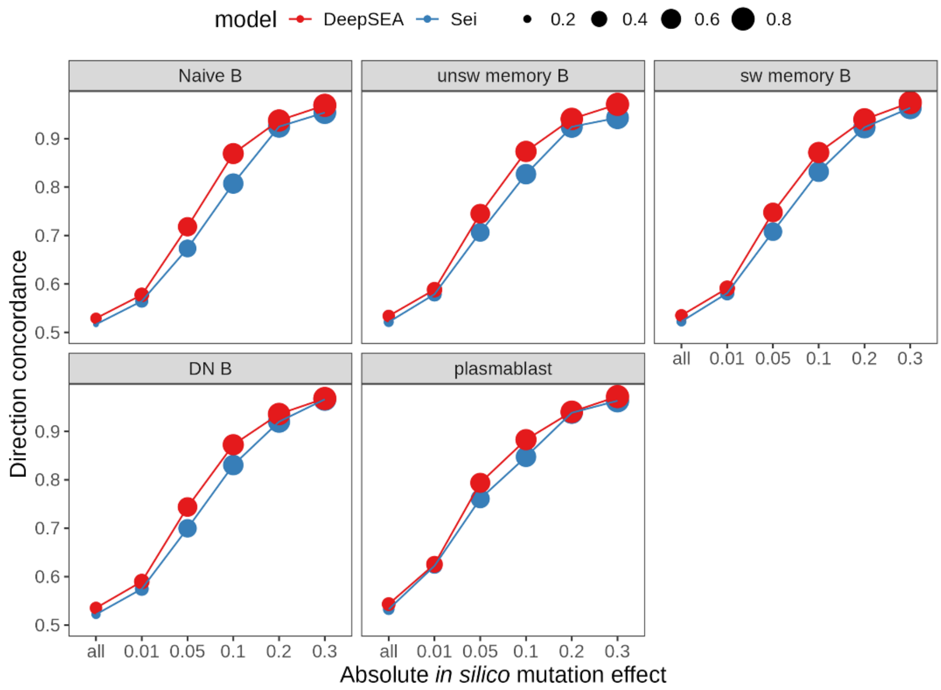


**Supplementary Fig. 4 | Comparison of a Sei-based prediction model. A,** Direction concordance between *in silico* mutation effects from CAMBUS ML for each DeepSEA-based and Sei-based model and effect size from our caQTL study.

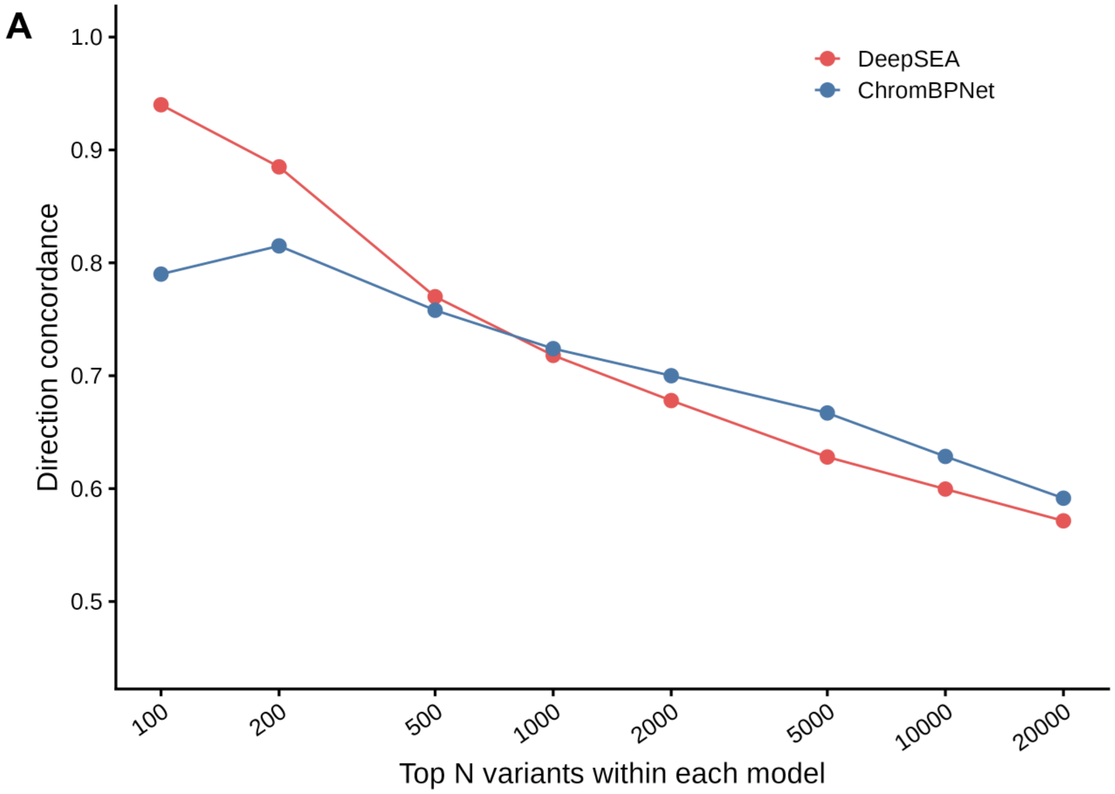


**Supplementary Fig. 5 | Comparison of a ChromBPNet-based prediction model. A,** Direction concordance between effect sizes from the previous caQTL study and *in silico* mutation effects from DeepSEA and ChromBPNet for each top N variants.

### Supplementary Tables

Supplementary Table 1 | The summary of 29 immune cell types (categorized into 8 subgroups).

| officail name | name | subgroup | NumSamples | NumSamples(QCed) | marker |
|---|---|---|---|---|---|
| Naïve CD4 T cells | Naive CD4 | CD4 | 52 | 46 | CD3+/CD4+CD8-/CCR7+CD45RA+ |
| T helper 1 cells | Th1 | CD4 | 53 | 48 | CD3+/CD4+CD8-/non-naive CD4+/CD25-/CXCR5-CCR6-/CXCR3+CCR4- |
| T helper 2 cells | Th2 | CD4 | 53 | 47 | CD3+/CD4+CD8-/non-naive CD4+/CD25-/CXCR5-CCR6-/CXCR3-CCR4+ |
| T helper 17 cells | Th17 | CD4 | 54 | 49 | CD3+/CD4+CD8-/non-naive CD4+/CD25-/CXCR5-CCR6+/CXCR3- |
| T follicular helper cells | Tfh | CD4 | 54 | 49 | CD3+/CD4+CD8-/non-naive CD4+/CD25-/CXCR5+ |
| Fraction I naïve regulatory T cells | FraI | CD4 | 52 | 46 | CD3+/CD4+CD8-/CD25+CD45RA+ |
| Fraction II effector regulatory T cells | FraII (aTreg) | CD4 | 53 | 50 | CD3+/CD4+CD8-/CD25++CD45RA- |
| Fraction III non-regulatory T cells | FraIII | CD4 | 53 | 48 | CD3+/CD4+CD8-/CD25lowCD45RA- |
| LAG3+ regulatory T cell | LAG3 Treg | CD4 | 52 | 48 | CD3+/CD4+CD8-/CD25-LAG3+/non-naïve CD4 |
| Memory CD4 T cells | Memory CD4 | CD4 | 54 | 48 | CD3+/CD4+CD8-/non-naive CD4+/CD25- |
| T helper non1/2 cells | Thx | CD4 | 54 | 48 | CD3+/CD4+CD8-/non-naive |

| | | | | | |
|---|---|---|---|---|---|
| | | | | | CD4+/CD25-/CXCR5-CCR6-/CXCR3-CCR4- |
| Naïve CD8 T cells | Naive CD8 | CD8 | 54 | 50 | CD3+CD19-/CD4-CD8+/CD45RA+CCR7+ |
| Effector Memory CD8 T cells | CD8+effector | CD8 | 53 | 47 | CD3+CD19-/CD4-CD8+/CD45RA+CCR7- |
| Central Memory CD8 T cells | CD8+central memory | CD8 | 52 | 45 | CD3+CD19-/CD4-CD8+/CD45RA-CCR7+ |
| CD8+ T effector memory CD45RA+ cells | CD8+effector memory | CD8 | 54 | 48 | CD3+CD19-/CD4-CD8+/CD45RA-CCR7- |
| Naïve B cells | Naive B | B | 54 | 48 | CD3-CD19+/IgD+CD27- |
| Unswitched memory B cells | unsw memory B | B | 53 | 49 | CD3-CD19+/IgD+CD27+ |
| Switched memory B cells | sw memory B | B | 52 | 48 | CD3-CD19+/IgD-CD27+/CD38- |
| Double Negative B cells | DN B | B | 52 | 49 | CD3-CD19+/IgD-CD27- |
| Plasmablasts | plasmablast | B | 52 | 43 | CD3-CD19+/IgD-CD27++/CD38+ |
| Plasmacytoid dendritic cells | plasmacytoid DC | DC | 52 | 44 | CD3-CD19-/HLARDR+/CD56-/CD14-CD16-/CD11c-CD123+ |
| Myeloid dendritic cells | myeloid DC | DC | 53 | 50 | CD3-CD19-/HLARDR+/CD56-/CD14-CD16-/CD11c+CD123- |
| CD16 positive monocytes | monocyte CD16+ | Mono | 53 | 48 | CD3-CD19-/HLARDR+/CD56-/CD14+CD16+ |
| Classical monocytes | monocyte CD16- | Mono | 53 | 49 | CD3-CD19-/HLARDR+/CD56-/CD14+CD16- |
| Non-classical monocytes | nonclassical Monocyte | Mono | 53 | 49 | CD3-CD19-/HLARDR+/CD56-/CD14dimCD16+ |
| Intermediate monocytes | intermediate Monocyte | Mono | 52 | 48 | CD3-CD19-/HLARDR+/CD56-/CD14++CD16+ |
| Natural Killer cells | NK | NK | 53 | 47 | CD3-CD19-/CD14-/CD56+ |
| peripheral blood | PBMC | PBMC | 53 | 47 | |

| | | | | |
|---|---|---|---|---|
| mononuclear cells | | | | |
| Neutrophil | Neutrophil | Neutrophil | 53 | 50 |

221

Supplementary Table 2 | The direction concordance between *in silico* mutation effect and caQTL effect size.

N_matched_var, Number of condordanct variant; N_total_var, Number of all the variant in this threshold; mutation_effect, threshold of *in silico* mutation effects to calculate direction concordance.

| name | subgroup | N_matched_var | N_total_var | direction_concordance | Pearson_corr | mutation_effect |
|---|---|---|---|---|---|---|
| Naive CD4 | CD4 | 375 | 663 | 0.57 | 0.27 | all |
| Naive CD4 | CD4 | 128 | 185 | 0.69 | 0.48 | 0.01 |
| Naive CD4 | CD4 | 62 | 65 | 0.95 | 0.74 | 0.05 |
| Naive CD4 | CD4 | 35 | 35 | 1.00 | 0.84 | 0.1 |
| Naive CD4 | CD4 | 20 | 20 | 1.00 | 0.86 | 0.2 |
| Naive CD4 | CD4 | 11 | 11 | 1.00 | 0.93 | 0.3 |
| Naive CD8 | CD8 | 351 | 668 | 0.53 | 0.20 | all |
| Naive CD8 | CD8 | 101 | 162 | 0.62 | 0.38 | 0.01 |
| Naive CD8 | CD8 | 38 | 41 | 0.93 | 0.75 | 0.05 |
| Naive CD8 | CD8 | 19 | 20 | 0.95 | 0.92 | 0.1 |
| Naive CD8 | CD8 | 12 | 12 | 1.00 | 0.96 | 0.2 |
| Naive CD8 | CD8 | 7 | 7 | 1.00 | 0.96 | 0.3 |
| Naive B | B | 365 | 700 | 0.52 | 0.19 | all |
| Naive B | B | 126 | 209 | 0.60 | 0.34 | 0.01 |
| Naive B | B | 41 | 47 | 0.87 | 0.69 | 0.05 |
| Naive B | B | 24 | 26 | 0.92 | 0.76 | 0.1 |
| Naive B | B | 11 | 11 | 1.00 | 0.90 | 0.2 |
| plasmacytoid DC | DC | 3195 | 5461 | 0.59 | 0.38 | all |
| plasmacytoid DC | DC | 1493 | 1998 | 0.75 | 0.56 | 0.01 |
| plasmacytoid DC | DC | 843 | 944 | 0.89 | 0.73 | 0.05 |
| plasmacytoid DC | DC | 563 | 599 | 0.94 | 0.81 | 0.1 |
| plasmacytoid DC | DC | 300 | 305 | 0.98 | 0.89 | 0.2 |
| plasmacytoid DC | DC | 181 | 183 | 0.99 | 0.92 | 0.3 |

| | | | | | | |
|---|---|---|---|---|---|---|
| monocyte CD16+ | Mono | 1872 | 3160 | 0.59 | 0.35 | all |
| monocyte CD16+ | Mono | 831 | 1122 | 0.74 | 0.55 | 0.01 |
| monocyte CD16+ | Mono | 498 | 561 | 0.89 | 0.71 | 0.05 |
| monocyte CD16+ | Mono | 352 | 371 | 0.95 | 0.81 | 0.1 |
| monocyte CD16+ | Mono | 194 | 199 | 0.97 | 0.88 | 0.2 |
| monocyte CD16+ | Mono | 113 | 115 | 0.98 | 0.91 | 0.3 |
| NK | NK | 1335 | 2392 | 0.56 | 0.30 | all |
| NK | NK | 462 | 630 | 0.73 | 0.51 | 0.01 |
| NK | NK | 217 | 248 | 0.88 | 0.72 | 0.05 |
| NK | NK | 148 | 158 | 0.94 | 0.80 | 0.1 |
| NK | NK | 66 | 67 | 0.99 | 0.90 | 0.2 |
| NK | NK | 35 | 35 | 1.00 | 0.93 | 0.3 |
| PBMC | PBMC | 346 | 673 | 0.51 | 0.17 | all |
| PBMC | PBMC | 93 | 153 | 0.61 | 0.36 | 0.01 |
| PBMC | PBMC | 35 | 39 | 0.90 | 0.68 | 0.05 |
| PBMC | PBMC | 21 | 21 | 1.00 | 0.88 | 0.1 |
| PBMC | PBMC | 10 | 10 | 1.00 | 0.93 | 0.2 |
| Neutrophil | Neutrophil | 619 | 1222 | 0.51 | 0.02 | all |
| Neutrophil | Neutrophil | 242 | 454 | 0.53 | 0.04 | 0.01 |
| Neutrophil | Neutrophil | 29 | 46 | 0.63 | 0.18 | 0.05 |
| Neutrophil | Neutrophil | 3 | 7 | 0.43 | -0.25 | 0.1 |

223 Supplementary Table 3 | The mean difference of seven known constraint metrics between cOCRs and CAMBUS control.

| name | subgroup | constraint_metric | P |
|---|---|---|---|
| Naive CD4 | CD4 | Gnocchi | 2.50.E-69 |
| Naive CD4 | CD4 | phyloP46way | 1.92.E-06 |
| Naive CD4 | CD4 | phyloP100way | 9.94.E-01 |
| Naive CD4 | CD4 | phyloPPrimates | 6.61.E-55 |
| Naive CD4 | CD4 | cactus241way | 2.03.E-20 |
| Naive CD4 | CD4 | DR | 1.14.E-10 |
| Naive CD4 | CD4 | GERP | 1.00.E+00 |
| Th1 | CD4 | Gnocchi | 8.80.E-77 |
| Th1 | CD4 | phyloP46way | 1.83.E-18 |
| Th1 | CD4 | phyloP100way | 2.20.E-03 |
| Th1 | CD4 | phyloPPrimates | 4.51.E-92 |
| Th1 | CD4 | cactus241way | 2.47.E-39 |
| Th1 | CD4 | DR | 1.49.E-12 |
| Th1 | CD4 | GERP | 1.00.E+00 |
| Th2 | CD4 | Gnocchi | 3.36.E-91 |
| Th2 | CD4 | phyloP46way | 1.37.E-23 |
| Th2 | CD4 | phyloP100way | 1.13.E-02 |
| Th2 | CD4 | phyloPPrimates | 1.07.E-106 |
| Th2 | CD4 | cactus241way | 1.65.E-48 |
| Th2 | CD4 | DR | 5.27.E-15 |
| Th2 | CD4 | GERP | 1.00.E+00 |
| Th17 | CD4 | Gnocchi | 3.55.E-106 |
| Th17 | CD4 | phyloP46way | 1.60.E-25 |
| Th17 | CD4 | phyloP100way | 6.09.E-04 |
| Th17 | CD4 | phyloPPrimates | 6.49.E-126 |
| Th17 | CD4 | cactus241way | 6.34.E-53 |
| Th17 | CD4 | DR | 6.40.E-21 |
| Th17 | CD4 | GERP | 1.00.E+00 |
| Tfh | CD4 | Gnocchi | 8.84.E-68 |
| Tfh | CD4 | phyloP46way | 3.08.E-17 |
| Tfh | CD4 | phyloP100way | 7.99.E-02 |
| Tfh | CD4 | phyloPPrimates | 5.50.E-74 |
| Tfh | CD4 | cactus241way | 8.99.E-31 |
| Tfh | CD4 | DR | 6.56.E-13 |
| Tfh | CD4 | GERP | 1.00.E+00 |
| FraI | CD4 | Gnocchi | 1.20.E-66 |

| | | | |
|---|---|---|---|
| FraI | CD4 | phyloP46way | 1.80.E-11 |
| FraI | CD4 | phyloP100way | 6.00.E-01 |
| FraI | CD4 | phyloPPrimates | 1.67.E-59 |
| FraI | CD4 | cactus241way | 4.94.E-28 |
| FraI | CD4 | DR | 1.93.E-12 |
| FraI | CD4 | GERP | 1.00.E+00 |
| FraII (aTreg) | CD4 | Gnocchi | 6.22.E-62 |
| FraII (aTreg) | CD4 | phyloP46way | 1.02.E-12 |
| FraII (aTreg) | CD4 | phyloP100way | 5.89.E-01 |
| FraII (aTreg) | CD4 | phyloPPrimates | 1.36.E-58 |
| FraII (aTreg) | CD4 | cactus241way | 1.28.E-25 |
| FraII (aTreg) | CD4 | DR | 1.57.E-10 |
| FraII (aTreg) | CD4 | GERP | 1.00.E+00 |
| FraIII | CD4 | Gnocchi | 1.83.E-77 |
| FraIII | CD4 | phyloP46way | 7.04.E-20 |
| FraIII | CD4 | phyloP100way | 1.88.E-02 |
| FraIII | CD4 | phyloPPrimates | 1.17.E-92 |
| FraIII | CD4 | cactus241way | 1.71.E-35 |
| FraIII | CD4 | DR | 4.62.E-15 |
| FraIII | CD4 | GERP | 1.00.E+00 |
| LAG3 Treg | CD4 | Gnocchi | 2.20.E-82 |
| LAG3 Treg | CD4 | phyloP46way | 3.38.E-24 |
| LAG3 Treg | CD4 | phyloP100way | 2.66.E-04 |
| LAG3 Treg | CD4 | phyloPPrimates | 8.00.E-107 |
| LAG3 Treg | CD4 | cactus241way | 4.36.E-45 |
| LAG3 Treg | CD4 | DR | 5.06.E-12 |
| LAG3 Treg | CD4 | GERP | 1.00.E+00 |
| Memory CD4 | CD4 | Gnocchi | 4.45.E-102 |
| Memory CD4 | CD4 | phyloP46way | 1.07.E-19 |
| Memory CD4 | CD4 | phyloP100way | 2.79.E-02 |
| Memory CD4 | CD4 | phyloPPrimates | 2.69.E-107 |
| Memory CD4 | CD4 | cactus241way | 5.55.E-46 |
| Memory CD4 | CD4 | DR | 2.69.E-16 |
| Memory CD4 | CD4 | GERP | 1.00.E+00 |
| Thx | CD4 | Gnocchi | 1.26.E-85 |
| Thx | CD4 | phyloP46way | 3.65.E-20 |
| Thx | CD4 | phyloP100way | 3.54.E-03 |
| Thx | CD4 | phyloPPrimates | 2.88.E-101 |
| Thx | CD4 | cactus241way | 1.89.E-40 |
| Thx | CD4 | DR | 1.73.E-12 |
| Thx | CD4 | GERP | 1.00.E+00 |

| | | | |
|---|---|---|---|
| Naive CD8 | CD8 | Gnocchi | 1.26.E-20 |
| Naive CD8 | CD8 | phyloP46way | 4.86.E-02 |
| Naive CD8 | CD8 | phyloP100way | 9.99.E-01 |
| Naive CD8 | CD8 | phyloPPrimates | 6.70.E-14 |
| Naive CD8 | CD8 | cactus241way | 2.31.E-04 |
| Naive CD8 | CD8 | DR | 6.18.E-05 |
| Naive CD8 | CD8 | GERP | 1.00.E+00 |
| CD8+effector | CD8 | Gnocchi | 1.87.E-44 |
| CD8+effector | CD8 | phyloP46way | 6.37.E-05 |
| CD8+effector | CD8 | phyloP100way | 9.62.E-01 |
| CD8+effector | CD8 | phyloPPrimates | 1.14.E-30 |
| CD8+effector | CD8 | cactus241way | 9.03.E-13 |
| CD8+effector | CD8 | DR | 1.77.E-08 |
| CD8+effector | CD8 | GERP | 1.00.E+00 |
| CD8+central memory | CD8 | Gnocchi | 3.37.E-58 |
| CD8+central memory | CD8 | phyloP46way | 7.46.E-08 |
| CD8+central memory | CD8 | phyloP100way | 8.02.E-01 |
| CD8+central memory | CD8 | phyloPPrimates | 2.37.E-52 |
| CD8+central memory | CD8 | cactus241way | 4.65.E-23 |
| CD8+central memory | CD8 | DR | 5.94.E-10 |
| CD8+central memory | CD8 | GERP | 1.00.E+00 |
| CD8+effector memory | CD8 | Gnocchi | 7.61.E-40 |
| CD8+effector memory | CD8 | phyloP46way | 6.62.E-06 |
| CD8+effector memory | CD8 | phyloP100way | 9.14.E-01 |
| CD8+effector memory | CD8 | phyloPPrimates | 7.10.E-32 |
| CD8+effector memory | CD8 | cactus241way | 1.07.E-13 |
| CD8+effector memory | CD8 | DR | 1.19.E-08 |
| CD8+effector memory | CD8 | GERP | 1.00.E+00 |
| Naive B | B | Gnocchi | 2.61.E-18 |
| Naive B | B | phyloP46way | 3.97.E-01 |
| Naive B | B | phyloP100way | 9.95.E-01 |
| Naive B | B | phyloPPrimates | 2.93.E-08 |
| Naive B | B | cactus241way | 4.13.E-04 |
| Naive B | B | DR | 3.50.E-02 |
| Naive B | B | GERP | 1.00.E+00 |
| unsw memory B | B | Gnocchi | 7.17.E-62 |
| unsw memory B | B | phyloP46way | 9.34.E-05 |
| unsw memory B | B | phyloP100way | 9.99.E-01 |
| unsw memory B | B | phyloPPrimates | 1.54.E-35 |
| unsw memory B | B | cactus241way | 1.07.E-13 |
| unsw memory B | B | DR | 3.53.E-09 |
| unsw memory B | B | GERP | 1.00.E+00 |
| sw memory B | B | Gnocchi | 1.11.E-84 |
| sw memory B | B | phyloP46way | 8.95.E-05 |

| | | | |
|---|---|---|---|
| sw memory B | B | phyloP100way | 1.00.E+00 |
| sw memory B | B | phyloPPrimates | 1.46.E-40 |
| sw memory B | B | cactus241way | 4.68.E-16 |
| sw memory B | B | DR | 1.66.E-08 |
| sw memory B | B | GERP | 1.00.E+00 |
| DN B | B | Gnocchi | 1.86.E-113 |
| DN B | B | phyloP46way | 6.42.E-08 |
| DN B | B | phyloP100way | 1.00.E+00 |
| DN B | B | phyloPPrimates | 9.78.E-59 |
| DN B | B | cactus241way | 8.27.E-22 |
| DN B | B | DR | 5.63.E-12 |
| DN B | B | GERP | 1.00.E+00 |
| plasmablast | B | Gnocchi | 4.52.E-116 |
| plasmablast | B | phyloP46way | 3.56.E-06 |
| plasmablast | B | phyloP100way | 1.00.E+00 |
| plasmablast | B | phyloPPrimates | 1.62.E-83 |
| plasmablast | B | cactus241way | 3.43.E-30 |
| plasmablast | B | DR | 1.85.E-12 |
| plasmablast | B | GERP | 1.00.E+00 |
| plasmacytoid DC | DC | Gnocchi | 4.77.E-18 |
| plasmacytoid DC | DC | phyloP46way | 9.70.E-01 |
| plasmacytoid DC | DC | phyloP100way | 1.00.E+00 |
| plasmacytoid DC | DC | phyloPPrimates | 9.17.E-05 |
| plasmacytoid DC | DC | cactus241way | 7.46.E-02 |
| plasmacytoid DC | DC | DR | 3.06.E-01 |
| plasmacytoid DC | DC | GERP | 1.00.E+00 |
| myeloid DC | DC | Gnocchi | 1.78.E-14 |
| myeloid DC | DC | phyloP46way | 1.57.E-02 |
| myeloid DC | DC | phyloP100way | 4.64.E-01 |
| myeloid DC | DC | phyloPPrimates | 1.63.E-13 |
| myeloid DC | DC | cactus241way | 2.44.E-06 |
| myeloid DC | DC | DR | 9.83.E-03 |
| myeloid DC | DC | GERP | 1.00.E+00 |
| monocyte CD16+ | Mono | Gnocchi | 2.26.E-74 |
| monocyte CD16+ | Mono | phyloP46way | 1.94.E-24 |
| monocyte CD16+ | Mono | phyloP100way | 9.38.E-04 |
| monocyte CD16+ | Mono | phyloPPrimates | 1.63.E-100 |
| monocyte CD16+ | Mono | cactus241way | 6.16.E-52 |
| monocyte CD16+ | Mono | DR | 1.19.E-16 |
| monocyte CD16+ | Mono | GERP | 1.00.E+00 |
| monocyte CD16- | Mono | Gnocchi | 3.32.E-144 |

| | | | |
|---|---|---|---|
| monocyte CD16- | Mono | phyloP46way | 4.57.E-31 |
| monocyte CD16- | Mono | phyloP100way | 2.47.E-01 |
| monocyte CD16- | Mono | phyloPPrimates | 7.11.E-142 |
| monocyte CD16- | Mono | cactus241way | 4.48.E-70 |
| monocyte CD16- | Mono | DR | 1.98.E-32 |
| monocyte CD16- | Mono | GERP | 1.00.E+00 |
| nonclassical Monocyte | Mono | Gnocchi | 2.64.E-25 |
| nonclassical Monocyte | Mono | phyloP46way | 4.65.E-06 |
| nonclassical Monocyte | Mono | phyloP100way | 2.28.E-01 |
| nonclassical Monocyte | Mono | phyloPPrimates | 1.44.E-33 |
| nonclassical Monocyte | Mono | cactus241way | 2.85.E-15 |
| nonclassical Monocyte | Mono | DR | 6.21.E-05 |
| nonclassical Monocyte | Mono | GERP | 1.00.E+00 |
| intermediate Monocyte | Mono | Gnocchi | 1.57.E-137 |
| intermediate Monocyte | Mono | phyloP46way | 1.62.E-32 |
| intermediate Monocyte | Mono | phyloP100way | 1.80.E-01 |
| intermediate Monocyte | Mono | phyloPPrimates | 5.25.E-137 |
| intermediate Monocyte | Mono | cactus241way | 1.13.E-66 |
| intermediate Monocyte | Mono | DR | 8.65.E-30 |
| intermediate Monocyte | Mono | GERP | 1.00.E+00 |
| NK | NK | Gnocchi | 7.92.E-91 |
| NK | NK | phyloP46way | 8.65.E-12 |
| NK | NK | phyloP100way | 2.13.E-01 |
| NK | NK | phyloPPrimates | 6.92.E-74 |
| NK | NK | cactus241way | 2.81.E-31 |
| NK | NK | DR | 2.37.E-11 |
| NK | NK | GERP | 1.00.E+00 |
| PBMC | PBMC | Gnocchi | 0.00.E+00 |
| PBMC | PBMC | phyloP46way | 3.89.E-36 |
| PBMC | PBMC | phyloP100way | 9.99.E-01 |
| PBMC | PBMC | phyloPPrimates | 0.00.E+00 |
| PBMC | PBMC | cactus241way | 1.43.E-118 |
| PBMC | PBMC | DR | 3.23.E-50 |
| PBMC | PBMC | GERP | 1.00.E+00 |
| Neutrophil | Neutrophil | Gnocchi | 1.39.E-75 |
| Neutrophil | Neutrophil | phyloP46way | 2.53.E-08 |
| Neutrophil | Neutrophil | phyloP100way | 9.94.E-01 |
| Neutrophil | Neutrophil | phyloPPrimates | 1.46.E-47 |
| Neutrophil | Neutrophil | cactus241way | 5.52.E-27 |
| Neutrophil | Neutrophil | DR | 1.18.E-07 |
| Neutrophil | Neutrophil | GERP | 1.00.E+00 |

Supplementary Table 4 | Enrichment of cOCRs in NaiveCD4 comparing with the known cCREs using various Gnocchi.
OR, odds ratio; CI_lo, lower 95% confidence interval; CI_hi, upper 95% confidence interval.

| Gnocchi | name | OR | CI_lo | CI_hi | P |
|---|---|---|---|---|---|
| -5 | Naive CD4 | 0.31 | 0.16 | 0.59 | 1 |
| -4 | Naive CD4 | 0.38 | 0.21 | 0.69 | 1 |
| -3 | Naive CD4 | 0.11 | 0.05 | 0.24 | 1 |
| -2 | Naive CD4 | 0.3 | 0.21 | 0.42 | 1 |
| -1 | Naive CD4 | 0.42 | 0.33 | 0.54 | 1 |
| 0 | Naive CD4 | 0.59 | 0.49 | 0.72 | 1 |
| 1 | Naive CD4 | 1.23 | 1.05 | 1.45 | 0.0073 |
| 2 | Naive CD4 | 2.71 | 2.32 | 3.17 | 3.4E-29 |
| 3 | Naive CD4 | 5.5 | 4.57 | 6.61 | 1.1E-49 |
| 4 | Naive CD4 | 11.45 | 9.33 | 14.05 | 4.7E-68 |
| -5 | ENCODE cCRE-PLS | 0.55 | 0.49 | 0.62 | 1 |
| -4 | ENCODE cCRE-PLS | 0.5 | 0.44 | 0.57 | 1 |
| -3 | ENCODE cCRE-PLS | 0.42 | 0.38 | 0.46 | 1 |
| -2 | ENCODE cCRE-PLS | 0.51 | 0.48 | 0.55 | 1 |
| -1 | ENCODE cCRE-PLS | 0.62 | 0.59 | 0.65 | 1 |
| 0 | ENCODE cCRE-PLS | 0.88 | 0.84 | 0.91 | 1 |
| 1 | ENCODE cCRE-PLS | 1.25 | 1.2 | 1.31 | 2.3E-27 |
| 2 | ENCODE cCRE-PLS | 1.97 | 1.88 | 2.06 | 7.7E-163 |
| 3 | ENCODE cCRE-PLS | 3.23 | 3.05 | 3.43 | 7.4E-244 |
| 4 | ENCODE cCRE-PLS | 7.28 | 6.82 | 7.78 | <2.2E-308 |
| -5 | ENCODE cCRE-pELS | 0.37 | 0.35 | 0.4 | 1 |
| -4 | ENCODE cCRE-pELS | 0.37 | 0.35 | 0.4 | 1 |
| -3 | ENCODE cCRE-pELS | 0.43 | 0.41 | 0.45 | 1 |
| -2 | ENCODE cCRE-pELS | 0.54 | 0.52 | 0.56 | 1 |
| -1 | ENCODE cCRE-pELS | 0.72 | 0.7 | 0.74 | 1 |
| 0 | ENCODE cCRE-pELS | 0.98 | 0.96 | 1 | 0.98 |
| 1 | ENCODE cCRE-pELS | 1.37 | 1.35 | 1.4 | 1.2E-208 |
| 2 | ENCODE cCRE-pELS | 1.92 | 1.87 | 1.96 | <2.2E-308 |
| 3 | ENCODE cCRE-pELS | 2.65 | 2.56 | 2.74 | <2.2E-308 |
| 4 | ENCODE cCRE-pELS | 4.35 | 4.16 | 4.54 | <2.2E-308 |
| -5 | ENCODE cCRE-dELS | 0.3 | 0.3 | 0.31 | 1 |
| -4 | ENCODE cCRE-dELS | 0.41 | 0.4 | 0.42 | 1 |
| -3 | ENCODE cCRE-dELS | 0.51 | 0.5 | 0.52 | 1 |
| -2 | ENCODE cCRE-dELS | 0.66 | 0.65 | 0.67 | 1 |
| -1 | ENCODE cCRE-dELS | 0.86 | 0.86 | 0.87 | 1 |
| 0 | ENCODE cCRE-dELS | 1.12 | 1.11 | 1.13 | 5.4E-174 |
| 1 | ENCODE cCRE-dELS | 1.4 | 1.39 | 1.42 | <2.2E-308 |
| 2 | ENCODE cCRE-dELS | 1.68 | 1.66 | 1.7 | <2.2E-308 |
| 3 | ENCODE cCRE-dELS | 1.98 | 1.94 | 2.01 | <2.2E-308 |
| 4 | ENCODE cCRE-dELS | 2.14 | 2.08 | 2.2 | <2.2E-308 |
| -5 | ENCODE CTCF-only | 0.7 | 0.66 | 0.74 | 1 |

| | | | | | |
|---|---|---|---|---|---|
| -4 | ENCODE CTCF-only | 0.89 | 0.84 | 0.93 | 1 |
| -3 | ENCODE CTCF-only | 0.98 | 0.94 | 1.01 | 0.92 |
| -2 | ENCODE CTCF-only | 1.04 | 1.02 | 1.07 | 0.00069 |
| -1 | ENCODE CTCF-only | 1.04 | 1.01 | 1.06 | 0.00053 |
| 0 | ENCODE CTCF-only | 1.03 | 1.01 | 1.05 | 0.0023 |
| 1 | ENCODE CTCF-only | 1 | 0.97 | 1.02 | 0.65 |
| 2 | ENCODE CTCF-only | 1 | 0.97 | 1.04 | 0.41 |
| 3 | ENCODE CTCF-only | 0.94 | 0.89 | 0.99 | 0.99 |
| 4 | ENCODE CTCF-only | 0.82 | 0.75 | 0.9 | 1 |
| -5 | Super enhancers | 0.28 | 0.27 | 0.28 | 1 |
| -4 | Super enhancers | 0.34 | 0.34 | 0.35 | 1 |
| -3 | Super enhancers | 0.44 | 0.43 | 0.44 | 1 |
| -2 | Super enhancers | 0.6 | 0.59 | 0.6 | 1 |
| -1 | Super enhancers | 0.83 | 0.83 | 0.84 | 1 |
| 0 | Super enhancers | 1.16 | 1.16 | 1.17 | <2.2E-308 |
| 1 | Super enhancers | 1.58 | 1.57 | 1.59 | <2.2E-308 |
| 2 | Super enhancers | 2.05 | 2.03 | 2.07 | <2.2E-308 |
| 3 | Super enhancers | 2.63 | 2.58 | 2.68 | <2.2E-308 |
| 4 | Super enhancers | 3.42 | 3.31 | 3.52 | <2.2E-308 |
| -5 | FANTOM enhancers | 0.33 | 0.3 | 0.36 | 1 |
| -4 | FANTOM enhancers | 0.4 | 0.37 | 0.43 | 1 |
| -3 | FANTOM enhancers | 0.45 | 0.43 | 0.48 | 1 |
| -2 | FANTOM enhancers | 0.58 | 0.56 | 0.6 | 1 |
| -1 | FANTOM enhancers | 0.76 | 0.74 | 0.78 | 1 |
| 0 | FANTOM enhancers | 1.01 | 0.99 | 1.03 | 0.17 |
| 1 | FANTOM enhancers | 1.34 | 1.32 | 1.37 | 1.5E-168 |
| 2 | FANTOM enhancers | 1.82 | 1.77 | 1.86 | <2.2E-308 |
| 3 | FANTOM enhancers | 2.52 | 2.44 | 2.61 | <2.2E-308 |
| 4 | FANTOM enhancers | 3.58 | 3.41 | 3.76 | <2.2E-308 |

Supplementary Table 5 | The enrichment analysis for archaic genomes from 1000 genomes in cOCRs.
POP, population; OR, odds ratio; CI_lo, lower 95% confidence interval; CI_hi, upper 95% confidence interval.

| origin | POP | OR | CI_lo | CI_hi | P |
|---|---|---|---|---|---|
| Neanderthal | AMR | 1.99 | 1.89 | 2.09 | 9.75.E-157 |
| Neanderthal | EAS | 1.94 | 1.86 | 2.02 | 4.93.E-226 |
| Neanderthal | EUR | 2.01 | 1.92 | 2.1 | 1.50.E-207 |
| Neanderthal | SAS | 1.94 | 1.88 | 2 | 2.74.E-340 |
| Denisovans | AMR | 1.99 | 1.86 | 2.12 | 1.35.E-90 |
| Denisovans | EAS | 1.83 | 1.72 | 1.94 | 5.65.E-88 |
| Denisovans | EUR | 2.13 | 2 | 2.27 | 7.59.E-120 |
| Denisovans | SAS | 2.24 | 2.12 | 2.37 | 1.07.E-167 |

Supplementary Table 6 | The enrichment analysis for HGMD in cOCRs. OR, odds ratio; CI_lo, lower 95% confidence interval; CI_hi, upper 95% confidence interval.

| name | subgroup | OR | ci_lo | ci_hi | P |
|---|---|---|---|---|---|
| Naive CD4 | CD4 | 2.80 | 1.76 | 4.48 | 1.46.E-04 |
| Th1 | CD4 | 2.01 | 1.38 | 2.94 | 6.53.E-04 |
| Th2 | CD4 | 2.97 | 2.12 | 4.17 | 4.71.E-08 |
| Th17 | CD4 | 2.45 | 1.85 | 3.24 | 2.04.E-08 |
| Tfh | CD4 | 2.15 | 1.35 | 3.44 | 2.74.E-03 |
| FraI | CD4 | 2.43 | 1.5 | 3.93 | 1.06.E-03 |
| FraII (aTreg) | CD4 | 2.55 | 1.65 | 3.94 | 1.58.E-04 |
| FraIII | CD4 | 1.98 | 1.46 | 2.7 | 5.41.E-05 |
| LAG3 Treg | CD4 | 1.77 | 1.32 | 2.38 | 3.11.E-04 |
| Memory CD4 | CD4 | 2.86 | 2.06 | 3.98 | 4.96.E-08 |
| Thx | CD4 | 2.16 | 1.59 | 2.93 | 6.96.E-06 |
| Naive CD8 | CD8 | 3.04 | 1.13 | 8.18 | 4.55.E-02 |
| CD8+effector | CD8 | 3.06 | 1.72 | 5.42 | 8.45.E-04 |
| CD8+central memory | CD8 | 2.09 | 1.18 | 3.69 | 1.57.E-02 |
| CD8+effector memory | CD8 | 3.13 | 1.72 | 5.69 | 1.14.E-03 |
| Naive B | B | 4.40 | 1.82 | 10.69 | 6.46.E-03 |
| unsw memory B | B | 3.20 | 1.76 | 5.82 | 9.34.E-04 |
| sw memory B | B | 4.20 | 2.69 | 6.56 | 1.96.E-07 |
| DN B | B | 3.45 | 2.26 | 5.28 | 1.24.E-06 |
| plasmablast | B | 2.70 | 2.02 | 3.61 | 3.83.E-09 |
| plasmacytoid DC | DC | 1.98 | 1.12 | 3.51 | 2.24.E-02 |
| myeloid DC | DC | 1.78 | 0.98 | 3.24 | 5.15.E-02 |
| monocyte CD16+ | Mono | 2.55 | 1.95 | 3.35 | 2.04.E-09 |
| monocyte CD16- | Mono | 3.41 | 2.52 | 4.61 | 1.63.E-11 |
| nonclassical Monocyte | Mono | 2.33 | 1.65 | 3.28 | 1.31.E-05 |
| intermediate Monocyte | Mono | 3.43 | 2.61 | 4.5 | 4.89.E-14 |
| NK | NK | 2.10 | 1.48 | 2.99 | 1.43.E-04 |
| PBMC | PBMC | 3.28 | 2.76 | 3.89 | 6.57.E-32 |
| Neutrophil | Neutrophil | 6.53 | 4.27 | 9.99 | 1.92.E-11 |

| pooled OR | CI_lo | CI_hi | P |
|---|---|---|---|
| 2.70 | 2.43 | 2.99 | 1.71e-77 |

Supplementary Table 7 | Number of shared super enhancer/FANTOM5 enancers with cOCRs and all OCRs.
Total, Number of OCRs; cCRE+, Number of OCRs overlapped with cCRE; cCRE+ & cOCRs, Number of cOCRs overlapped with cCRE

| name | subgroup | cCRE | Total | cCRE+ | cCRE+ & cOCRs |
|---|---|---|---|---|---|
| Naive CD4 | CD4 | ENCODE cCRE-PLS | 254,701 | 17,315 | 302 |
| Naive CD4 | CD4 | ENCODE cCRE-pELS | 254,701 | 25,146 | 433 |
| Naive CD4 | CD4 | ENCODE cCRE-dELS | 254,701 | 66,686 | 571 |
| Naive CD4 | CD4 | ENCODE CTCF-only | 254,701 | 3,874 | 6 |
| Naive CD4 | CD4 | FANTOM enhancers | 254,701 | 11,430 | 284 |
| Naive CD4 | CD4 | Super enhancers | 254,701 | 187,605 | 1,218 |
| Th1 | CD4 | ENCODE cCRE-PLS | 220,655 | 17,444 | 581 |
| Th1 | CD4 | ENCODE cCRE-pELS | 220,655 | 25,561 | 809 |
| Th1 | CD4 | ENCODE cCRE-dELS | 220,655 | 66,995 | 1,282 |
| Th1 | CD4 | ENCODE CTCF-only | 220,655 | 3,952 | 14 |
| Th1 | CD4 | FANTOM enhancers | 220,655 | 12,608 | 614 |
| Th1 | CD4 | Super enhancers | 220,655 | 169,900 | 2,313 |
| Th2 | CD4 | ENCODE cCRE-PLS | 252,630 | 17,668 | 549 |
| Th2 | CD4 | ENCODE cCRE-pELS | 252,630 | 25,851 | 768 |
| Th2 | CD4 | ENCODE cCRE-dELS | 252,630 | 70,101 | 1,149 |
| Th2 | CD4 | ENCODE CTCF-only | 252,630 | 4,321 | 14 |
| Th2 | CD4 | FANTOM enhancers | 252,630 | 12,425 | 548 |
| Th2 | CD4 | Super enhancers | 252,630 | 188,707 | 2,194 |
| Th17 | CD4 | ENCODE cCRE-PLS | 231,143 | 17,767 | 857 |
| Th17 | CD4 | ENCODE cCRE-pELS | 231,143 | 25,913 | 1,203 |
| Th17 | CD4 | ENCODE cCRE-dELS | 231,143 | 70,767 | 2,085 |
| Th17 | CD4 | ENCODE CTCF-only | 231,143 | 4,554 | 43 |
| Th17 | CD4 | FANTOM enhancers | 231,143 | 13,266 | 901 |

| | | | | | |
|---|---|---|---|---|---|
| Th17 | CD4 | Super enhancers | 231,143 | 175,664 | 3,635 |
| Tfh | CD4 | ENCODE cCRE-PLS | 251,684 | 17,679 | 387 |
| Tfh | CD4 | ENCODE cCRE-pELS | 251,684 | 25,906 | 543 |
| Tfh | CD4 | ENCODE cCRE-dELS | 251,684 | 69,612 | 781 |
| Tfh | CD4 | ENCODE CTCF-only | 251,684 | 4,389 | 8 |
| Tfh | CD4 | FANTOM enhancers | 251,684 | 12,380 | 398 |
| Tfh | CD4 | Super enhancers | 251,684 | 187,502 | 1,519 |
| FraI | CD4 | ENCODE cCRE-PLS | 239,064 | 17,288 | 309 |
| FraI | CD4 | ENCODE cCRE-pELS | 239,064 | 25,017 | 439 |
| FraI | CD4 | ENCODE cCRE-dELS | 239,064 | 64,870 | 609 |
| FraI | CD4 | ENCODE CTCF-only | 239,064 | 3,780 | 7 |
| FraI | CD4 | FANTOM enhancers | 239,064 | 11,567 | 312 |
| FraI | CD4 | Super enhancers | 239,064 | 178,586 | 1,261 |
| FraII (aTreg) | CD4 | ENCODE cCRE-PLS | 239,027 | 17,412 | 356 |
| FraII (aTreg) | CD4 | ENCODE cCRE-pELS | 239,027 | 25,339 | 508 |
| FraII (aTreg) | CD4 | ENCODE cCRE-dELS | 239,027 | 66,448 | 735 |
| FraII (aTreg) | CD4 | ENCODE CTCF-only | 239,027 | 3,964 | 7 |
| FraII (aTreg) | CD4 | FANTOM enhancers | 239,027 | 12,117 | 366 |
| FraII (aTreg) | CD4 | Super enhancers | 239,027 | 180,828 | 1,465 |
| FraIII | CD4 | ENCODE cCRE-PLS | 202,271 | 17,542 | 766 |
| FraIII | CD4 | ENCODE cCRE-pELS | 202,271 | 25,473 | 1,058 |
| FraIII | CD4 | ENCODE cCRE-dELS | 202,271 | 65,635 | 1,883 |
| FraIII | CD4 | ENCODE CTCF-only | 202,271 | 4,058 | 28 |
| FraIII | CD4 | FANTOM enhancers | 202,271 | 12,848 | 850 |
| FraIII | CD4 | Super enhancers | 202,271 | 158,205 | 3,238 |
| LAG3 Treg | CD4 | ENCODE cCRE-PLS | 198,977 | 17,535 | 894 |
| LAG3 Treg | CD4 | ENCODE cCRE-pELS | 198,977 | 25,442 | 1,254 |
| LAG3 Treg | CD4 | ENCODE cCRE-dELS | 198,977 | 65,283 | 2,263 |
| LAG3 Treg | CD4 | ENCODE CTCF-only | 198,977 | 4,094 | 39 |
| LAG3 Treg | CD4 | FANTOM enhancers | 198,977 | 12,817 | 941 |

| | | | | | |
|---|---|---|---|---|---|
| LAG3 Treg | CD4 | Super enhancers | 198,977 | 157,045 | 3,854 |
| Memory CD4 | CD4 | ENCODE cCRE-PLS | 249,443 | 17,549 | 598 |
| Memory CD4 | CD4 | ENCODE cCRE-pELS | 249,443 | 25,727 | 849 |
| Memory CD4 | CD4 | ENCODE cCRE-dELS | 249,443 | 70,511 | 1,253 |
| Memory CD4 | CD4 | ENCODE CTCF-only | 249,443 | 4,178 | 17 |
| Memory CD4 | CD4 | FANTOM enhancers | 249,443 | 12,715 | 597 |
| Memory CD4 | CD4 | Super enhancers | 249,443 | 186,524 | 2,384 |
| Thx | CD4 | ENCODE cCRE-PLS | 211,500 | 17,234 | 766 |
| Thx | CD4 | ENCODE cCRE-pELS | 211,500 | 25,209 | 1,066 |
| Thx | CD4 | ENCODE cCRE-dELS | 211,500 | 67,083 | 1,853 |
| Thx | CD4 | ENCODE CTCF-only | 211,500 | 4,044 | 29 |
| Thx | CD4 | FANTOM enhancers | 211,500 | 12,873 | 826 |
| Thx | CD4 | Super enhancers | 211,500 | 165,077 | 3,239 |
| Naive CD8 | CD8 | ENCODE cCRE-PLS | 342,178 | 18,261 | 50 |
| Naive CD8 | CD8 | ENCODE cCRE-pELS | 342,178 | 27,562 | 70 |
| Naive CD8 | CD8 | ENCODE cCRE-dELS | 342,178 | 83,687 | 89 |
| Naive CD8 | CD8 | ENCODE CTCF-only | 342,178 | 4,776 | 0 |
| Naive CD8 | CD8 | FANTOM enhancers | 342,178 | 12,848 | 45 |
| Naive CD8 | CD8 | Super enhancers | 342,178 | 245,722 | 295 |
| CD8+effector | CD8 | ENCODE cCRE-PLS | 288,600 | 17,699 | 186 |
| CD8+effector | CD8 | ENCODE cCRE-pELS | 288,600 | 26,348 | 261 |
| CD8+effector | CD8 | ENCODE cCRE-dELS | 288,600 | 77,290 | 369 |
| CD8+effector | CD8 | ENCODE CTCF-only | 288,600 | 4,469 | 2 |
| CD8+effector | CD8 | FANTOM enhancers | 288,600 | 13,194 | 210 |
| CD8+effector | CD8 | Super enhancers | 288,600 | 212,446 | 790 |
| CD8+central memory | CD8 | ENCODE cCRE-PLS | 266,387 | 17,613 | 267 |
| CD8+central memory | CD8 | ENCODE cCRE-pELS | 266,387 | 25,679 | 385 |
| CD8+central memory | CD8 | ENCODE cCRE-dELS | 266,387 | 70,869 | 549 |
| CD8+central memory | CD8 | ENCODE CTCF-only | 266,387 | 4,006 | 2 |
| CD8+central memory | CD8 | FANTOM enhancers | 266,387 | 12,413 | 277 |

| | | | | | |
|---|---|---|---|---|---|
| CD8+central memory | CD8 | Super enhancers | 266,387 | 196,695 | 1,117 |
| CD8+effector memory | CD8 | ENCODE cCRE-PLS | 285,942 | 17,684 | 174 |
| CD8+effector memory | CD8 | ENCODE cCRE-pELS | 285,942 | 26,278 | 240 |
| CD8+effector memory | CD8 | ENCODE cCRE-dELS | 285,942 | 75,112 | 314 |
| CD8+effector memory | CD8 | ENCODE CTCF-only | 285,942 | 4,206 | 1 |
| CD8+effector memory | CD8 | FANTOM enhancers | 285,942 | 12,975 | 176 |
| CD8+effector memory | CD8 | Super enhancers | 285,942 | 211,524 | 707 |
| Naive B | B | ENCODE cCRE-PLS | 357,770 | 18,396 | 41 |
| Naive B | B | ENCODE cCRE-pELS | 357,770 | 28,100 | 58 |
| Naive B | B | ENCODE cCRE-dELS | 357,770 | 88,532 | 64 |
| Naive B | B | ENCODE CTCF-only | 357,770 | 4,777 | 1 |
| Naive B | B | FANTOM enhancers | 357,770 | 13,722 | 41 |
| Naive B | B | Super enhancers | 357,770 | 256,176 | 267 |
| unsw memory B | B | ENCODE cCRE-PLS | 326,961 | 18,135 | 162 |
| unsw memory B | B | ENCODE cCRE-pELS | 326,961 | 27,738 | 240 |
| unsw memory B | B | ENCODE cCRE-dELS | 326,961 | 86,669 | 332 |
| unsw memory B | B | ENCODE CTCF-only | 326,961 | 4,685 | 2 |
| unsw memory B | B | FANTOM enhancers | 326,961 | 14,167 | 158 |
| unsw memory B | B | Super enhancers | 326,961 | 234,873 | 758 |
| sw memory B | B | ENCODE cCRE-PLS | 329,416 | 18,186 | 250 |
| sw memory B | B | ENCODE cCRE-pELS | 329,416 | 28,247 | 346 |
| sw memory B | B | ENCODE cCRE-dELS | 329,416 | 88,952 | 474 |
| sw memory B | B | ENCODE CTCF-only | 329,416 | 4,744 | 2 |
| sw memory B | B | FANTOM enhancers | 329,416 | 14,624 | 229 |
| sw memory B | B | Super enhancers | 329,416 | 241,018 | 1,024 |
| DN B | B | ENCODE cCRE-PLS | 323,974 | 18,126 | 333 |
| DN B | B | ENCODE cCRE-pELS | 323,974 | 27,647 | 475 |
| DN B | B | ENCODE cCRE-dELS | 323,974 | 86,834 | 666 |
| DN B | B | ENCODE CTCF-only | 323,974 | 4,599 | 5 |
| DN B | B | FANTOM enhancers | 323,974 | 14,440 | 301 |

| | | | | | |
|---|---|---|---|---|---|
| DN B | B | Super enhancers | 323,974 | 234,959 | 1,383 |
| plasmablast | B | ENCODE cCRE-PLS | 225,711 | 17,167 | 743 |
| plasmablast | B | ENCODE cCRE-pELS | 225,711 | 25,368 | 1,053 |
| plasmablast | B | ENCODE cCRE-dELS | 225,711 | 73,534 | 1,602 |
| plasmablast | B | ENCODE CTCF-only | 225,711 | 4,194 | 34 |
| plasmablast | B | FANTOM enhancers | 225,711 | 12,622 | 627 |
| plasmablast | B | Super enhancers | 225,711 | 177,343 | 3,007 |
| plasmacytoid DC | DC | ENCODE cCRE-PLS | 208,009 | 17,971 | 183 |
| plasmacytoid DC | DC | ENCODE cCRE-pELS | 208,009 | 28,760 | 252 |
| plasmacytoid DC | DC | ENCODE cCRE-dELS | 208,009 | 88,586 | 575 |
| plasmacytoid DC | DC | ENCODE CTCF-only | 208,009 | 5,154 | 9 |
| plasmacytoid DC | DC | FANTOM enhancers | 208,009 | 16,174 | 252 |
| plasmacytoid DC | DC | Super enhancers | 208,009 | 173,263 | 913 |
| myeloid DC | DC | ENCODE cCRE-PLS | 240,586 | 18,516 | 180 |
| myeloid DC | DC | ENCODE cCRE-pELS | 240,586 | 29,958 | 263 |
| myeloid DC | DC | ENCODE cCRE-dELS | 240,586 | 97,237 | 662 |
| myeloid DC | DC | ENCODE CTCF-only | 240,586 | 5,116 | 4 |
| myeloid DC | DC | FANTOM enhancers | 240,586 | 20,023 | 307 |
| myeloid DC | DC | Super enhancers | 240,586 | 199,154 | 1,001 |
| monocyte CD16+ | Mono | ENCODE cCRE-PLS | 279,179 | 17,832 | 752 |
| monocyte CD16+ | Mono | ENCODE cCRE-pELS | 279,179 | 28,608 | 1,146 |
| monocyte CD16+ | Mono | ENCODE cCRE-dELS | 279,179 | 94,798 | 2,289 |
| monocyte CD16+ | Mono | ENCODE CTCF-only | 279,179 | 4,828 | 35 |
| monocyte CD16+ | Mono | FANTOM enhancers | 279,179 | 18,864 | 989 |
| monocyte CD16+ | Mono | Super enhancers | 279,179 | 218,474 | 3,963 |
| monocyte CD16- | Mono | ENCODE cCRE-PLS | 348,077 | 17,952 | 652 |
| monocyte CD16- | Mono | ENCODE cCRE-pELS | 348,077 | 28,417 | 962 |
| monocyte CD16- | Mono | ENCODE cCRE-dELS | 348,077 | 97,994 | 1,520 |
| monocyte CD16- | Mono | ENCODE CTCF-only | 348,077 | 4,931 | 17 |
| monocyte CD16- | Mono | FANTOM enhancers | 348,077 | 18,360 | 681 |

| | | | | | |
|---|---|---|---|---|---|
| monocyte CD16- | Mono | Super enhancers | 348,077 | 253,729 | 2,898 |
| nonclassical Monocyte | Mono | ENCODE cCRE-PLS | 232,732 | 17,734 | 434 |
| nonclassical Monocyte | Mono | ENCODE cCRE-pELS | 232,732 | 28,311 | 658 |
| nonclassical Monocyte | Mono | ENCODE cCRE-dELS | 232,732 | 90,532 | 1,455 |
| nonclassical Monocyte | Mono | ENCODE CTCF-only | 232,732 | 4,775 | 25 |
| nonclassical Monocyte | Mono | FANTOM enhancers | 232,732 | 18,935 | 645 |
| nonclassical Monocyte | Mono | Super enhancers | 232,732 | 192,418 | 2,375 |
| intermediate Monocyte | Mono | ENCODE cCRE-PLS | 332,111 | 18,101 | 731 |
| intermediate Monocyte | Mono | ENCODE cCRE-pELS | 332,111 | 28,952 | 1,107 |
| intermediate Monocyte | Mono | ENCODE cCRE-dELS | 332,111 | 98,777 | 1,899 |
| intermediate Monocyte | Mono | ENCODE CTCF-only | 332,111 | 4,899 | 30 |
| intermediate Monocyte | Mono | FANTOM enhancers | 332,111 | 18,956 | 843 |
| intermediate Monocyte | Mono | Super enhancers | 332,111 | 245,567 | 3,487 |
| NK | NK | ENCODE cCRE-PLS | 195,677 | 17,217 | 574 |
| NK | NK | ENCODE cCRE-pELS | 195,677 | 24,726 | 793 |
| NK | NK | ENCODE cCRE-dELS | 195,677 | 67,055 | 1,363 |
| NK | NK | ENCODE CTCF-only | 195,677 | 3,895 | 23 |
| NK | NK | FANTOM enhancers | 195,677 | 13,153 | 635 |
| NK | NK | Super enhancers | 195,677 | 154,485 | 2,359 |
| PBMC | PBMC | ENCODE cCRE-PLS | 294,828 | 17,720 | 1,978 |
| PBMC | PBMC | ENCODE cCRE-pELS | 294,828 | 26,970 | 2,905 |
| PBMC | PBMC | ENCODE cCRE-dELS | 294,828 | 87,413 | 4,753 |
| PBMC | PBMC | ENCODE CTCF-only | 294,828 | 4,230 | 89 |
| PBMC | PBMC | FANTOM enhancers | 294,828 | 16,958 | 1,900 |
| PBMC | PBMC | Super enhancers | 294,828 | 222,514 | 9,028 |
| Neutrophil | Neutrophil | ENCODE cCRE-PLS | 509,449 | 16,602 | 194 |
| Neutrophil | Neutrophil | ENCODE cCRE-pELS | 509,449 | 26,123 | 299 |
| Neutrophil | Neutrophil | ENCODE cCRE-dELS | 509,449 | 104,855 | 322 |
| Neutrophil | Neutrophil | ENCODE CTCF-only | 509,449 | 4,946 | 4 |
| Neutrophil | Neutrophil | FANTOM enhancers | 509,449 | 13,909 | 138 |

| | | | | | | |
|---|---|---|---|---|---|---|
| Neutrophil | Neutrophil | Super enhancers | 509,449 | 367,128 | | 961 |

Supplementary Table 8 | Enrichment of known cCREs for cOCRs.
POP, population; OR, odds ratio; CI_lo, lower 95% confidence interval; CI_hi, upper 95% confidence interval.

| name | subgroup | cCREs | OR | CI_lo | CI_hi | P |
|---|---|---|---|---|---|---|
| Naive CD4 | CD4 | ENCODE cCRE-PLS | 4.29 | 3.77 | 4.89 | 1.40.E-81 |
| Naive CD4 | CD4 | ENCODE cCRE-pELS | 4.75 | 4.22 | 5.33 | 3.60.E-121 |
| Naive CD4 | CD4 | ENCODE cCRE-dELS | 2.32 | 2.07 | 2.59 | 2.30.E-47 |
| Naive CD4 | CD4 | ENCODE CTCF-only | 0.31 | 0.14 | 0.69 | 1.00.E+00 |
| Naive CD4 | CD4 | FANTOM enhancers | 6.16 | 5.39 | 7.03 | 6.30.E-111 |
| Naive CD4 | CD4 | Super enhancers | 9.08 | 6.8 | 12.11 | 2.00.E-102 |
| Th1 | CD4 | ENCODE cCRE-PLS | 3.65 | 3.32 | 4.01 | 7.60.E-126 |
| Th1 | CD4 | ENCODE cCRE-pELS | 3.8 | 3.49 | 4.14 | 1.10.E-171 |
| Th1 | CD4 | ENCODE cCRE-dELS | 2.55 | 2.35 | 2.76 | 2.80.E-114 |
| Th1 | CD4 | ENCODE CTCF-only | 0.32 | 0.19 | 0.54 | 1.00.E+00 |
| Th1 | CD4 | FANTOM enhancers | 5.56 | 5.07 | 6.11 | 6.10.E-211 |
| Th1 | CD4 | Super enhancers | 5.66 | 4.72 | 6.8 | 3.30.E-131 |
| Th2 | CD4 | ENCODE cCRE-PLS | 4.17 | 3.78 | 4.59 | 1.10.E-140 |
| Th2 | CD4 | ENCODE cCRE-pELS | 4.39 | 4.03 | 4.8 | 1.40.E-195 |

| | | | | | | |
|---|---|---|---|---|---|---|
| Th2 | CD4 | ENCODE cCRE-dELS | 2.6 | 2.39 | 2.82 | 7.60.E-111 |
| Th2 | CD4 | ENCODE CTCF-only | 0.35 | 0.21 | 0.6 | 1.00.E+00 |
| Th2 | CD4 | FANTOM enhancers | 6.04 | 5.48 | 6.66 | 1.20.E-205 |
| Th2 | CD4 | Super enhancers | 6.95 | 5.72 | 8.43 | 6.40.E-155 |
| Th17 | CD4 | ENCODE cCRE-PLS | 3.43 | 3.18 | 3.71 | 3.20.E-170 |
| Th17 | CD4 | ENCODE cCRE-pELS | 3.59 | 3.35 | 3.85 | 9.80.E-238 |
| Th17 | CD4 | ENCODE cCRE-dELS | 2.67 | 2.5 | 2.84 | 2.20.E-196 |
| Th17 | CD4 | ENCODE CTCF-only | 0.56 | 0.41 | 0.76 | 1.00.E+00 |
| Th17 | CD4 | FANTOM enhancers | 5.01 | 4.64 | 5.4 | 2.90.E-278 |
| Th17 | CD4 | Super enhancers | 5.17 | 4.51 | 5.92 | 1.10.E-201 |
| Tfh | CD4 | ENCODE cCRE-PLS | 4.28 | 3.81 | 4.8 | 5.50.E-103 |
| Tfh | CD4 | ENCODE cCRE-pELS | 4.55 | 4.09 | 5.05 | 5.30.E-144 |
| Tfh | CD4 | ENCODE cCRE-dELS | 2.54 | 2.3 | 2.81 | 8.80.E-74 |
| Tfh | CD4 | ENCODE CTCF-only | 0.29 | 0.14 | 0.57 | 1.00.E+00 |
| Tfh | CD4 | FANTOM enhancers | 6.49 | 5.78 | 7.28 | 3.00.E-159 |
| Tfh | CD4 | Super enhancers | 8 | 6.24 | 10.26 | 1.90.E-116 |
| FraI | CD4 | ENCODE cCRE-PLS | 3.96 | 3.49 | 4.51 | 5.50.E-76 |

| | | | | | | |
|---|---|---|---|---|---|---|
| FraI | CD4 | ENCODE cCRE-pELS | 4.32 | 3.85 | 4.85 | 8.00.E-111 |
| FraI | CD4 | ENCODE cCRE-dELS | 2.34 | 2.09 | 2.61 | 2.20.E-50 |
| FraI | CD4 | ENCODE CTCF-only | 0.33 | 0.16 | 0.7 | 1.00.E+00 |
| FraI | CD4 | FANTOM enhancers | 6.15 | 5.41 | 7 | 1.10.E-120 |
| FraI | CD4 | Super enhancers | 8.9 | 6.67 | 11.87 | 1.60.E-100 |
| FraII (aTreg) | CD4 | ENCODE cCRE-PLS | 3.87 | 3.44 | 4.36 | 8.10.E-85 |
| FraII (aTreg) | CD4 | ENCODE cCRE-pELS | 4.21 | 3.78 | 4.68 | 3.70.E-124 |
| FraII (aTreg) | CD4 | ENCODE cCRE-dELS | 2.41 | 2.18 | 2.67 | 2.40.E-63 |
| FraII (aTreg) | CD4 | ENCODE CTCF-only | 0.27 | 0.13 | 0.57 | 1.00.E+00 |
| FraII (aTreg) | CD4 | FANTOM enhancers | 5.91 | 5.25 | 6.65 | 8.30.E-136 |
| FraII (aTreg) | CD4 | Super enhancers | 7.73 | 5.98 | 9.99 | 3.50.E-105 |
| FraIII | CD4 | ENCODE cCRE-PLS | 3.04 | 2.8 | 3.3 | 1.30.E-127 |
| FraIII | CD4 | ENCODE cCRE-pELS | 3.11 | 2.89 | 3.35 | 2.20.E-170 |
| FraIII | CD4 | ENCODE cCRE-dELS | 2.56 | 2.39 | 2.74 | 1.50.E-160 |
| FraIII | CD4 | ENCODE CTCF-only | 0.4 | 0.28 | 0.59 | 1.00.E+00 |
| FraIII | CD4 | FANTOM enhancers | 4.88 | 4.51 | 5.29 | 9.80.E-254 |
| FraIII | CD4 | Super enhancers | 5.07 | 4.36 | 5.89 | 7.60.E-160 |
| LAG3 Treg | CD4 | ENCODE cCRE-PLS | 2.89 | 2.68 | 3.12 | 9.90.E-137 |

| | | | | | | |
|---|---|---|---|---|---|---|
| LAG3 Treg | CD4 | ENCODE cCRE-pELS | 3.01 | 2.81 | 3.22 | 9.10.E-191 |
| LAG3 Treg | CD4 | ENCODE cCRE-dELS | 2.53 | 2.38 | 2.69 | 2.60.E-187 |
| LAG3 Treg | CD4 | ENCODE CTCF-only | 0.46 | 0.33 | 0.63 | 1.00.E+00 |
| LAG3 Treg | CD4 | FANTOM enhancers | 4.33 | 4.02 | 4.67 | 7.90.E-248 |
| LAG3 Treg | CD4 | Super enhancers | 4.29 | 3.76 | 4.89 | 2.60.E-161 |
| Memory CD4 | CD4 | ENCODE cCRE-PLS | 4.14 | 3.77 | 4.55 | 9.70.E-152 |
| Memory CD4 | CD4 | ENCODE cCRE-pELS | 4.46 | 4.1 | 4.84 | 2.60.E-218 |
| Memory CD4 | CD4 | ENCODE cCRE-dELS | 2.54 | 2.35 | 2.75 | 5.50.E-115 |
| Memory CD4 | CD4 | ENCODE CTCF-only | 0.4 | 0.25 | 0.65 | 1.00.E+00 |
| Memory CD4 | CD4 | FANTOM enhancers | 5.82 | 5.3 | 6.39 | 4.80.E-216 |
| Memory CD4 | CD4 | Super enhancers | 6.59 | 5.49 | 7.91 | 5.20.E-163 |
| Thx | CD4 | ENCODE cCRE-PLS | 3.25 | 2.99 | 3.53 | 5.10.E-141 |
| Thx | CD4 | ENCODE cCRE-pELS | 3.34 | 3.11 | 3.6 | 7.30.E-191 |
| Thx | CD4 | ENCODE cCRE-dELS | 2.54 | 2.37 | 2.72 | 1.60.E-158 |
| Thx | CD4 | ENCODE CTCF-only | 0.44 | 0.3 | 0.63 | 1.00.E+00 |
| Thx | CD4 | FANTOM enhancers | 4.91 | 4.53 | 5.32 | 3.10.E-249 |

| | | | | | | |
|---|---|---|---|---|---|---|
| Thx | CD4 | Super enhancers | 4.95 | 4.27 | 5.75 | 1.10.E-158 |
| Naive CD8 | CD8 | ENCODE cCRE-PLS | 3.53 | 2.61 | 4.79 | 1.30.E-12 |
| Naive CD8 | CD8 | ENCODE cCRE-pELS | 3.46 | 2.65 | 4.52 | 6.20.E-16 |
| Naive CD8 | CD8 | ENCODE cCRE-dELS | 1.3 | 1.01 | 1.66 | 2.50.E-02 |
| Naive CD8 | CD8 | ENCODE CTCF-only | 0.12 | 0.01 | 1.88 | 1.00.E+00 |
| Naive CD8 | CD8 | FANTOM enhancers | 4.51 | 3.28 | 6.19 | 4.80.E-15 |
| Naive CD8 | CD8 | Super enhancers | 19.3 | 8.6 | 43.31 | 2.10.E-34 |
| CD8+effector | CD8 | ENCODE cCRE-PLS | 4.52 | 3.83 | 5.33 | 1.90.E-54 |
| CD8+effector | CD8 | ENCODE cCRE-pELS | 4.68 | 4.04 | 5.43 | 4.10.E-74 |
| CD8+effector | CD8 | ENCODE cCRE-dELS | 2.26 | 1.97 | 2.59 | 1.40.E-29 |
| CD8+effector | CD8 | ENCODE CTCF-only | 0.16 | 0.04 | 0.63 | 1.00.E+00 |
| CD8+effector | CD8 | FANTOM enhancers | 7.23 | 6.18 | 8.47 | 3.00.E-93 |
| CD8+effector | CD8 | Super enhancers | 10.89 | 7.37 | 16.1 | 9.00.E-72 |
| CD8+central memory | CD8 | ENCODE cCRE-PLS | 4.24 | 3.7 | 4.87 | 9.60.E-72 |
| CD8+central memory | CD8 | ENCODE cCRE-pELS | 4.68 | 4.14 | 5.29 | 3.80.E-107 |
| CD8+central memory | CD8 | ENCODE cCRE-dELS | 2.5 | 2.22 | 2.8 | 9.00.E-52 |
| CD8+central memory | CD8 | ENCODE CTCF-only | 0.11 | 0.03 | 0.45 | 1.00.E+00 |
| CD8+central memory | CD8 | FANTOM enhancers | 6.45 | 5.63 | 7.39 | 6.10.E-112 |
| CD8+central memory | CD8 | Super enhancers | 10.15 | 7.37 | 13.97 | 2.30.E-97 |
| CD8+effector memory | CD8 | ENCODE cCRE-PLS | 4.76 | 4.01 | 5.64 | 1.40.E-53 |
| CD8+effector memory | CD8 | ENCODE cCRE-pELS | 4.85 | 4.15 | 5.66 | 1.60.E-70 |
| CD8+effector memory | CD8 | ENCODE cCRE-dELS | 2.12 | 1.83 | 2.46 | 9.70.E-23 |
| CD8+effector memory | CD8 | ENCODE CTCF-only | 0.09 | 0.01 | 0.65 | 1.00.E+00 |
| CD8+effector memory | CD8 | FANTOM enhancers | 6.7 | 5.65 | 7.94 | 2.60.E-74 |
| CD8+effector memory | CD8 | Super enhancers | 11.31 | 7.4 | 17.28 | 3.30.E-64 |
| Naive B | B | ENCODE cCRE-PLS | 3.3 | 2.37 | 4.61 | 6.90.E-10 |
| Naive B | B | ENCODE cCRE-pELS | 3.21 | 2.4 | 4.29 | 2.10.E-12 |

| | | | | | | |
|---|---|---|---|---|---|---|
| Naive B | B | ENCODE cCRE-dELS | 0.94 | 0.71 | 1.25 | 6.80.E-01 |
| Naive B | B | ENCODE CTCF-only | 0.27 | 0.04 | 1.96 | 9.70.E-01 |
| Naive B | B | FANTOM enhancers | 4.49 | 3.22 | 6.26 | 8.70.E-14 |
| Naive B | B | Super enhancers | 35.3 | 11.31 | 110.14 | 1.50.E-34 |
| unsw memory B | B | ENCODE cCRE-PLS | 4.41 | 3.71 | 5.24 | 5.20.E-47 |
| unsw memory B | B | ENCODE cCRE-pELS | 4.72 | 4.06 | 5.5 | 9.30.E-70 |
| unsw memory B | B | ENCODE cCRE-dELS | 2.02 | 1.75 | 2.33 | 2.10.E-21 |
| unsw memory B | B | ENCODE CTCF-only | 0.18 | 0.04 | 0.7 | 1.00.E+00 |
| unsw memory B | B | FANTOM enhancers | 5.54 | 4.65 | 6.6 | 7.30.E-58 |
| unsw memory B | B | Super enhancers | 9.91 | 6.88 | 14.27 | 9.80.E-72 |
| sw memory B | B | ENCODE cCRE-PLS | 5.27 | 4.57 | 6.08 | 1.50.E-84 |
| sw memory B | B | ENCODE cCRE-pELS | 5.15 | 4.53 | 5.86 | 1.50.E-107 |
| sw memory B | B | ENCODE cCRE-dELS | 2.18 | 1.93 | 2.46 | 8.30.E-35 |
| sw memory B | B | ENCODE CTCF-only | 0.13 | 0.03 | 0.52 | 1.00.E+00 |
| sw memory B | B | FANTOM enhancers | 5.92 | 5.11 | 6.86 | 2.10.E-87 |
| sw memory B | B | Super enhancers | 9.88 | 7.15 | 13.66 | 3.70.E-91 |
| DN B | B | ENCODE cCRE-PLS | 5.11 | 4.52 | 5.78 | 1.30.E-108 |
| DN B | B | ENCODE cCRE-pELS | 5.32 | 4.76 | 5.94 | 7.40.E-151 |
| DN B | B | ENCODE cCRE-dELS | 2.37 | 2.14 | 2.64 | 5.80.E-57 |
| DN B | B | ENCODE CTCF-only | 0.24 | 0.1 | 0.59 | 1.00.E+00 |
| DN B | B | FANTOM enhancers | 5.7 | 5.02 | 6.49 | 1.40.E-110 |
| DN B | B | Super enhancers | 10.69 | 8.04 | 14.22 | 3.10.E-129 |
| plasmablast | B | ENCODE cCRE-PLS | 3.71 | 3.41 | 4.04 | 1.00.E-163 |

| | | | | | | |
|---|---|---|---|---|---|---|
| plasmablast | B | ENCODE cCRE-pELS | 3.92 | 3.64 | 4.23 | 2.40.E-231 |
| plasmablast | B | ENCODE cCRE-dELS | 2.11 | 1.97 | 2.26 | 6.70.E-95 |
| plasmablast | B | ENCODE CTCF-only | 0.57 | 0.41 | 0.8 | 1.00.E+00 |
| plasmablast | B | FANTOM enhancers | 4.16 | 3.8 | 4.54 | 1.30.E-162 |
| plasmablast | B | Super enhancers | 4.91 | 4.2 | 5.74 | 3.80.E-143 |
| plasmacytoid DC | DC | ENCODE cCRE-PLS | 2.35 | 2 | 2.76 | 1.80.E-21 |
| plasmacytoid DC | DC | ENCODE cCRE-pELS | 2.08 | 1.8 | 2.4 | 4.30.E-21 |
| plasmacytoid DC | DC | ENCODE cCRE-dELS | 1.79 | 1.58 | 2.03 | 2.60.E-20 |
| plasmacytoid DC | DC | ENCODE CTCF-only | 0.35 | 0.18 | 0.68 | 1.00.E+00 |
| plasmacytoid DC | DC | FANTOM enhancers | 3.95 | 3.43 | 4.56 | 1.20.E-61 |
| plasmacytoid DC | DC | Super enhancers | 1.93 | 1.56 | 2.38 | 2.40.E-11 |
| myeloid DC | DC | ENCODE cCRE-PLS | 2.41 | 2.05 | 2.83 | 1.80.E-22 |
| myeloid DC | DC | ENCODE cCRE-pELS | 2.27 | 1.98 | 2.62 | 6.50.E-27 |
| myeloid DC | DC | ENCODE cCRE-dELS | 2.36 | 2.08 | 2.67 | 4.60.E-44 |
| myeloid DC | DC | ENCODE CTCF-only | 0.17 | 0.06 | 0.46 | 1.00.E+00 |
| myeloid DC | DC | FANTOM enhancers | 4.4 | 3.85 | 5.02 | 1.00.E-82 |
| myeloid DC | DC | Super enhancers | 2.78 | 2.2 | 3.51 | 4.50.E-23 |
| monocyte CD16+ | Mono | ENCODE cCRE-PLS | 3.16 | 2.92 | 3.42 | 4.00.E-137 |
| monocyte CD16+ | Mono | ENCODE cCRE-pELS | 3.24 | 3.03 | 3.47 | 1.00.E-202 |
| monocyte CD16+ | Mono | ENCODE cCRE-dELS | 2.28 | 2.15 | 2.42 | 1.70.E-154 |
| monocyte CD16+ | Mono | ENCODE CTCF-only | 0.47 | 0.34 | 0.66 | 1.00.E+00 |
| monocyte CD16+ | Mono | FANTOM enhancers | 4.2 | 3.9 | 4.51 | 2.30.E-252 |

| | | | | | | |
|---|---|---|---|---|---|---|
| monocyte CD16+ | Mono | Super enhancers | 3.96 | 3.51 | 4.47 | 7.30.E-161 |
| monocyte CD16- | Mono | ENCODE cCRE-PLS | 5.03 | 4.61 | 5.49 | 8.10.E-208 |
| monocyte CD16- | Mono | ENCODE cCRE-pELS | 5.22 | 4.83 | 5.64 | 1.90.E-298 |
| monocyte CD16- | Mono | ENCODE cCRE-dELS | 2.56 | 2.38 | 2.75 | 1.20.E-141 |
| monocyte CD16- | Mono | ENCODE CTCF-only | 0.39 | 0.24 | 0.63 | 1.00.E+00 |
| monocyte CD16- | Mono | FANTOM enhancers | 5.2 | 4.76 | 5.67 | 6.80.E-223 |
| monocyte CD16- | Mono | Super enhancers | 7.87 | 6.63 | 9.34 | 9.00.E-234 |
| nonclassical Monocyte | Mono | ENCODE cCRE-PLS | 2.48 | 2.24 | 2.76 | 3.80.E-54 |
| nonclassical Monocyte | Mono | ENCODE cCRE-pELS | 2.51 | 2.29 | 2.74 | 3.20.E-77 |
| nonclassical Monocyte | Mono | ENCODE cCRE-dELS | 2.08 | 1.92 | 2.25 | 2.60.E-75 |
| nonclassical Monocyte | Mono | ENCODE CTCF-only | 0.47 | 0.32 | 0.7 | 1.00.E+00 |
| nonclassical Monocyte | Mono | FANTOM enhancers | 3.82 | 3.49 | 4.18 | 2.10.E-146 |
| nonclassical Monocyte | Mono | Super enhancers | 2.8 | 2.4 | 3.26 | 6.80.E-53 |
| intermediate Monocyte | Mono | ENCODE cCRE-PLS | 4.3 | 3.96 | 4.67 | 1.10.E-197 |
| intermediate Monocyte | Mono | ENCODE cCRE-pELS | 4.51 | 4.2 | 4.84 | 4.20.E-295 |
| intermediate Monocyte | Mono | ENCODE cCRE-dELS | 2.52 | 2.36 | 2.69 | 1.10.E-166 |
| intermediate Monocyte | Mono | ENCODE CTCF-only | 0.55 | 0.38 | 0.79 | 1.00.E+00 |
| intermediate Monocyte | Mono | FANTOM enhancers | 4.91 | 4.54 | 5.31 | 2.30.E-258 |

| | | | | | | |
|---|---|---|---|---|---|---|
| intermediate Monocyte | Mono | Super enhancers | 6.4 | 5.53 | 7.4 | 3.90.E-242 |
| NK | NK | ENCODE cCRE-PLS | 3.12 | 2.83 | 3.42 | 1.10.E-99 |
| NK | NK | ENCODE cCRE-pELS | 3.24 | 2.98 | 3.53 | 4.80.E-136 |
| NK | NK | ENCODE cCRE-dELS | 2.33 | 2.15 | 2.53 | 1.40.E-96 |
| NK | NK | ENCODE CTCF-only | 0.46 | 0.3 | 0.69 | 1.00.E+00 |
| NK | NK | FANTOM enhancers | 4.77 | 4.35 | 5.22 | 9.30.E-186 |
| NK | NK | Super enhancers | 5.03 | 4.2 | 6.03 | 1.40.E-112 |
| PBMC | PBMC | ENCODE cCRE-PLS | 4.09 | 3.89 | 4.31 | 0.00.E+00 |
| PBMC | PBMC | ENCODE cCRE-pELS | 4.35 | 4.16 | 4.55 | 0.00.E+00 |
| PBMC | PBMC | ENCODE cCRE-dELS | 2.36 | 2.26 | 2.46 | 0.00.E+00 |
| PBMC | PBMC | ENCODE CTCF-only | 0.65 | 0.52 | 0.8 | 1.00.E+00 |
| PBMC | PBMC | FANTOM enhancers | 4.08 | 3.87 | 4.3 | 0.00.E+00 |
| PBMC | PBMC | Super enhancers | 5.8 | 5.3 | 6.34 | 0.00.E+00 |
| Neutrophil | Neutrophil | ENCODE cCRE-PLS | 7.25 | 6.19 | 8.49 | 3.60.E-89 |
| Neutrophil | Neutrophil | ENCODE cCRE-pELS | 8.03 | 7.01 | 9.2 | 4.10.E-140 |
| Neutrophil | Neutrophil | ENCODE cCRE-dELS | 1.87 | 1.63 | 2.13 | 8.80.E-19 |
| Neutrophil | Neutrophil | ENCODE CTCF-only | 0.41 | 0.16 | 1.11 | 9.90.E-01 |
| Neutrophil | Neutrophil | FANTOM enhancers | 5.78 | 4.83 | 6.93 | 2.10.E-54 |
| Neutrophil | Neutrophil | Super enhancers | 13.8 | 9.41 | 20.23 | 1.40.E-99 |

235

Supplementary Table 9 | Enrichment of cOCRs in the known cCREs adjusted for seven established constraint metrics and sequence features.
OR, odds ratio; CI_lo, lower 95% confidence interval;
CI_hi, upper 95% confidence interval.

| name | subgroup | type | cCREs | OR | CI_lo | CI_hi | P | OR (non_coding) | CI_lo (non_coding) | CI_hi (non_coding) | P (non_coding) |
|---|---|---|---|---|---|---|---|---|---|---|---|
| Naive CD4 | CD4 | cOCRs | ENCODE cCRE-PLS | 4.52 | 3.66 | 5.58 | 2.87.E-44 | 4.74 | 3.49 | 6.46 | 4.05.E-23 |
| Naive CD4 | CD4 | CAMBUS_control | ENCODE cCRE-PLS | 0.44 | 0.37 | 0.53 | 2.63.E-18 | 0.49 | 0.37 | 0.64 | 1.49.E-07 |
| Naive CD4 | CD4 | cOCRs | ENCODE cCRE-pELS | 4.40 | 3.64 | 5.32 | 2.80.E-53 | 3.94 | 3.06 | 5.08 | 2.23.E-26 |
| Naive CD4 | CD4 | CAMBUS_control | ENCODE cCRE-pELS | 0.42 | 0.36 | 0.49 | 2.51.E-28 | 0.46 | 0.37 | 0.56 | 1.87.E-13 |
| Naive CD4 | CD4 | cOCRs | ENCODE cCRE-dELS | 2.72 | 2.28 | 3.23 | 1.22.E-29 | 3.50 | 2.78 | 4.40 | 8.03.E-27 |
| Naive CD4 | CD4 | CAMBUS_control | ENCODE cCRE-dELS | 0.65 | 0.57 | 0.74 | 2.06.E-11 | 0.65 | 0.56 | 0.75 | 5.22.E-09 |
| Naive CD4 | CD4 | cOCRs | ENCODE CTCF-only | 0.30 | 0.08 | 1.20 | 8.98.E-02 | 0.42 | 0.10 | 1.67 | 2.15.E-01 |

| | | | | | | | | | | | |
|---|---|---|---|---|---|---|---|---|---|---|---|
| Naive CD4 | CD4 | CAMBUS_control | ENCODE CTCF-only | 1.47 | 0.81 | 2.66 | 2.08.E-01 | 1.24 | 0.68 | 2.26 | 4.75.E-01 |
| Naive CD4 | CD4 | cOCRs | FANTOM enhancers | 6.42 | 5.30 | 7.77 | 4.37.E-81 | 9.02 | 7.18 | 11.33 | 1.13.E-79 |
| Naive CD4 | CD4 | CAMBUS_control | FANTOM enhancers | 0.27 | 0.23 | 0.32 | 2.04.E-52 | 0.24 | 0.20 | 0.29 | 2.10.E-50 |
| Naive CD4 | CD4 | cOCRs | Super enhancers | 6.18 | 3.80 | 10.05 | 2.12.E-13 | 10.67 | 5.02 | 22.66 | 7.34.E-10 |
| Naive CD4 | CD4 | CAMBUS_control | Super enhancers | 0.44 | 0.37 | 0.52 | 2.87.E-22 | 0.43 | 0.36 | 0.52 | 1.47.E-20 |
| Th1 | CD4 | cOCRs | ENCODE cCRE-PLS | 4.55 | 3.94 | 5.24 | 1.25.E-96 | 4.96 | 4.06 | 6.07 | 1.20.E-54 |
| Th1 | CD4 | CAMBUS_control | ENCODE cCRE-PLS | 0.33 | 0.29 | 0.37 | 8.56.E-62 | 0.32 | 0.27 | 0.39 | 5.34.E-32 |
| Th1 | CD4 | cOCRs | ENCODE cCRE-pELS | 4.09 | 3.61 | 4.63 | 2.22.E-110 | 3.62 | 3.08 | 4.25 | 1.71.E-55 |
| Th1 | CD4 | CAMBUS_control | ENCODE cCRE-pELS | 0.34 | 0.30 | 0.38 | 6.44.E-79 | 0.37 | 0.32 | 0.43 | 1.87.E-39 |
| Th1 | CD4 | cOCRs | ENCOD | 2.71 | 2.42 | 3.04 | 4.15.E- | 3.64 | 3.15 | 4.21 | 6.61.E- |

| | | | | | | | | | | | |
|---|---|---|---|---|---|---|---|---|---|---|---|
| | | | E cCRE-dELS | | | | 66 | | | | 68 |
| Th1 | CD4 | CAMBUS_control | ENCODE cCRE-dELS | 0.48 | 0.44 | 0.53 | 1.23.E-49 | 0.42 | 0.38 | 0.47 | 4.13.E-50 |
| Th1 | CD4 | cOCRs | ENCODE CTCF-only | 0.30 | 0.14 | 0.63 | 1.51.E-03 | 0.32 | 0.15 | 0.73 | 6.13.E-03 |
| Th1 | CD4 | CAMBUS_control | ENCODE CTCF-only | 1.88 | 1.16 | 3.04 | 9.93.E-03 | 1.66 | 1.01 | 2.73 | 4.43.E-02 |
| Th1 | CD4 | cOCRs | FANTOM enhancers | 5.51 | 4.88 | 6.23 | 2.18.E-163 | 7.02 | 6.09 | 8.08 | 1.12.E-160 |
| Th1 | CD4 | CAMBUS_control | FANTOM enhancers | 0.23 | 0.20 | 0.26 | 3.49.E-139 | 0.20 | 0.17 | 0.22 | 5.68.E-136 |
| Th1 | CD4 | cOCRs | Super enhancers | 4.02 | 3.12 | 5.18 | 4.59.E-27 | 5.96 | 4.21 | 8.43 | 6.06.E-24 |
| Th1 | CD4 | CAMBUS_control | Super enhancers | 0.42 | 0.37 | 0.49 | 4.52.E-29 | 0.39 | 0.33 | 0.46 | 3.93.E-28 |
| Th2 | CD4 | cOCRs | ENCODE cCRE-PLS | 5.09 | 4.39 | 5.91 | 1.96.E-102 | 5.46 | 4.42 | 6.76 | 4.43.E-55 |
| Th2 | CD4 | CAMBUS_contro | ENCODE cCRE- | 0.35 | 0.31 | 0.40 | 5.11.E-55 | 0.36 | 0.30 | 0.44 | 3.05.E-25 |

| | | | | | | | | | | | |
|---|---|---|---|---|---|---|---|---|---|---|---|
| | | l | PLS | | | | | | | | |
| Th2 | CD4 | cOCRs | ENCODE cCRE-pELS | 4.58 | 4.02 | 5.22 | 2.00.E-116 | 4.07 | 3.44 | 4.83 | 9.31.E-59 |
| Th2 | CD4 | CAMBUS_control | ENCODE cCRE-pELS | 0.37 | 0.33 | 0.41 | 6.89.E-68 | 0.42 | 0.36 | 0.49 | 1.32.E-30 |
| Th2 | CD4 | cOCRs | ENCODE cCRE-dELS | 2.71 | 2.40 | 3.05 | 7.43.E-61 | 3.80 | 3.26 | 4.43 | 6.51.E-65 |
| Th2 | CD4 | CAMBUS_control | ENCODE cCRE-dELS | 0.61 | 0.56 | 0.67 | 3.98.E-26 | 0.57 | 0.52 | 0.63 | 2.25.E-26 |
| Th2 | CD4 | cOCRs | ENCODE CTCF-only | 0.26 | 0.11 | 0.63 | 2.82.E-03 | 0.27 | 0.10 | 0.71 | 8.37.E-03 |
| Th2 | CD4 | CAMBUS_control | ENCODE CTCF-only | 1.93 | 1.23 | 3.04 | 4.42.E-03 | 1.76 | 1.10 | 2.80 | 1.79.E-02 |
| Th2 | CD4 | cOCRs | FANTOM enhancers | 6.07 | 5.34 | 6.91 | 9.88.E-164 | 8.13 | 7.00 | 9.45 | 4.50.E-165 |
| Th2 | CD4 | CAMBUS_control | FANTOM enhancers | 0.27 | 0.24 | 0.30 | 8.61.E-105 | 0.24 | 0.21 | 0.27 | 3.35.E-103 |
| Th2 | CD4 | cOCRs | Super enhancers | 4.93 | 3.71 | 6.55 | 3.68.E-28 | 7.98 | 5.30 | 12.02 | 2.81.E-23 |

| | | | | | | | | | | | |
|---|---|---|---|---|---|---|---|---|---|---|---|
| Th2 | CD4 | CAMBUS_control | Super enhancers | 0.47 | 0.41 | 0.53 | 1.66.E-35 | 0.45 | 0.40 | 0.51 | 2.89.E-33 |
| Th17 | CD4 | cOCRs | ENCODE cCRE-PLS | 4.41 | 3.94 | 4.95 | 3.29.E-143 | 4.69 | 3.97 | 5.53 | 4.88.E-75 |
| Th17 | CD4 | CAMBUS_control | ENCODE cCRE-PLS | 0.29 | 0.26 | 0.32 | 2.80.E-111 | 0.28 | 0.24 | 0.33 | 1.29.E-55 |
| Th17 | CD4 | cOCRs | ENCODE cCRE-pELS | 4.05 | 3.67 | 4.47 | 2.74.E-171 | 3.63 | 3.19 | 4.13 | 6.05.E-86 |
| Th17 | CD4 | CAMBUS_control | ENCODE cCRE-pELS | 0.31 | 0.28 | 0.34 | 8.63.E-137 | 0.34 | 0.30 | 0.39 | 1.55.E-66 |
| Th17 | CD4 | cOCRs | ENCODE cCRE-dELS | 2.77 | 2.54 | 3.03 | 4.21.E-111 | 3.72 | 3.32 | 4.17 | 4.18.E-114 |
| Th17 | CD4 | CAMBUS_control | ENCODE cCRE-dELS | 0.44 | 0.40 | 0.47 | 2.83.E-91 | 0.37 | 0.34 | 0.41 | 7.15.E-93 |
| Th17 | CD4 | cOCRs | ENCODE CTCF-only | 0.52 | 0.34 | 0.79 | 2.35.E-03 | 0.60 | 0.38 | 0.93 | 2.22.E-02 |
| Th17 | CD4 | CAMBUS_control | ENCODE CTCF-only | 1.80 | 1.25 | 2.61 | 1.69.E-03 | 1.59 | 1.09 | 2.33 | 1.74.E-02 |
| Th17 | CD4 | cOCRs | FANTOM enhance | 5.31 | 4.81 | 5.86 | 1.02.E-242 | 6.73 | 6.01 | 7.53 | 5.58.E-243 |

| | | | | | | | | | | | |
|---|---|---|---|---|---|---|---|---|---|---|---|
| | | | rs | | | | | | | | |
| Th17 | CD4 | CAMBUS_control | FANTOM enhancers | 0.22 | 0.20 | 0.24 | 1.24.E-212 | 0.19 | 0.17 | 0.21 | 3.88.E-212 |
| Th17 | CD4 | cOCRs | Super enhancers | 4.26 | 3.51 | 5.17 | 6.50.E-49 | 5.03 | 3.96 | 6.38 | 4.50.E-40 |
| Th17 | CD4 | CAMBUS_control | Super enhancers | 0.36 | 0.32 | 0.41 | 6.41.E-51 | 0.35 | 0.30 | 0.40 | 2.42.E-44 |
| Tfh | CD4 | cOCRs | ENCODE cCRE-PLS | 4.89 | 4.09 | 5.85 | 2.08.E-68 | 6.04 | 4.71 | 7.74 | 1.15.E-45 |
| Tfh | CD4 | CAMBUS_control | ENCODE cCRE-PLS | 0.34 | 0.29 | 0.40 | 2.55.E-40 | 0.31 | 0.25 | 0.39 | 1.43.E-24 |
| Tfh | CD4 | cOCRs | ENCODE cCRE-pELS | 4.80 | 4.11 | 5.62 | 5.65.E-86 | 4.79 | 3.92 | 5.86 | 1.47.E-52 |
| Tfh | CD4 | CAMBUS_control | ENCODE cCRE-pELS | 0.33 | 0.29 | 0.38 | 4.08.E-55 | 0.34 | 0.28 | 0.40 | 4.18.E-33 |
| Tfh | CD4 | cOCRs | ENCODE cCRE-dELS | 2.73 | 2.36 | 3.16 | 8.60.E-42 | 3.52 | 2.92 | 4.25 | 1.61.E-39 |
| Tfh | CD4 | CAMBUS_control | ENCODE cCRE-dELS | 0.55 | 0.49 | 0.62 | 3.24.E-23 | 0.54 | 0.47 | 0.61 | 2.28.E-19 |
| Tfh | CD4 | cOCRs | ENCOD | 0.24 | 0.08 | 0.74 | 1.30.E- | 0.21 | 0.05 | 0.83 | 2.62.E- |

| | | | | | | | | | | | |
|---|---|---|---|---|---|---|---|---|---|---|---|
| | | | E CTCF-only | | | | 02 | | | | 02 |
| Tfh | CD4 | CAMBUS_control | ENCODE CTCF-only | 2.64 | 1.31 | 5.29 | 6.37.E-03 | 2.93 | 1.31 | 6.55 | 8.73.E-03 |
| Tfh | CD4 | cOCRs | FANTOM enhancers | 6.71 | 5.74 | 7.84 | 4.02.E-127 | 8.75 | 7.28 | 10.53 | 6.02.E-118 |
| Tfh | CD4 | CAMBUS_control | FANTOM enhancers | 0.21 | 0.19 | 0.25 | 2.00.E-98 | 0.19 | 0.16 | 0.22 | 8.58.E-91 |
| Tfh | CD4 | cOCRs | Super enhancers | 4.83 | 3.40 | 6.85 | 1.06.E-18 | 8.20 | 4.88 | 13.78 | 1.84.E-15 |
| Tfh | CD4 | CAMBUS_control | Super enhancers | 0.50 | 0.42 | 0.58 | 3.22.E-18 | 0.48 | 0.40 | 0.57 | 4.08.E-17 |
| FraI | CD4 | cOCRs | ENCODE cCRE-PLS | 3.91 | 3.18 | 4.80 | 1.33.E-38 | 4.72 | 3.51 | 6.35 | 1.06.E-24 |
| FraI | CD4 | CAMBUS_control | ENCODE cCRE-PLS | 0.42 | 0.35 | 0.51 | 2.73.E-19 | 0.43 | 0.32 | 0.56 | 1.57.E-09 |
| FraI | CD4 | cOCRs | ENCODE cCRE-pELS | 3.99 | 3.33 | 4.79 | 9.08.E-50 | 3.82 | 2.99 | 4.88 | 5.54.E-27 |
| FraI | CD4 | CAMBUS_contro | ENCODE cCRE- | 0.39 | 0.33 | 0.46 | 1.18.E-30 | 0.41 | 0.33 | 0.51 | 2.93.E-16 |

| | | | | | | | | | | | |
|---|---|---|---|---|---|---|---|---|---|---|---|
| | | l | pELS | | | | | | | | |
| FraI | CD4 | cOCRs | ENCODE cCRE-dELS | 2.45 | 2.07 | 2.90 | 1.65.E-25 | 3.17 | 2.54 | 3.95 | 2.33.E-24 |
| FraI | CD4 | CAMBUS_control | ENCODE cCRE-dELS | 0.60 | 0.52 | 0.69 | 3.06.E-13 | 0.59 | 0.50 | 0.69 | 8.22.E-11 |
| FraI | CD4 | cOCRs | ENCODE CTCF-only | 0.38 | 0.12 | 1.20 | 9.93.E-02 | 0.52 | 0.17 | 1.64 | 2.66.E-01 |
| FraI | CD4 | CAMBUS_control | ENCODE CTCF-only | 1.49 | 0.77 | 2.87 | 2.38.E-01 | 1.23 | 0.64 | 2.39 | 5.32.E-01 |
| FraI | CD4 | cOCRs | FANTOM enhancers | 5.78 | 4.80 | 6.96 | 1.71.E-76 | 8.14 | 6.52 | 10.17 | 1.55.E-76 |
| FraI | CD4 | CAMBUS_control | FANTOM enhancers | 0.25 | 0.21 | 0.30 | 1.81.E-55 | 0.21 | 0.18 | 0.26 | 3.11.E-54 |
| FraI | CD4 | cOCRs | Super enhancers | 5.36 | 3.42 | 8.40 | 2.65.E-13 | 10.94 | 5.15 | 23.23 | 4.67.E-10 |
| FraI | CD4 | CAMBUS_control | Super enhancers | 0.48 | 0.39 | 0.58 | 1.28.E-14 | 0.46 | 0.38 | 0.57 | 1.49.E-13 |
| FraII (aTreg) | CD4 | cOCRs | ENCODE cCRE-PLS | 4.73 | 3.93 | 5.69 | 1.32.E-60 | 6.01 | 4.65 | 7.76 | 8.95.E-43 |

| | | | | | | | | | | | |
|---|---|---|---|---|---|---|---|---|---|---|---|
| FraII (aTreg) | CD4 | CAMBUS_control | ENCODE cCRE-PLS | 0.52 | 0.45 | 0.61 | 7.09.E-17 | 0.49 | 0.40 | 0.61 | 5.91.E-11 |
| FraII (aTreg) | CD4 | cOCRs | ENCODE cCRE-pELS | 4.71 | 4.01 | 5.53 | 5.81.E-79 | 4.57 | 3.72 | 5.62 | 6.91.E-47 |
| FraII (aTreg) | CD4 | CAMBUS_control | ENCODE cCRE-pELS | 0.51 | 0.45 | 0.58 | 3.37.E-26 | 0.53 | 0.45 | 0.62 | 5.63.E-15 |
| FraII (aTreg) | CD4 | cOCRs | ENCODE cCRE-dELS | 2.72 | 2.34 | 3.15 | 2.20.E-39 | 3.50 | 2.88 | 4.24 | 3.19.E-37 |
| FraII (aTreg) | CD4 | CAMBUS_control | ENCODE cCRE-dELS | 0.81 | 0.74 | 0.89 | 1.57.E-05 | 0.81 | 0.73 | 0.90 | 8.74.E-05 |
| FraII (aTreg) | CD4 | cOCRs | ENCODE CTCF-only | 0.18 | 0.04 | 0.71 | 1.42.E-02 | 0.23 | 0.06 | 0.92 | 3.84.E-02 |
| FraII (aTreg) | CD4 | CAMBUS_control | ENCODE CTCF-only | 1.90 | 1.22 | 2.95 | 4.53.E-03 | 1.75 | 1.11 | 2.76 | 1.56.E-02 |
| FraII (aTreg) | CD4 | cOCRs | FANTOM enhancers | 5.89 | 5.01 | 6.94 | 9.79.E-101 | 7.81 | 6.46 | 9.45 | 4.01.E-99 |
| FraII (aTreg) | CD4 | CAMBUS_control | FANTOM enhancers | 0.39 | 0.34 | 0.45 | 1.95.E-40 | 0.37 | 0.32 | 0.43 | 8.36.E-38 |
| FraII | CD4 | cOCRs | Super | 5.02 | 3.45 | 7.30 | 3.38.E- | 8.29 | 4.77 | 14.41 | 6.70.E- |

| | | | | | | | | | | | |
|---|---|---|---|---|---|---|---|---|---|---|---|
| (aTreg) | | | enhancers | | | | 17 | | | | 14 |
| FraII (aTreg) | CD4 | CAMBUS_control | Super enhancers | 0.59 | 0.53 | 0.66 | 3.51.E-21 | 0.57 | 0.50 | 0.64 | 9.51.E-22 |
| FraIII | CD4 | cOCRs | ENCODE cCRE-PLS | 4.02 | 3.57 | 4.53 | 3.10.E-115 | 4.22 | 3.55 | 5.00 | 4.63.E-61 |
| FraIII | CD4 | CAMBUS_control | ENCODE cCRE-PLS | 0.32 | 0.28 | 0.35 | 1.78.E-87 | 0.31 | 0.27 | 0.37 | 2.04.E-44 |
| FraIII | CD4 | cOCRs | ENCODE cCRE-pELS | 3.47 | 3.12 | 3.85 | 1.38.E-121 | 3.17 | 2.77 | 3.62 | 4.48.E-63 |
| FraIII | CD4 | CAMBUS_control | ENCODE cCRE-pELS | 0.35 | 0.31 | 0.38 | 1.20.E-100 | 0.38 | 0.33 | 0.43 | 6.38.E-52 |
| FraIII | CD4 | cOCRs | ENCODE cCRE-dELS | 2.64 | 2.40 | 2.91 | 8.80.E-89 | 3.48 | 3.08 | 3.92 | 1.21.E-91 |
| FraIII | CD4 | CAMBUS_control | ENCODE cCRE-dELS | 0.48 | 0.44 | 0.52 | 1.49.E-64 | 0.41 | 0.38 | 0.46 | 1.91.E-66 |
| FraIII | CD4 | cOCRs | ENCODE CTCF-only | 0.36 | 0.21 | 0.61 | 1.53.E-04 | 0.41 | 0.24 | 0.72 | 1.63.E-03 |
| FraIII | CD4 | CAMBUS_control | ENCODE CTCF-only | 2.71 | 1.70 | 4.32 | 2.71.E-05 | 2.39 | 1.48 | 3.86 | 3.74.E-04 |
| FraIII | CD4 | cOCRs | FANTO | 5.09 | 4.59 | 5.64 | 1.06.E- | 6.12 | 5.44 | 6.88 | 1.30.E- |

| | | | | | | | | | | | |
|---|---|---|---|---|---|---|---|---|---|---|---|
| | | | M enhancers | | | | 210 | | | | 200 |
| FraIII | CD4 | CAMBUS_control | FANTOM enhancers | 0.23 | 0.21 | 0.26 | 2.69.E-182 | 0.20 | 0.18 | 0.23 | 3.13.E-173 |
| FraIII | CD4 | cOCRs | Super enhancers | 4.29 | 3.45 | 5.34 | 4.16.E-39 | 5.30 | 4.02 | 6.98 | 2.07.E-32 |
| FraIII | CD4 | CAMBUS_control | Super enhancers | 0.35 | 0.31 | 0.41 | 5.74.E-44 | 0.33 | 0.28 | 0.39 | 6.05.E-40 |
| LAG3 Treg | CD4 | cOCRs | ENCODE cCRE-PLS | 3.93 | 3.52 | 4.38 | 3.85.E-132 | 4.14 | 3.52 | 4.86 | 2.24.E-67 |
| LAG3 Treg | CD4 | CAMBUS_control | ENCODE cCRE-PLS | 0.33 | 0.30 | 0.37 | 2.49.E-95 | 0.34 | 0.29 | 0.39 | 1.20.E-44 |
| LAG3 Treg | CD4 | cOCRs | ENCODE cCRE-pELS | 3.37 | 3.06 | 3.71 | 3.08.E-138 | 3.01 | 2.65 | 3.41 | 6.23.E-66 |
| LAG3 Treg | CD4 | CAMBUS_control | ENCODE cCRE-pELS | 0.37 | 0.33 | 0.40 | 3.80.E-110 | 0.40 | 0.36 | 0.45 | 1.63.E-52 |
| LAG3 Treg | CD4 | cOCRs | ENCODE cCRE-dELS | 2.59 | 2.37 | 2.82 | 1.02.E-102 | 3.34 | 3.00 | 3.73 | 4.52.E-103 |
| LAG3 Treg | CD4 | CAMBUS_contro | ENCODE cCRE- | 0.50 | 0.47 | 0.54 | 8.79.E-71 | 0.45 | 0.41 | 0.49 | 1.03.E-69 |

| | | | | | | | | | | | |
|---|---|---|---|---|---|---|---|---|---|---|---|
| | | l | dELS | | | | | | | | |
| LAG3 Treg | CD4 | cOCRs | ENCODE CTCF-only | 0.36 | 0.23 | 0.57 | 1.75.E-05 | 0.40 | 0.24 | 0.65 | 2.57.E-04 |
| LAG3 Treg | CD4 | CAMBUS_control | ENCODE CTCF-only | 2.21 | 1.53 | 3.20 | 2.46.E-05 | 1.97 | 1.35 | 2.89 | 4.99.E-04 |
| LAG3 Treg | CD4 | cOCRs | FANTOM enhancers | 4.37 | 3.97 | 4.81 | 1.56.E-197 | 5.33 | 4.77 | 5.94 | 1.64.E-196 |
| LAG3 Treg | CD4 | CAMBUS_control | FANTOM enhancers | 0.28 | 0.25 | 0.30 | 3.59.E-167 | 0.24 | 0.21 | 0.26 | 3.69.E-166 |
| LAG3 Treg | CD4 | cOCRs | Super enhancers | 3.65 | 3.04 | 4.39 | 4.06.E-43 | 4.25 | 3.38 | 5.34 | 1.85.E-35 |
| LAG3 Treg | CD4 | CAMBUS_control | Super enhancers | 0.36 | 0.31 | 0.41 | 5.77.E-53 | 0.34 | 0.29 | 0.39 | 8.52.E-47 |
| Memory CD4 | CD4 | cOCRs | ENCODE cCRE-PLS | 4.76 | 4.12 | 5.50 | 1.26.E-99 | 5.11 | 4.16 | 6.28 | 2.71.E-54 |
| Memory CD4 | CD4 | CAMBUS_control | ENCODE cCRE-PLS | 0.30 | 0.26 | 0.34 | 4.43.E-68 | 0.31 | 0.26 | 0.38 | 3.36.E-32 |
| Memory CD4 | CD4 | cOCRs | ENCODE cCRE-pELS | 4.54 | 4.00 | 5.14 | 8.04.E-124 | 4.05 | 3.44 | 4.76 | 1.31.E-63 |

| | | | | | | | | | | |
|---|---|---|---|---|---|---|---|---|---|---|---|
| Memory CD4 | CD4 | CAMBUS_control | ENCODE cCRE-pELS | 0.30 | 0.27 | 0.34 | 2.90.E-91 | 0.34 | 0.29 | 0.40 | 4.93.E-44 |
| Memory CD4 | CD4 | cOCRs | ENCODE cCRE-dELS | 2.70 | 2.41 | 3.03 | 2.12.E-65 | 3.76 | 3.25 | 4.35 | 1.06.E-70 |
| Memory CD4 | CD4 | CAMBUS_control | ENCODE cCRE-dELS | 0.50 | 0.45 | 0.55 | 1.72.E-44 | 0.43 | 0.39 | 0.49 | 6.54.E-47 |
| Memory CD4 | CD4 | cOCRs | ENCODE CTCF-only | 0.34 | 0.16 | 0.72 | 4.63.E-03 | 0.37 | 0.17 | 0.83 | 1.60.E-02 |
| Memory CD4 | CD4 | CAMBUS_control | ENCODE CTCF-only | 2.38 | 1.35 | 4.20 | 2.85.E-03 | 2.17 | 1.20 | 3.94 | 1.07.E-02 |
| Memory CD4 | CD4 | cOCRs | FANTOM enhancers | 5.88 | 5.18 | 6.66 | 3.64.E-169 | 7.54 | 6.53 | 8.71 | 1.21.E-165 |
| Memory CD4 | CD4 | CAMBUS_control | FANTOM enhancers | 0.21 | 0.19 | 0.24 | 8.98.E-147 | 0.18 | 0.16 | 0.21 | 5.80.E-143 |
| Memory CD4 | CD4 | cOCRs | Super enhancers | 5.07 | 3.86 | 6.65 | 1.12.E-31 | 6.86 | 4.80 | 9.81 | 4.74.E-26 |
| Memory CD4 | CD4 | CAMBUS_control | Super enhancers | 0.40 | 0.35 | 0.47 | 2.67.E-33 | 0.40 | 0.34 | 0.47 | 7.46.E-29 |
| Thx | CD4 | cOCRs | ENCOD | 4.21 | 3.74 | 4.75 | 2.18.E- | 4.42 | 3.72 | 5.25 | 8.76.E- |

| | | | | | | | | | | | |
|---|---|---|---|---|---|---|---|---|---|---|---|
| | | | E cCRE-PLS | | | | 121 | | | | 64 |
| Thx | CD4 | CAMBUS_control | ENCODE cCRE-PLS | 0.29 | 0.25 | 0.32 | 8.66.E-99 | 0.28 | 0.24 | 0.33 | 4.54.E-50 |
| Thx | CD4 | cOCRs | ENCODE cCRE-pELS | 3.73 | 3.36 | 4.14 | 1.97.E-136 | 3.24 | 2.83 | 3.72 | 1.34.E-64 |
| Thx | CD4 | CAMBUS_control | ENCODE cCRE-pELS | 0.31 | 0.28 | 0.35 | 5.81.E-116 | 0.35 | 0.31 | 0.40 | 5.85.E-55 |
| Thx | CD4 | cOCRs | ENCODE cCRE-dELS | 2.77 | 2.52 | 3.05 | 1.60.E-98 | 3.59 | 3.19 | 4.05 | 5.17.E-98 |
| Thx | CD4 | CAMBUS_control | ENCODE cCRE-dELS | 0.43 | 0.39 | 0.47 | 4.69.E-81 | 0.37 | 0.33 | 0.41 | 1.05.E-79 |
| Thx | CD4 | cOCRs | ENCODE CTCF-only | 0.40 | 0.24 | 0.67 | 5.08.E-04 | 0.47 | 0.27 | 0.79 | 4.62.E-03 |
| Thx | CD4 | CAMBUS_control | ENCODE CTCF-only | 2.56 | 1.59 | 4.14 | 1.18.E-04 | 2.25 | 1.37 | 3.70 | 1.28.E-03 |
| Thx | CD4 | cOCRs | FANTOM enhancers | 5.10 | 4.60 | 5.66 | 2.16.E-208 | 6.02 | 5.35 | 6.78 | 4.26.E-195 |
| Thx | CD4 | CAMBUS_control | FANTOM enhance | 0.22 | 0.20 | 0.24 | 1.52.E-197 | 0.19 | 0.17 | 0.21 | 2.62.E-185 |

| | | | | | | | | | | | |
|---|---|---|---|---|---|---|---|---|---|---|---|
| | | | rs | | | | | | | | |
| Thx | CD4 | cOCRs | Super enhancers | 4.23 | 3.42 | 5.23 | 2.12.E-40 | 5.26 | 4.02 | 6.88 | 8.90.E-34 |
| Thx | CD4 | CAMBUS_control | Super enhancers | 0.32 | 0.27 | 0.37 | 1.80.E-45 | 0.29 | 0.24 | 0.35 | 1.78.E-40 |
| Naive CD8 | CD8 | cOCRs | ENCODE cCRE-PLS | 4.21 | 2.47 | 7.18 | 1.33.E-07 | 13.17 | 6.64 | 26.11 | 1.58.E-13 |
| Naive CD8 | CD8 | CAMBUS_control | ENCODE cCRE-PLS | 0.70 | 0.45 | 1.09 | 1.12.E-01 | 0.46 | 0.27 | 0.78 | 4.06.E-03 |
| Naive CD8 | CD8 | cOCRs | ENCODE cCRE-pELS | 4.39 | 2.68 | 7.16 | 3.54.E-09 | 6.01 | 3.16 | 11.46 | 4.93.E-08 |
| Naive CD8 | CD8 | CAMBUS_control | ENCODE cCRE-pELS | 0.62 | 0.42 | 0.92 | 1.73.E-02 | 0.60 | 0.36 | 1.01 | 5.42.E-02 |
| Naive CD8 | CD8 | cOCRs | ENCODE cCRE-dELS | 2.05 | 1.30 | 3.25 | 2.14.E-03 | 2.09 | 1.13 | 3.87 | 1.89.E-02 |
| Naive CD8 | CD8 | CAMBUS_control | ENCODE cCRE-dELS | 0.84 | 0.60 | 1.18 | 3.06.E-01 | 0.97 | 0.65 | 1.44 | 8.76.E-01 |
| Naive CD8 | CD8 | cOCRs | ENCODE CTCF-only | 0.00 | 0.00 | 6.E+47 | 8.83.E-01 | 0.00 | 0.00 | 3.E+64 | 9.09.E-01 |
| Naive CD8 | CD8 | CAMBUS_contro | ENCODE CTCF- | 23440.47 | 0.00 | 1.E+56 | 8.69.E-01 | 26058.47 | 0.00 | 4.E+61 | 8.80.E-01 |

| | | | | | | | | | | | |
|---|---|---|---|---|---|---|---|---|---|---|---|
| | | l | only | | | | | | | | |
| Naive CD8 | CD8 | cOCRs | FANTOM enhancers | 5.26 | 3.12 | 8.86 | 4.53.E-10 | 10.40 | 5.58 | 19.37 | 1.61.E-13 |
| Naive CD8 | CD8 | CAMBUS_control | FANTOM enhancers | 0.31 | 0.20 | 0.50 | 1.27.E-06 | 0.24 | 0.14 | 0.39 | 8.37.E-09 |
| Naive CD8 | CD8 | cOCRs | Super enhancers | 3.71 | 1.15 | 11.93 | 2.81.E-02 | 2.53 | 0.77 | 8.33 | 1.28.E-01 |
| Naive CD8 | CD8 | CAMBUS_control | Super enhancers | 1.42 | 1.04 | 1.93 | 2.81.E-02 | 1.54 | 1.11 | 2.14 | 9.73.E-03 |
| CD8+effector | CD8 | cOCRs | ENCODE cCRE-PLS | 4.87 | 3.73 | 6.35 | 1.46.E-31 | 6.42 | 4.43 | 9.30 | 7.67.E-23 |
| CD8+effector | CD8 | CAMBUS_control | ENCODE cCRE-PLS | 0.31 | 0.24 | 0.39 | 2.43.E-21 | 0.28 | 0.20 | 0.39 | 3.76.E-13 |
| CD8+effector | CD8 | cOCRs | ENCODE cCRE-pELS | 4.93 | 3.89 | 6.24 | 6.17.E-40 | 4.27 | 3.11 | 5.86 | 3.14.E-19 |
| CD8+effector | CD8 | CAMBUS_control | ENCODE cCRE-pELS | 0.29 | 0.24 | 0.37 | 9.50.E-29 | 0.35 | 0.26 | 0.47 | 3.88.E-12 |
| CD8+effector | CD8 | cOCRs | ENCODE cCRE-dELS | 2.69 | 2.16 | 3.34 | 5.96.E-19 | 3.74 | 2.80 | 5.00 | 5.17.E-19 |

| | | | | | | | | | | | |
|---|---|---|---|---|---|---|---|---|---|---|---|
| CD8+effector | CD8 | CAMBUS_control | ENCODE cCRE-dELS | 0.50 | 0.41 | 0.60 | 9.10.E-13 | 0.44 | 0.35 | 0.56 | 4.97.E-12 |
| CD8+effector | CD8 | cOCRs | ENCODE CTCF-only | 0.00 | 0.00 | 1.0.E+61 | 8.85.E-01 | 0.00 | 0.00 | 1.E+79 | 9.09.E-01 |
| CD8+effector | CD8 | CAMBUS_control | ENCODE CTCF-only | 1.99 | 0.64 | 6.21 | 2.35.E-01 | 1.58 | 0.51 | 4.93 | 4.31.E-01 |
| CD8+effector | CD8 | cOCRs | FANTOM enhancers | 7.25 | 5.74 | 9.16 | 3.40.E-62 | 9.82 | 7.39 | 13.05 | 7.33.E-56 |
| CD8+effector | CD8 | CAMBUS_control | FANTOM enhancers | 0.17 | 0.13 | 0.21 | 7.11.E-56 | 0.14 | 0.11 | 0.18 | 1.03.E-49 |
| CD8+effector | CD8 | cOCRs | Super enhancers | 7.53 | 3.87 | 14.64 | 2.63.E-09 | 12.20 | 4.51 | 33.02 | 8.51.E-07 |
| CD8+effector | CD8 | CAMBUS_control | Super enhancers | 0.49 | 0.37 | 0.64 | 3.20.E-07 | 0.49 | 0.36 | 0.67 | 5.32.E-06 |
| CD8+central memory | CD8 | cOCRs | ENCODE cCRE-PLS | 4.74 | 3.79 | 5.94 | 6.52.E-42 | 5.91 | 4.31 | 8.12 | 3.61.E-28 |
| CD8+central memory | CD8 | CAMBUS_control | ENCODE cCRE-PLS | 0.37 | 0.30 | 0.45 | 8.92.E-22 | 0.37 | 0.28 | 0.50 | 5.59.E-11 |
| CD8+ce | CD8 | cOCRs | ENCOD | 4.82 | 3.95 | 5.87 | 1.48.E- | 4.74 | 3.66 | 6.13 | 2.30.E- |

| | | | | | | | | | | | |
|---|---|---|---|---|---|---|---|---|---|---|---|
| ntral memory | | | E cCRE-pELS | | | | 54 | | | | 32 |
| CD8+central memory | CD8 | CAMBUS_control | ENCODE cCRE-pELS | 0.35 | 0.29 | 0.42 | 2.36.E-32 | 0.37 | 0.30 | 0.47 | 4.15.E-17 |
| CD8+central memory | CD8 | cOCRs | ENCODE cCRE-dELS | 2.77 | 2.31 | 3.31 | 1.77.E-28 | 3.61 | 2.85 | 4.57 | 1.05.E-26 |
| CD8+central memory | CD8 | CAMBUS_control | ENCODE cCRE-dELS | 0.54 | 0.47 | 0.63 | 2.60.E-16 | 0.53 | 0.44 | 0.62 | 7.42.E-14 |
| CD8+central memory | CD8 | cOCRs | ENCODE CTCF-only | 0.15 | 0.02 | 1.06 | 5.73.E-02 | 0.20 | 0.03 | 1.40 | 1.05.E-01 |
| CD8+central memory | CD8 | CAMBUS_control | ENCODE CTCF-only | 1.15 | 0.62 | 2.15 | 6.58.E-01 | 0.96 | 0.52 | 1.80 | 9.09.E-01 |
| CD8+central memory | CD8 | cOCRs | FANTOM enhancers | 6.78 | 5.58 | 8.23 | 1.09.E-82 | 9.50 | 7.54 | 11.98 | 4.29.E-81 |
| CD8+central memory | CD8 | CAMBUS_control | FANTOM enhancers | 0.21 | 0.18 | 0.26 | 1.14.E-63 | 0.18 | 0.15 | 0.22 | 6.91.E-62 |
| CD8+central memory | CD8 | cOCRs | Super enhancers | 5.80 | 3.60 | 9.33 | 4.83.E-13 | 11.39 | 5.35 | 24.27 | 2.90.E-10 |
| CD8+central | CD8 | CAMBUS_contro | Super enhance | 0.49 | 0.40 | 0.59 | 5.48.E-13 | 0.48 | 0.39 | 0.59 | 4.47.E-12 |

| | | | | | | | | | | | |
|---|---|---|---|---|---|---|---|---|---|---|---|
| memory | | l | rs | | | | | | | | |
| CD8+effector memory | CD8 | cOCRs | ENCODE cCRE-PLS | 5.01 | 3.78 | 6.63 | 2.36.E-29 | 7.27 | 4.90 | 10.77 | 5.35.E-23 |
| CD8+effector memory | CD8 | CAMBUS_control | ENCODE cCRE-PLS | 0.47 | 0.37 | 0.60 | 4.89.E-10 | 0.46 | 0.33 | 0.65 | 7.99.E-06 |
| CD8+effector memory | CD8 | cOCRs | ENCODE cCRE-pELS | 4.70 | 3.64 | 6.05 | 9.45.E-33 | 4.28 | 3.03 | 6.07 | 2.56.E-16 |
| CD8+effector memory | CD8 | CAMBUS_control | ENCODE cCRE-pELS | 0.48 | 0.39 | 0.59 | 4.92.E-12 | 0.57 | 0.42 | 0.77 | 2.16.E-04 |
| CD8+effector memory | CD8 | cOCRs | ENCODE cCRE-dELS | 2.64 | 2.09 | 3.34 | 4.77.E-16 | 3.68 | 2.66 | 5.09 | 3.08.E-15 |
| CD8+effector memory | CD8 | CAMBUS_control | ENCODE cCRE-dELS | 0.66 | 0.56 | 0.79 | 2.12.E-06 | 0.65 | 0.54 | 0.79 | 1.72.E-05 |
| CD8+effector memory | CD8 | cOCRs | ENCODE CTCF-only | 0.00 | 0.00 | 4.E+65 | 8.93.E-01 | 0.00 | 0.00 | 6.E+87 | 9.17.E-01 |
| CD8+effector memory | CD8 | CAMBUS_control | ENCODE CTCF-only | 2.25 | 0.84 | 6.03 | 1.06.E-01 | 2.55 | 0.82 | 7.93 | 1.06.E-01 |
| CD8+effector memory | CD8 | cOCRs | FANTOM enhancers | 6.73 | 5.23 | 8.66 | 1.01.E-49 | 10.38 | 7.59 | 14.18 | 8.30.E-49 |
| CD8+eff | CD8 | CAMBU | FANTO | 0.27 | 0.21 | 0.33 | 3.34.E- | 0.23 | 0.18 | 0.30 | 4.25.E- |

| | | | | | | | | | | | |
|---|---|---|---|---|---|---|---|---|---|---|---|
| ector memory | | S_control | M enhancers | | | | 31 | | | | 30 |
| CD8+effector memory | CD8 | cOCRs | Super enhancers | 5.36 | 2.84 | 10.13 | 2.27.E-07 | 9.03 | 3.35 | 24.34 | 1.36.E-05 |
| CD8+effector memory | CD8 | CAMBUS_control | Super enhancers | 0.66 | 0.54 | 0.79 | 1.44.E-05 | 0.63 | 0.51 | 0.77 | 1.03.E-05 |
| Naive B | B | cOCRs | ENCODE cCRE-PLS | 4.21 | 2.26 | 7.83 | 5.68.E-06 | 5.77 | 2.44 | 13.61 | 6.36.E-05 |
| Naive B | B | CAMBUS_control | ENCODE cCRE-PLS | 1.33 | 0.81 | 2.17 | 2.58.E-01 | 1.43 | 0.72 | 2.86 | 3.09.E-01 |
| Naive B | B | cOCRs | ENCODE cCRE-pELS | 5.06 | 2.89 | 8.85 | 1.28.E-08 | 6.24 | 3.05 | 12.79 | 5.60.E-07 |
| Naive B | B | CAMBUS_control | ENCODE cCRE-pELS | 0.82 | 0.54 | 1.25 | 3.63.E-01 | 0.76 | 0.44 | 1.30 | 3.12.E-01 |
| Naive B | B | cOCRs | ENCODE cCRE-dELS | 2.04 | 1.21 | 3.45 | 7.24.E-03 | 2.30 | 1.15 | 4.61 | 1.87.E-02 |
| Naive B | B | CAMBUS_control | ENCODE cCRE-dELS | 0.77 | 0.54 | 1.12 | 1.73.E-01 | 0.87 | 0.57 | 1.31 | 4.95.E-01 |
| Naive B | B | cOCRs | ENCODE CTCF-only | 0.00 | 0.00 | 5.E+54 | 8.97.E-01 | 0.00 | 0.00 | 9.E+72 | 9.20.E-01 |

| | | | | | | | | | | | |
|---|---|---|---|---|---|---|---|---|---|---|---|
| Naive B | B | CAMBUS_control | ENCODE CTCF-only | 0.54 | 0.20 | 1.47 | 2.29.E-01 | 0.52 | 0.19 | 1.40 | 1.93.E-01 |
| Naive B | B | cOCRs | FANTOM enhancers | 5.73 | 3.18 | 10.34 | 6.70.E-09 | 8.11 | 3.95 | 16.65 | 1.13.E-08 |
| Naive B | B | CAMBUS_control | FANTOM enhancers | 0.32 | 0.19 | 0.53 | 1.49.E-05 | 0.30 | 0.17 | 0.54 | 5.38.E-05 |
| Naive B | B | cOCRs | Super enhancers | 14969.53 | 0.00 | 3.E+24 | 6.86.E-01 | 17825.03 | 0.00 | 3.E+30 | 7.50.E-01 |
| Naive B | B | CAMBUS_control | Super enhancers | 1.30 | 0.96 | 1.77 | 9.30.E-02 | 1.33 | 0.97 | 1.83 | 7.56.E-02 |
| unsw memory B | B | cOCRs | ENCODE cCRE-PLS | 4.43 | 3.37 | 5.82 | 1.91.E-26 | 6.35 | 4.24 | 9.51 | 3.21.E-19 |
| unsw memory B | B | CAMBUS_control | ENCODE cCRE-PLS | 0.34 | 0.26 | 0.44 | 6.04.E-17 | 0.31 | 0.21 | 0.45 | 7.37.E-10 |
| unsw memory B | B | cOCRs | ENCODE cCRE-pELS | 4.26 | 3.33 | 5.44 | 6.57.E-31 | 4.39 | 3.12 | 6.16 | 1.41.E-17 |
| unsw memory B | B | CAMBUS_control | ENCODE cCRE-pELS | 0.32 | 0.25 | 0.40 | 5.18.E-23 | 0.33 | 0.24 | 0.45 | 4.89.E-12 |
| unsw | B | cOCRs | ENCOD | 2.82 | 2.25 | 3.54 | 1.91.E- | 3.75 | 2.74 | 5.13 | 1.39.E- |

| | | | | | | | | | | | |
|---|---|---|---|---|---|---|---|---|---|---|---|
| memory B | | | E cCRE-dELS | | | | 19 | | | | 16 |
| unsw memory B | B | CAMBUS_control | ENCODE cCRE-dELS | 0.39 | 0.32 | 0.48 | 3.98.E-18 | 0.34 | 0.26 | 0.45 | 6.49.E-15 |
| unsw memory B | B | cOCRs | ENCODE CTCF-only | 0.00 | 0.00 | 4.E+63 | 8.91.E-01 | 0.00 | 0.00 | 6.E+50 | 8.75.E-01 |
| unsw memory B | B | CAMBUS_control | ENCODE CTCF-only | 4.29 | 0.60 | 30.55 | 1.46.E-01 | 3.17 | 0.45 | 22.51 | 2.48.E-01 |
| unsw memory B | B | cOCRs | FANTOM enhancers | 5.72 | 4.44 | 7.37 | 2.44.E-41 | 10.09 | 7.42 | 13.72 | 4.01.E-49 |
| unsw memory B | B | CAMBUS_control | FANTOM enhancers | 0.19 | 0.15 | 0.24 | 4.18.E-39 | 0.12 | 0.09 | 0.16 | 1.00.E-46 |
| unsw memory B | B | cOCRs | Super enhancers | 6.62 | 3.40 | 12.89 | 2.77.E-08 | 7.87 | 3.22 | 19.22 | 6.02.E-06 |
| unsw memory B | B | CAMBUS_control | Super enhancers | 0.62 | 0.46 | 0.84 | 1.74.E-03 | 0.75 | 0.54 | 1.04 | 8.16.E-02 |
| sw memory B | B | cOCRs | ENCODE cCRE-PLS | 5.12 | 4.07 | 6.44 | 2.37.E-44 | 6.40 | 4.61 | 8.88 | 1.46.E-28 |
| sw memory | B | CAMBUS_contro | ENCODE cCRE- | 0.30 | 0.25 | 0.38 | 1.22.E-27 | 0.30 | 0.22 | 0.41 | 3.46.E-14 |

| | | | | | | | | | | | |
|---|---|---|---|---|---|---|---|---|---|---|---|
| B | | l | PLS | | | | | | | | |
| sw memory B | B | cOCRs | ENCODE cCRE-pELS | 4.70 | 3.83 | 5.78 | 3.43.E-49 | 4.48 | 3.39 | 5.92 | 4.27.E-26 |
| sw memory B | B | CAMBUS_control | ENCODE cCRE-pELS | 0.28 | 0.23 | 0.34 | 1.41.E-38 | 0.31 | 0.24 | 0.40 | 2.75.E-19 |
| sw memory B | B | cOCRs | ENCODE cCRE-dELS | 2.58 | 2.14 | 3.13 | 1.49.E-22 | 3.30 | 2.56 | 4.26 | 5.49.E-20 |
| sw memory B | B | CAMBUS_control | ENCODE cCRE-dELS | 0.43 | 0.36 | 0.52 | 2.12.E-20 | 0.39 | 0.31 | 0.48 | 2.84.E-17 |
| sw memory B | B | cOCRs | ENCODE CTCF-only | 0.00 | 0.00 | 6.E+52 | 8.70.E-01 | 0.00 | 0.00 | 1.E+70 | 8.99.E-01 |
| sw memory B | B | CAMBUS_control | ENCODE CTCF-only | 3.25 | 0.81 | 12.95 | 9.54.E-02 | 2.44 | 0.61 | 9.79 | 2.09.E-01 |
| sw memory B | B | cOCRs | FANTOM enhancers | 6.00 | 4.84 | 7.42 | 8.02.E-61 | 9.69 | 7.50 | 12.52 | 9.08.E-68 |
| sw memory B | B | CAMBUS_control | FANTOM enhancers | 0.19 | 0.15 | 0.23 | 2.45.E-56 | 0.13 | 0.10 | 0.16 | 9.86.E-63 |
| sw memory B | B | cOCRs | Super enhancers | 7.62 | 4.19 | 13.88 | 3.08.E-11 | 11.40 | 4.68 | 27.77 | 8.39.E-08 |

| | | | | | | | | | | | |
|---|---|---|---|---|---|---|---|---|---|---|---|
| sw memory B | B | CAMBUS_control | Super enhancers | 0.53 | 0.41 | 0.68 | 1.06.E-06 | 0.60 | 0.45 | 0.79 | 3.29.E-04 |
| DN B | B | cOCRs | ENCODE cCRE-PLS | 4.82 | 3.96 | 5.86 | 9.88.E-56 | 6.50 | 4.93 | 8.58 | 6.42.E-40 |
| DN B | B | CAMBUS_control | ENCODE cCRE-PLS | 0.26 | 0.22 | 0.32 | 1.75.E-43 | 0.22 | 0.17 | 0.29 | 1.33.E-27 |
| DN B | B | cOCRs | ENCODE cCRE-pELS | 4.60 | 3.86 | 5.47 | 3.60.E-66 | 4.86 | 3.86 | 6.11 | 1.93.E-41 |
| DN B | B | CAMBUS_control | ENCODE cCRE-pELS | 0.26 | 0.22 | 0.30 | 3.32.E-57 | 0.25 | 0.20 | 0.31 | 6.30.E-35 |
| DN B | B | cOCRs | ENCODE cCRE-dELS | 2.64 | 2.25 | 3.09 | 7.35.E-33 | 3.42 | 2.77 | 4.22 | 4.02.E-30 |
| DN B | B | CAMBUS_control | ENCODE cCRE-dELS | 0.40 | 0.35 | 0.47 | 1.96.E-30 | 0.34 | 0.28 | 0.41 | 3.16.E-27 |
| DN B | B | cOCRs | ENCODE CTCF-only | 0.28 | 0.07 | 1.11 | 7.00.E-02 | 0.38 | 0.10 | 1.54 | 1.77.E-01 |
| DN B | B | CAMBUS_control | ENCODE CTCF-only | 3.99 | 0.99 | 16.00 | 5.09.E-02 | 3.00 | 0.75 | 12.03 | 1.20.E-01 |
| DN B | B | cOCRs | FANTOM enhance | 5.58 | 4.66 | 6.69 | 2.17.E-77 | 8.76 | 7.07 | 10.84 | 3.37.E-88 |

| | | | | | | | | | | | |
|---|---|---|---|---|---|---|---|---|---|---|---|
| | | | rs | | | | | | | | |
| DN B | B | CAMBUS_control | FANTOM enhancers | 0.19 | 0.15 | 0.22 | 1.39.E-75 | 0.12 | 0.10 | 0.15 | 2.82.E-86 |
| DN B | B | cOCRs | Super enhancers | 7.80 | 4.74 | 12.83 | 6.03.E-16 | 14.53 | 6.47 | 32.65 | 9.26.E-11 |
| DN B | B | CAMBUS_control | Super enhancers | 0.38 | 0.29 | 0.50 | 2.03.E-12 | 0.41 | 0.30 | 0.55 | 6.33.E-09 |
| plasmablast | B | cOCRs | ENCODE cCRE-PLS | 3.51 | 3.09 | 3.98 | 2.64.E-83 | 3.95 | 3.29 | 4.75 | 2.46.E-49 |
| plasmablast | B | CAMBUS_control | ENCODE cCRE-PLS | 0.37 | 0.33 | 0.42 | 1.36.E-60 | 0.35 | 0.30 | 0.42 | 3.72.E-33 |
| plasmablast | B | cOCRs | ENCODE cCRE-pELS | 3.62 | 3.25 | 4.04 | 1.86.E-118 | 3.40 | 2.95 | 3.93 | 2.52.E-62 |
| plasmablast | B | CAMBUS_control | ENCODE cCRE-pELS | 0.37 | 0.33 | 0.41 | 7.46.E-88 | 0.40 | 0.35 | 0.46 | 3.01.E-42 |
| plasmablast | B | cOCRs | ENCODE cCRE-dELS | 2.61 | 2.36 | 2.89 | 1.74.E-78 | 3.16 | 2.78 | 3.59 | 1.64.E-69 |
| plasmablast | B | CAMBUS_control | ENCODE cCRE-dELS | 0.57 | 0.52 | 0.62 | 3.62.E-41 | 0.55 | 0.50 | 0.61 | 5.19.E-34 |
| plasmabl | B | cOCRs | ENCOD | 0.54 | 0.32 | 0.92 | 2.37.E- | 0.62 | 0.35 | 1.09 | 9.79.E- |

| | | | | | | | | | | | |
|---|---|---|---|---|---|---|---|---|---|---|---|
| ast | | | E CTCF-only | | | | 02 | | | | 02 |
| plasmablast | B | CAMBUS_control | ENCODE CTCF-only | 1.67 | 1.13 | 2.49 | 1.08.E-02 | 1.64 | 1.07 | 2.53 | 2.46.E-02 |
| plasmablast | B | cOCRs | FANTOM enhancers | 4.34 | 3.86 | 4.89 | 6.68.E-131 | 5.43 | 4.74 | 6.22 | 2.60.E-132 |
| plasmablast | B | CAMBUS_control | FANTOM enhancers | 0.31 | 0.28 | 0.35 | 1.29.E-94 | 0.28 | 0.24 | 0.31 | 3.36.E-93 |
| plasmablast | B | cOCRs | Super enhancers | 4.11 | 3.26 | 5.18 | 9.35.E-33 | 5.30 | 3.89 | 7.23 | 6.62.E-26 |
| plasmablast | B | CAMBUS_control | Super enhancers | 0.40 | 0.35 | 0.46 | 3.33.E-40 | 0.37 | 0.32 | 0.43 | 1.01.E-36 |
| plasmacytoid DC | DC | cOCRs | ENCODE cCRE-PLS | 2.27 | 1.80 | 2.85 | 2.58.E-12 | 2.73 | 2.00 | 3.73 | 2.13.E-10 |
| plasmacytoid DC | DC | CAMBUS_control | ENCODE cCRE-PLS | 0.49 | 0.39 | 0.61 | 5.05.E-10 | 0.40 | 0.29 | 0.54 | 5.99.E-09 |
| plasmacytoid DC | DC | cOCRs | ENCODE cCRE-pELS | 1.80 | 1.48 | 2.19 | 3.53.E-09 | 1.83 | 1.43 | 2.35 | 1.56.E-06 |
| plasmacytoid DC | DC | CAMBUS_contro | ENCODE cCRE- | 0.56 | 0.46 | 0.67 | 1.65.E-09 | 0.53 | 0.42 | 0.68 | 3.34.E-07 |

| | | | | | | | | | | |
|---|---|---|---|---|---|---|---|---|---|---|
| | | l | pELS | | | | | | | |
| plasmacytoid DC | DC | cOCRs | ENCODE cCRE-dELS | 1.99 | 1.70 | 2.34 | 2.47.E-17 | 2.20 | 1.82 | 2.65 | 3.77.E-16 |
| plasmacytoid DC | DC | CAMBUS_control | ENCODE cCRE-dELS | 0.55 | 0.47 | 0.64 | 1.41.E-14 | 0.49 | 0.41 | 0.59 | 2.47.E-14 |
| plasmacytoid DC | DC | cOCRs | ENCODE CTCF-only | 0.54 | 0.27 | 1.09 | 8.59.E-02 | 0.48 | 0.21 | 1.07 | 7.29.E-02 |
| plasmacytoid DC | DC | CAMBUS_control | ENCODE CTCF-only | 1.92 | 0.95 | 3.86 | 6.75.E-02 | 2.16 | 0.96 | 4.84 | 6.11.E-02 |
| plasmacytoid DC | DC | cOCRs | FANTOM enhancers | 3.75 | 3.13 | 4.49 | 3.39.E-47 | 4.45 | 3.65 | 5.43 | 2.21.E-49 |
| plasmacytoid DC | DC | CAMBUS_control | FANTOM enhancers | 0.28 | 0.23 | 0.33 | 9.90.E-46 | 0.23 | 0.19 | 0.28 | 2.32.E-48 |
| plasmacytoid DC | DC | cOCRs | Super enhancers | 1.57 | 1.20 | 2.05 | 9.09.E-04 | 1.43 | 1.07 | 1.91 | 1.50.E-02 |
| plasmacytoid DC | DC | CAMBUS_control | Super enhancers | 0.64 | 0.49 | 0.83 | 6.67.E-04 | 0.69 | 0.52 | 0.92 | 1.12.E-02 |
| myeloid DC | DC | cOCRs | ENCODE cCRE-PLS | 2.98 | 2.37 | 3.74 | 9.77.E-21 | 3.03 | 2.16 | 4.26 | 1.81.E-10 |

| | | | | | | | | | | | |
|---|---|---|---|---|---|---|---|---|---|---|---|
| myeloid DC | DC | CAMBUS_control | ENCODE cCRE-PLS | 0.51 | 0.41 | 0.64 | 7.28.E-09 | 0.50 | 0.35 | 0.69 | 4.56.E-05 |
| myeloid DC | DC | cOCRs | ENCODE cCRE-pELS | 2.54 | 2.12 | 3.06 | 2.70.E-23 | 2.36 | 1.87 | 2.99 | 9.79.E-13 |
| myeloid DC | DC | CAMBUS_control | ENCODE cCRE-pELS | 0.42 | 0.35 | 0.50 | 6.01.E-23 | 0.40 | 0.32 | 0.49 | 3.46.E-17 |
| myeloid DC | DC | cOCRs | ENCODE cCRE-dELS | 2.44 | 2.08 | 2.85 | 7.14.E-29 | 2.77 | 2.30 | 3.35 | 4.29.E-26 |
| myeloid DC | DC | CAMBUS_control | ENCODE cCRE-dELS | 0.47 | 0.41 | 0.55 | 3.03.E-24 | 0.42 | 0.35 | 0.50 | 6.86.E-23 |
| myeloid DC | DC | cOCRs | ENCODE CTCF-only | 0.14 | 0.03 | 0.54 | 4.79.E-03 | 0.17 | 0.04 | 0.66 | 1.09.E-02 |
| myeloid DC | DC | CAMBUS_control | ENCODE CTCF-only | 8.20 | 2.05 | 32.87 | 2.96.E-03 | 6.78 | 1.70 | 27.01 | 6.63.E-03 |
| myeloid DC | DC | cOCRs | FANTOM enhancers | 4.68 | 3.98 | 5.49 | 3.08.E-79 | 5.37 | 4.48 | 6.44 | 2.56.E-74 |
| myeloid DC | DC | CAMBUS_control | FANTOM enhancers | 0.25 | 0.21 | 0.29 | 4.02.E-71 | 0.21 | 0.18 | 0.25 | 1.04.E-69 |
| myeloid | DC | cOCRs | Super | 2.60 | 1.92 | 3.50 | 4.17.E- | 2.51 | 1.79 | 3.50 | 7.27.E- |

| | | | | | | | | | | | |
|---|---|---|---|---|---|---|---|---|---|---|---|
| DC | | | enhancers | | | | 10 | | | | 08 |
| myeloid DC | DC | CAMBUS_control | Super enhancers | 0.47 | 0.37 | 0.60 | 4.38.E-09 | 0.51 | 0.38 | 0.67 | 1.34.E-06 |
| monocyte CD16+ | Mono | cOCRs | ENCODE cCRE-PLS | 3.70 | 3.28 | 4.18 | 9.90.E-101 | 3.70 | 3.10 | 4.40 | 9.43.E-49 |
| monocyte CD16+ | Mono | CAMBUS_control | ENCODE cCRE-PLS | 0.32 | 0.29 | 0.36 | 7.16.E-79 | 0.34 | 0.28 | 0.40 | 3.51.E-35 |
| monocyte CD16+ | Mono | cOCRs | ENCODE cCRE-pELS | 3.41 | 3.10 | 3.75 | 9.85.E-137 | 3.12 | 2.76 | 3.53 | 6.11.E-74 |
| monocyte CD16+ | Mono | CAMBUS_control | ENCODE cCRE-pELS | 0.32 | 0.29 | 0.35 | 1.05.E-122 | 0.35 | 0.31 | 0.39 | 1.18.E-65 |
| monocyte CD16+ | Mono | cOCRs | ENCODE cCRE-dELS | 2.35 | 2.16 | 2.55 | 4.48.E-90 | 2.67 | 2.42 | 2.94 | 6.07.E-85 |
| monocyte CD16+ | Mono | CAMBUS_control | ENCODE cCRE-dELS | 0.44 | 0.40 | 0.47 | 1.85.E-88 | 0.39 | 0.36 | 0.43 | 2.58.E-83 |
| monocyte CD16+ | Mono | cOCRs | ENCODE CTCF-only | 0.43 | 0.27 | 0.68 | 3.48.E-04 | 0.46 | 0.28 | 0.76 | 2.35.E-03 |
| monocyte CD16+ | Mono | CAMBUS_control | ENCODE CTCF-only | 2.08 | 1.35 | 3.20 | 8.62.E-04 | 1.90 | 1.21 | 3.00 | 5.43.E-03 |
| monocyt | Mono | cOCRs | FANTO | 4.33 | 3.95 | 4.75 | 2.48.E- | 4.81 | 4.34 | 5.33 | 4.41.E- |

| | | | | | | | | | | | |
|---|---|---|---|---|---|---|---|---|---|---|---|
| e CD16+ | | | M enhancers | | | | 210 | | | | 196 |
| monocyte CD16+ | Mono | CAMBUS_control | FANTOM enhancers | 0.24 | 0.22 | 0.26 | 1.04.E-204 | 0.22 | 0.20 | 0.24 | 1.62.E-190 |
| monocyte CD16+ | Mono | cOCRs | Super enhancers | 3.27 | 2.78 | 3.85 | 7.47.E-46 | 3.23 | 2.69 | 3.88 | 3.79.E-36 |
| monocyte CD16+ | Mono | CAMBUS_control | Super enhancers | 0.40 | 0.35 | 0.46 | 1.51.E-39 | 0.42 | 0.36 | 0.48 | 1.32.E-30 |
| monocyte CD16- | Mono | cOCRs | ENCODE cCRE-PLS | 4.20 | 3.64 | 4.84 | 2.32.E-87 | 4.24 | 3.44 | 5.23 | 8.75.E-42 |
| monocyte CD16- | Mono | CAMBUS_control | ENCODE cCRE-PLS | 0.30 | 0.27 | 0.35 | 1.37.E-64 | 0.33 | 0.27 | 0.41 | 1.14.E-26 |
| monocyte CD16- | Mono | cOCRs | ENCODE cCRE-pELS | 4.30 | 3.83 | 4.84 | 1.49.E-131 | 3.96 | 3.40 | 4.62 | 3.18.E-69 |
| monocyte CD16- | Mono | CAMBUS_control | ENCODE cCRE-pELS | 0.27 | 0.24 | 0.30 | 9.57.E-115 | 0.29 | 0.25 | 0.34 | 1.04.E-58 |
| monocyte CD16- | Mono | cOCRs | ENCODE cCRE-dELS | 2.83 | 2.55 | 3.15 | 3.37.E-85 | 3.80 | 3.33 | 4.34 | 1.32.E-85 |
| monocyte CD16- | Mono | CAMBUS_contro | ENCODE cCRE- | 0.36 | 0.33 | 0.40 | 1.50.E-84 | 0.29 | 0.25 | 0.32 | 4.11.E-85 |

| | | | | | | | | | | | |
|---|---|---|---|---|---|---|---|---|---|---|---|
| | | l | dELS | | | | | | | | |
| monocyte CD16- | Mono | cOCRs | ENCODE CTCF-only | 0.36 | 0.17 | 0.75 | 6.30.E-03 | 0.41 | 0.18 | 0.91 | 2.93.E-02 |
| monocyte CD16- | Mono | CAMBUS_control | ENCODE CTCF-only | 2.38 | 1.23 | 4.58 | 9.68.E-03 | 2.06 | 1.03 | 4.14 | 4.15.E-02 |
| monocyte CD16- | Mono | cOCRs | FANTOM enhancers | 5.52 | 4.91 | 6.21 | 4.89.E-179 | 6.89 | 6.03 | 7.88 | 3.25.E-174 |
| monocyte CD16- | Mono | CAMBUS_control | FANTOM enhancers | 0.19 | 0.17 | 0.21 | 6.06.E-175 | 0.15 | 0.13 | 0.17 | 5.01.E-170 |
| monocyte CD16- | Mono | cOCRs | Super enhancers | 6.69 | 5.07 | 8.81 | 1.90.E-41 | 8.08 | 5.70 | 11.45 | 8.72.E-32 |
| monocyte CD16- | Mono | CAMBUS_control | Super enhancers | 0.31 | 0.26 | 0.37 | 6.66.E-38 | 0.32 | 0.26 | 0.39 | 1.94.E-29 |
| nonclassical Monocyte | Mono | cOCRs | ENCODE cCRE-PLS | 3.12 | 2.68 | 3.63 | 5.42.E-49 | 3.10 | 2.49 | 3.87 | 7.80.E-24 |
| nonclassical Monocyte | Mono | CAMBUS_control | ENCODE cCRE-PLS | 0.41 | 0.35 | 0.48 | 3.11.E-31 | 0.43 | 0.34 | 0.53 | 2.45.E-14 |
| nonclass | Mono | cOCRs | ENCOD | 2.82 | 2.50 | 3.18 | 2.45.E- | 2.51 | 2.15 | 2.92 | 1.15.E- |

| | | | | | | | | | | | |
|---|---|---|---|---|---|---|---|---|---|---|---|
| ical Monocyte | | | E cCRE-pELS | | | | 63 | | | | 31 |
| nonclassical Monocyte | Mono | CAMBUS_control | ENCODE cCRE-pELS | 0.39 | 0.35 | 0.44 | 4.26.E-55 | 0.41 | 0.36 | 0.48 | 3.33.E-31 |
| nonclassical Monocyte | Mono | cOCRs | ENCODE cCRE-dELS | 2.19 | 1.98 | 2.43 | 3.49.E-50 | 2.47 | 2.19 | 2.79 | 5.99.E-48 |
| nonclassical Monocyte | Mono | CAMBUS_control | ENCODE cCRE-dELS | 0.48 | 0.43 | 0.53 | 5.38.E-48 | 0.43 | 0.38 | 0.48 | 1.18.E-45 |
| nonclassical Monocyte | Mono | cOCRs | ENCODE CTCF-only | 0.39 | 0.22 | 0.67 | 6.98.E-04 | 0.42 | 0.24 | 0.75 | 3.06.E-03 |
| nonclassical Monocyte | Mono | CAMBUS_control | ENCODE CTCF-only | 2.53 | 1.49 | 4.28 | 5.76.E-04 | 2.32 | 1.34 | 4.02 | 2.58.E-03 |
| nonclassical Monocyte | Mono | cOCRs | FANTOM enhancers | 4.12 | 3.69 | 4.60 | 1.01.E-138 | 4.44 | 3.93 | 5.02 | 3.22.E-125 |
| nonclassical Monocyt | Mono | CAMBUS_control | FANTOM enhance | 0.26 | 0.23 | 0.29 | 3.99.E-131 | 0.24 | 0.21 | 0.27 | 1.57.E-118 |

| | | | | | | | | | | | |
|---|---|---|---|---|---|---|---|---|---|---|---|
| e | | | rs | | | | | | | | |
| nonclassical Monocyte | Mono | cOCRs | Super enhancers | 2.40 | 1.98 | 2.91 | 8.33.E-19 | 2.35 | 1.90 | 2.92 | 7.01.E-15 |
| nonclassical Monocyte | Mono | CAMBUS_control | Super enhancers | 0.44 | 0.37 | 0.53 | 2.99.E-19 | 0.45 | 0.37 | 0.55 | 2.09.E-15 |
| intermediate Monocyte | Mono | cOCRs | ENCODE cCRE-PLS | 4.00 | 3.52 | 4.55 | 1.16.E-99 | 4.10 | 3.40 | 4.95 | 2.38.E-49 |
| intermediate Monocyte | Mono | CAMBUS_control | ENCODE cCRE-PLS | 0.30 | 0.27 | 0.34 | 1.04.E-78 | 0.32 | 0.26 | 0.38 | 3.48.E-35 |
| intermediate Monocyte | Mono | cOCRs | ENCODE cCRE-pELS | 4.03 | 3.64 | 4.48 | 1.72.E-151 | 3.88 | 3.39 | 4.43 | 2.61.E-88 |
| intermediate Monocyte | Mono | CAMBUS_control | ENCODE cCRE-pELS | 0.28 | 0.25 | 0.31 | 1.86.E-135 | 0.29 | 0.25 | 0.33 | 1.28.E-77 |
| intermediate Monocyte | Mono | cOCRs | ENCODE cCRE-dELS | 2.78 | 2.54 | 3.05 | 1.43.E-105 | 3.47 | 3.10 | 3.89 | 3.12.E-102 |
| intermed | Mono | CAMBU | ENCOD | 0.37 | 0.34 | 0.40 | 3.23.E- | 0.31 | 0.27 | 0.34 | 6.94.E- |

| | | | | | | | | | | | |
|---|---|---|---|---|---|---|---|---|---|---|---|
| iate Monocyte | | S_control | E cCRE-dELS | | | | 104 | | | | 101 |
| intermediate Monocyte | Mono | cOCRs | ENCODE CTCF-only | 0.47 | 0.27 | 0.80 | 6.18.E-03 | 0.51 | 0.28 | 0.92 | 2.60.E-02 |
| intermediate Monocyte | Mono | CAMBUS_control | ENCODE CTCF-only | 1.84 | 1.12 | 3.02 | 1.53.E-02 | 1.66 | 0.98 | 2.82 | 5.92.E-02 |
| intermediate Monocyte | Mono | cOCRs | FANTOM enhancers | 5.09 | 4.59 | 5.64 | 2.52.E-210 | 5.86 | 5.22 | 6.59 | 7.45.E-194 |
| intermediate Monocyte | Mono | CAMBUS_control | FANTOM enhancers | 0.20 | 0.18 | 0.22 | 1.44.E-207 | 0.18 | 0.16 | 0.20 | 4.01.E-191 |
| intermediate Monocyte | Mono | cOCRs | Super enhancers | 5.16 | 4.18 | 6.37 | 9.41.E-53 | 5.46 | 4.27 | 6.98 | 1.58.E-41 |
| intermediate Monocyte | Mono | CAMBUS_control | Super enhancers | 0.29 | 0.25 | 0.34 | 2.45.E-50 | 0.30 | 0.25 | 0.36 | 3.73.E-40 |
| NK | NK | cOCRs | ENCODE cCRE-PLS | 3.54 | 3.08 | 4.08 | 3.19.E-69 | 3.92 | 3.20 | 4.79 | 3.05.E-40 |

| | | | | | | | | | | | |
|---|---|---|---|---|---|---|---|---|---|---|---|
| NK | NK | CAMBUS_control | ENCODE cCRE-PLS | 0.29 | 0.25 | 0.33 | 4.08.E-67 | 0.27 | 0.22 | 0.33 | 8.83.E-38 |
| NK | NK | cOCRs | ENCODE cCRE-pELS | 3.35 | 2.97 | 3.78 | 3.31.E-84 | 2.99 | 2.54 | 3.51 | 1.47.E-40 |
| NK | NK | CAMBUS_control | ENCODE cCRE-pELS | 0.31 | 0.27 | 0.35 | 3.63.E-81 | 0.34 | 0.29 | 0.40 | 6.49.E-39 |
| NK | NK | cOCRs | ENCODE cCRE-dELS | 2.63 | 2.35 | 2.94 | 1.57.E-64 | 3.47 | 3.00 | 4.00 | 1.10.E-64 |
| NK | NK | CAMBUS_control | ENCODE cCRE-dELS | 0.38 | 0.34 | 0.43 | 2.28.E-64 | 0.29 | 0.25 | 0.34 | 7.67.E-65 |
| NK | NK | cOCRs | ENCODE CTCF-only | 0.43 | 0.24 | 0.78 | 5.14.E-03 | 0.44 | 0.23 | 0.86 | 1.54.E-02 |
| NK | NK | CAMBUS_control | ENCODE CTCF-only | 2.37 | 1.31 | 4.30 | 4.47.E-03 | 2.30 | 1.19 | 4.43 | 1.34.E-02 |
| NK | NK | cOCRs | FANTOM enhancers | 4.71 | 4.18 | 5.31 | 4.27.E-142 | 6.00 | 5.23 | 6.88 | 1.15.E-143 |
| NK | NK | CAMBUS_control | FANTOM enhancers | 0.21 | 0.19 | 0.24 | 3.78.E-142 | 0.17 | 0.15 | 0.19 | 3.48.E-144 |
| NK | NK | cOCRs | Super | 4.11 | 3.17 | 5.33 | 2.03.E- | 4.89 | 3.53 | 6.76 | 1.20.E- |

| | | | | | | | | | | | |
|---|---|---|---|---|---|---|---|---|---|---|---|
| | | | enhancers | | | | 26 | | | | 21 |
| NK | NK | CAMBUS_control | Super enhancers | 0.26 | 0.20 | 0.33 | 9.99.E-27 | 0.23 | 0.17 | 0.30 | 3.85.E-22 |
| PBMC | PBMC | cOCRs | ENCODE cCRE-PLS | 3.62 | 3.34 | 3.92 | 1.56.E-219 | 3.48 | 3.10 | 3.91 | 4.46.E-98 |
| PBMC | PBMC | CAMBUS_control | ENCODE cCRE-PLS | 0.28 | 0.26 | 0.31 | 7.01.E-211 | 0.30 | 0.27 | 0.34 | 9.85.E-93 |
| PBMC | PBMC | cOCRs | ENCODE cCRE-pELS | 3.73 | 3.49 | 3.99 | <2e-16 | 3.50 | 3.21 | 3.81 | 2.72.E-180 |
| PBMC | PBMC | CAMBUS_control | ENCODE cCRE-pELS | 0.27 | 0.25 | 0.29 | <2e-16 | 0.29 | 0.27 | 0.32 | 2.06.E-177 |
| PBMC | PBMC | cOCRs | ENCODE cCRE-dELS | 2.67 | 2.52 | 2.83 | 8.76.E-241 | 3.37 | 3.14 | 3.61 | 1.88.E-247 |
| PBMC | PBMC | CAMBUS_control | ENCODE cCRE-dELS | 0.38 | 0.36 | 0.40 | 5.87.E-241 | 0.30 | 0.28 | 0.32 | 7.16.E-248 |
| PBMC | PBMC | cOCRs | ENCODE CTCF-only | 0.67 | 0.50 | 0.90 | 8.77.E-03 | 0.67 | 0.47 | 0.94 | 1.93.E-02 |
| PBMC | PBMC | CAMBUS_control | ENCODE CTCF-only | 1.47 | 1.09 | 1.98 | 1.05.E-02 | 1.48 | 1.05 | 2.07 | 2.33.E-02 |
| PBMC | PBMC | cOCRs | FANTO | 4.47 | 4.17 | 4.80 | <2e-16 | 5.22 | 4.83 | 5.65 | <2e-16 |

| | | | | | | | | | | | |
|---|---|---|---|---|---|---|---|---|---|---|---|
| | | | M enhancers | | | | | | | | |
| PBMC | PBMC | CAMBUS_control | FANTOM enhancers | 0.22 | 0.21 | 0.24 | <2e-16 | 0.19 | 0.18 | 0.21 | <2e-16 |
| PBMC | PBMC | cOCRs | Super enhancers | 5.11 | 4.47 | 5.84 | 3.91.E-126 | 5.79 | 4.93 | 6.82 | 1.57.E-99 |
| PBMC | PBMC | CAMBUS_control | Super enhancers | 0.22 | 0.19 | 0.25 | 1.52.E-124 | 0.20 | 0.17 | 0.23 | 5.68.E-99 |
| Neutrophil | Neutrophil | cOCRs | ENCODE cCRE-PLS | 2.40 | 1.79 | 3.23 | 6.40.E-09 | 2.23 | 1.41 | 3.52 | 5.73.E-04 |
| Neutrophil | Neutrophil | CAMBUS_control | ENCODE cCRE-PLS | 0.42 | 0.32 | 0.57 | 1.20.E-08 | 0.45 | 0.28 | 0.71 | 5.77.E-04 |
| Neutrophil | Neutrophil | cOCRs | ENCODE cCRE-pELS | 3.39 | 2.66 | 4.32 | 5.11.E-23 | 3.21 | 2.30 | 4.47 | 7.70.E-12 |
| Neutrophil | Neutrophil | CAMBUS_control | ENCODE cCRE-pELS | 0.30 | 0.23 | 0.38 | 9.30.E-23 | 0.31 | 0.22 | 0.44 | 8.02.E-12 |
| Neutrophil | Neutrophil | cOCRs | ENCODE cCRE-dELS | 1.74 | 1.40 | 2.15 | 4.61.E-07 | 1.98 | 1.49 | 2.63 | 2.26.E-06 |
| Neutrophil | Neutrophil | CAMBUS_contro | ENCODE cCRE- | 0.58 | 0.47 | 0.72 | 7.21.E-07 | 0.51 | 0.38 | 0.67 | 2.64.E-06 |

| | | | | | | | | | | | |
|---|---|---|---|---|---|---|---|---|---|---|---|
| | | l | dELS | | | | | | | | |
| Neutrophil | Neutrophil | cOCRs | ENCODE CTCF-only | 0.00 | 0.00 | 2.E+60 | 8.88.E-01 | 0.00 | 0.00 | 1.E+47 | 8.70.E-01 |
| Neutrophil | Neutrophil | CAMBUS_control | ENCODE CTCF-only | 48131.64 | 0.00 | 1.E+69 | 8.87.E-01 | 20086.80 | 0.00 | 3.E+55 | 8.69.E-01 |
| Neutrophil | Neutrophil | cOCRs | FANTOM enhancers | 3.94 | 2.99 | 5.20 | 2.81.E-22 | 4.92 | 3.55 | 6.82 | 1.23.E-21 |
| Neutrophil | Neutrophil | CAMBUS_control | FANTOM enhancers | 0.26 | 0.19 | 0.34 | 3.98.E-22 | 0.20 | 0.15 | 0.28 | 1.33.E-21 |
| Neutrophil | Neutrophil | cOCRs | Super enhancers | 10.59 | 4.37 | 25.64 | 1.70.E-07 | 11.62 | 3.71 | 36.40 | 2.55.E-05 |
| Neutrophil | Neutrophil | CAMBUS_control | Super enhancers | 0.13 | 0.06 | 0.27 | 6.72.E-08 | 0.11 | 0.04 | 0.29 | 1.25.E-05 |

Supplementary Table 10 | Enrichment of cOCRs OCRs in the tissue expressed enhancers.
OR, odds ratio; CI_lo, lower 95% confidence interval;
CI_hi, upper 95% confidence interval.

| name | subgroup | constraint | enhancer_type | tissue | OR | ci_lo | ci_hi | P |
|---|---|---|---|---|---|---|---|---|
| Naive CD4 | CD4 | cOCRs | Super enhancers | Blood | 6.84 | 6.07 | 7.71 | 1.10.E-217 |
| Naive CD4 | CD4 | CAMBUS control | Super enhancers | Blood | 0.43 | 0.40 | 0.46 | 5.88.E-111 |
| Naive CD4 | CD4 | cOCRs | Super enhancers | Brain | 4.56 | 4.06 | 5.12 | 1.16.E-146 |
| Naive CD4 | CD4 | CAMBUS control | Super enhancers | Brain | 0.65 | 0.60 | 0.70 | 1.70.E-28 |
| Naive CD4 | CD4 | cOCRs | Super enhancers | Kidney | 3.65 | 3.24 | 4.11 | 3.45.E-102 |
| Naive CD4 | CD4 | CAMBUS control | Super enhancers | Kidney | 0.53 | 0.49 | 0.59 | 1.39.E-40 |
| Naive CD4 | CD4 | cOCRs | Super enhancers | Lung | 3.68 | 3.29 | 4.12 | 1.49.E-115 |
| Naive CD4 | CD4 | CAMBUS control | Super enhancers | Lung | 0.64 | 0.59 | 0.69 | 1.18.E-30 |
| Naive CD4 | CD4 | cOCRs | FANTOM enhancers | Hemato | 7.64 | 6.64 | 8.78 | 8.74.E-179 |
| Naive CD4 | CD4 | CAMBUS control | FANTOM enhancers | Hemato | 0.32 | 0.28 | 0.36 | 4.20.E-69 |
| Naive CD4 | CD4 | cOCRs | FANTOM enhancers | Brain | 6.54 | 5.62 | 7.62 | 3.71.E-128 |
| Naive CD4 | CD4 | CAMBUS control | FANTOM enhancers | Brain | 0.36 | 0.31 | 0.41 | 2.32.E-47 |
| Naive CD4 | CD4 | cOCRs | FANTOM enhancers | Kidney | 7.30 | 6.07 | 8.79 | 1.12.E-98 |
| Naive CD4 | CD4 | CAMBUS control | FANTOM enhancers | Kidney | 0.32 | 0.27 | 0.38 | 3.29.E-37 |
| Naive CD4 | CD4 | cOCRs | FANTOM enhancers | Lung | 7.01 | 6.01 | 8.18 | 3.82.E-135 |
| Naive CD4 | CD4 | CAMBUS control | FANTOM enhancers | Lung | 0.34 | 0.29 | 0.39 | 2.85.E-50 |
| Th1 | CD4 | cOCRs | Super enhancers | Blood | 4.67 | 4.29 | 5.07 | 1.82.E-287 |
| Th1 | CD4 | CAMBUS control | Super enhancers | Blood | 0.37 | 0.34 | 0.39 | 1.03.E-205 |
| Th1 | CD4 | cOCRs | Super enhancers | Brain | 2.98 | 2.75 | 3.24 | 1.51.E-155 |
| Th1 | CD4 | CAMBUS control | Super enhancers | Brain | 0.57 | 0.53 | 0.60 | 2.73.E-66 |
| Th1 | CD4 | cOCRs | Super enhancers | Kidney | 2.75 | 2.52 | 3.01 | 7.40.E-111 |
| Th1 | CD4 | CAMBUS control | Super enhancers | Kidney | 0.50 | 0.46 | 0.54 | 1.40.E-68 |

| | | | | | | | | |
|---|---|---|---|---|---|---|---|---|
| Th1 | CD4 | cOCRs | Super enhancers | Lung | 2.99 | 2.76 | 3.24 | 2.03.E-157 |
| Th1 | CD4 | CAMBUS control | Super enhancers | Lung | 0.52 | 0.49 | 0.56 | 3.72.E-85 |
| Th1 | CD4 | cOCRs | FANTOM enhancers | Hemato | 6.77 | 6.14 | 7.46 | 0.00.E+00 |
| Th1 | CD4 | CAMBUS control | FANTOM enhancers | Hemato | 0.24 | 0.22 | 0.27 | 3.14.E-203 |
| Th1 | CD4 | cOCRs | FANTOM enhancers | Brain | 5.59 | 5.01 | 6.23 | 6.38.E-211 |
| Th1 | CD4 | CAMBUS control | FANTOM enhancers | Brain | 0.28 | 0.26 | 0.32 | 5.44.E-126 |
| Th1 | CD4 | cOCRs | FANTOM enhancers | Kidney | 5.95 | 5.21 | 6.80 | 4.05.E-150 |
| Th1 | CD4 | CAMBUS control | FANTOM enhancers | Kidney | 0.26 | 0.23 | 0.30 | 4.42.E-91 |
| Th1 | CD4 | cOCRs | FANTOM enhancers | Lung | 6.17 | 5.54 | 6.88 | 1.71.E-238 |
| Th1 | CD4 | CAMBUS control | FANTOM enhancers | Lung | 0.26 | 0.23 | 0.29 | 9.36.E-147 |
| Th2 | CD4 | cOCRs | Super enhancers | Blood | 5.50 | 5.05 | 6.00 | 0.00.E+00 |
| Th2 | CD4 | CAMBUS control | Super enhancers | Blood | 0.44 | 0.41 | 0.47 | 2.45.E-168 |
| Th2 | CD4 | cOCRs | Super enhancers | Brain | 3.41 | 3.14 | 3.71 | 6.06.E-184 |
| Th2 | CD4 | CAMBUS control | Super enhancers | Brain | 0.68 | 0.64 | 0.72 | 4.23.E-38 |
| Th2 | CD4 | cOCRs | Super enhancers | Kidney | 3.05 | 2.78 | 3.34 | 1.26.E-127 |
| Th2 | CD4 | CAMBUS control | Super enhancers | Kidney | 0.57 | 0.53 | 0.61 | 4.36.E-52 |
| Th2 | CD4 | cOCRs | Super enhancers | Lung | 3.33 | 3.07 | 3.62 | 3.46.E-179 |
| Th2 | CD4 | CAMBUS control | Super enhancers | Lung | 0.61 | 0.57 | 0.64 | 1.33.E-61 |
| Th2 | CD4 | cOCRs | FANTOM enhancers | Hemato | 7.69 | 6.95 | 8.51 | 0.00.E+00 |
| Th2 | CD4 | CAMBUS control | FANTOM enhancers | Hemato | 0.29 | 0.26 | 0.32 | 2.40.E-151 |
| Th2 | CD4 | cOCRs | FANTOM enhancers | Brain | 6.24 | 5.57 | 6.99 | 1.94.E-220 |
| Th2 | CD4 | CAMBUS control | FANTOM enhancers | Brain | 0.34 | 0.30 | 0.38 | 3.04.E-90 |
| Th2 | CD4 | cOCRs | FANTOM enhancers | Kidney | 6.75 | 5.87 | 7.76 | 4.95.E-158 |
| Th2 | CD4 | CAMBUS control | FANTOM enhancers | Kidney | 0.31 | 0.27 | 0.35 | 8.16.E-67 |
| Th2 | CD4 | cOCRs | FANTOM enhancers | Lung | 6.80 | 6.07 | 7.62 | 1.11.E-239 |
| Th2 | CD4 | CAMBUS control | FANTOM enhancers | Lung | 0.31 | 0.28 | 0.35 | 4.62.E-102 |

| | | | | | | | | |
|---|---|---|---|---|---|---|---|---|
| Th17 | CD4 | cOCRs | Super enhancers | Blood | 4.18 | 3.91 | 4.46 | 0.00.E+00 |
| Th17 | CD4 | CAMBUS control | Super enhancers | Blood | 0.34 | 0.32 | 0.36 | 0.00.E+00 |
| Th17 | CD4 | cOCRs | Super enhancers | Brain | 2.82 | 2.64 | 3.00 | 1.42.E-220 |
| Th17 | CD4 | CAMBUS control | Super enhancers | Brain | 0.51 | 0.48 | 0.54 | 4.88.E-123 |
| Th17 | CD4 | cOCRs | Super enhancers | Kidney | 2.58 | 2.40 | 2.77 | 1.04.E-144 |
| Th17 | CD4 | CAMBUS control | Super enhancers | Kidney | 0.47 | 0.44 | 0.51 | 8.48.E-108 |
| Th17 | CD4 | cOCRs | Super enhancers | Lung | 2.84 | 2.66 | 3.03 | 5.77.E-225 |
| Th17 | CD4 | CAMBUS control | Super enhancers | Lung | 0.47 | 0.45 | 0.50 | 1.63.E-152 |
| Th17 | CD4 | cOCRs | FANTOM enhancers | Hemat o | 6.52 | 6.02 | 7.06 | 0.00.E+00 |
| Th17 | CD4 | CAMBUS control | FANTOM enhancers | Hemat o | 0.21 | 0.20 | 0.23 | 0.00.E+00 |
| Th17 | CD4 | cOCRs | FANTOM enhancers | Brain | 5.28 | 4.83 | 5.77 | 1.83.E-290 |
| Th17 | CD4 | CAMBUS control | FANTOM enhancers | Brain | 0.26 | 0.24 | 0.28 | 1.38.E-205 |
| Th17 | CD4 | cOCRs | FANTOM enhancers | Kidney | 5.63 | 5.04 | 6.30 | 1.19.E-200 |
| Th17 | CD4 | CAMBUS control | FANTOM enhancers | Kidney | 0.24 | 0.21 | 0.27 | 2.84.E-145 |
| Th17 | CD4 | cOCRs | FANTOM enhancers | Lung | 5.85 | 5.35 | 6.40 | 0.00.E+00 |
| Th17 | CD4 | CAMBUS control | FANTOM enhancers | Lung | 0.23 | 0.21 | 0.25 | 1.53.E-238 |
| Tfh | CD4 | cOCRs | Super enhancers | Blood | 5.94 | 5.34 | 6.59 | 4.12.E-242 |
| Tfh | CD4 | CAMBUS control | Super enhancers | Blood | 0.39 | 0.36 | 0.42 | 1.23.E-140 |
| Tfh | CD4 | cOCRs | Super enhancers | Brain | 3.86 | 3.49 | 4.27 | 3.17.E-150 |
| Tfh | CD4 | CAMBUS control | Super enhancers | Brain | 0.60 | 0.55 | 0.64 | 3.34.E-42 |
| Tfh | CD4 | cOCRs | Super enhancers | Kidney | 3.32 | 2.98 | 3.69 | 2.22.E-106 |
| Tfh | CD4 | CAMBUS control | Super enhancers | Kidney | 0.49 | 0.45 | 0.54 | 1.02.E-55 |
| Tfh | CD4 | cOCRs | Super enhancers | Lung | 3.58 | 3.24 | 3.96 | 5.74.E-138 |
| Tfh | CD4 | CAMBUS control | Super enhancers | Lung | 0.55 | 0.51 | 0.59 | 3.49.E-57 |
| Tfh | CD4 | cOCRs | FANTOM enhancers | Hemat o | 7.86 | 6.97 | 8.87 | 1.10.E-246 |
| Tfh | CD4 | CAMBUS control | FANTOM enhancers | Hemat o | 0.26 | 0.23 | 0.29 | 6.96.E-123 |

| | | | | | | | | |
|---|---|---|---|---|---|---|---|---|
| Tfh | CD4 | cOCRs | FANTOM enhancers | Brain | 6.66 | 5.83 | 7.60 | 6.84.E-173 |
| Tfh | CD4 | CAMBUS control | FANTOM enhancers | Brain | 0.29 | 0.26 | 0.33 | 3.27.E-82 |
| Tfh | CD4 | cOCRs | FANTOM enhancers | Kidney | 7.64 | 6.52 | 8.95 | 1.15.E-138 |
| Tfh | CD4 | CAMBUS control | FANTOM enhancers | Kidney | 0.25 | 0.22 | 0.29 | 1.13.E-69 |
| Tfh | CD4 | cOCRs | FANTOM enhancers | Lung | 7.09 | 6.20 | 8.10 | 8.92.E-181 |
| Tfh | CD4 | CAMBUS control | FANTOM enhancers | Lung | 0.28 | 0.24 | 0.32 | 1.61.E-87 |
| FraI | CD4 | cOCRs | Super enhancers | Blood | 6.40 | 5.69 | 7.20 | 5.47.E-211 |
| FraI | CD4 | CAMBUS control | Super enhancers | Blood | 0.36 | 0.33 | 0.39 | 2.15.E-130 |
| FraI | CD4 | cOCRs | Super enhancers | Brain | 4.10 | 3.66 | 4.59 | 1.17.E-133 |
| FraI | CD4 | CAMBUS control | Super enhancers | Brain | 0.54 | 0.50 | 0.59 | 3.48.E-47 |
| FraI | CD4 | cOCRs | Super enhancers | Kidney | 3.45 | 3.07 | 3.87 | 1.27.E-95 |
| FraI | CD4 | CAMBUS control | Super enhancers | Kidney | 0.48 | 0.44 | 0.54 | 6.91.E-46 |
| FraI | CD4 | cOCRs | Super enhancers | Lung | 3.68 | 3.30 | 4.11 | 8.74.E-119 |
| FraI | CD4 | CAMBUS control | Super enhancers | Lung | 0.51 | 0.47 | 0.56 | 4.42.E-55 |
| FraI | CD4 | cOCRs | FANTOM enhancers | Hemat o | 7.33 | 6.41 | 8.39 | 4.79.E-184 |
| FraI | CD4 | CAMBUS control | FANTOM enhancers | Hemat o | 0.26 | 0.23 | 0.29 | 8.12.E-99 |
| FraI | CD4 | cOCRs | FANTOM enhancers | Brain | 6.45 | 5.56 | 7.47 | 3.25.E-136 |
| FraI | CD4 | CAMBUS control | FANTOM enhancers | Brain | 0.28 | 0.25 | 0.33 | 1.62.E-70 |
| FraI | CD4 | cOCRs | FANTOM enhancers | Kidney | 7.33 | 6.15 | 8.74 | 4.66.E-109 |
| FraI | CD4 | CAMBUS control | FANTOM enhancers | Kidney | 0.25 | 0.21 | 0.29 | 1.18.E-58 |
| FraI | CD4 | cOCRs | FANTOM enhancers | Lung | 6.92 | 5.97 | 8.02 | 2.73.E-145 |
| FraI | CD4 | CAMBUS control | FANTOM enhancers | Lung | 0.26 | 0.23 | 0.30 | 7.63.E-78 |
| FraII (aTreg) | CD4 | cOCRs | Super enhancers | Blood | 5.79 | 5.20 | 6.45 | 9.31.E-226 |
| FraII (aTreg) | CD4 | CAMBUS control | Super enhancers | Blood | 0.61 | 0.58 | 0.65 | 1.47.E-59 |
| FraII (aTreg) | CD4 | cOCRs | Super enhancers | Brain | 4.00 | 3.60 | 4.44 | 4.18.E-150 |
| FraII (aTreg) | CD4 | CAMBUS control | Super enhancers | Brain | 0.89 | 0.84 | 0.94 | 1.28.E-04 |
| FraII (aTreg) | CD4 | cOCRs | Super enhancers | Kidney | 3.41 | 3.06 | 3.80 | 1.31.E-109 |
| FraII (aTreg) | CD4 | CAMBUS control | Super enhancers | Kidney | 0.69 | 0.64 | 0.74 | 5.77.E-22 |

| | | | | | | | | |
|---|---|---|---|---|---|---|---|---|
| FraII (aTreg) | CD4 | cOCRs | Super enhancers | Lung | 3.57 | 3.22 | 3.95 | 9.93.E-132 |
| FraII (aTreg) | CD4 | CAMBUS control | Super enhancers | Lung | 0.78 | 0.74 | 0.83 | 2.46.E-15 |
| FraII (aTreg) | CD4 | cOCRs | FANTOM enhancers | Hemato | 7.51 | 6.64 | 8.49 | 1.90.E-227 |
| FraII (aTreg) | CD4 | CAMBUS control | FANTOM enhancers | Hemato | 0.45 | 0.40 | 0.50 | 1.79.E-48 |
| FraII (aTreg) | CD4 | cOCRs | FANTOM enhancers | Brain | 6.62 | 5.79 | 7.56 | 2.08.E-168 |
| FraII (aTreg) | CD4 | CAMBUS control | FANTOM enhancers | Brain | 0.47 | 0.42 | 0.53 | 9.75.E-36 |
| FraII (aTreg) | CD4 | cOCRs | FANTOM enhancers | Kidney | 7.11 | 6.04 | 8.37 | 1.20.E-122 |
| FraII (aTreg) | CD4 | CAMBUS control | FANTOM enhancers | Kidney | 0.44 | 0.38 | 0.51 | 7.71.E-27 |
| FraII (aTreg) | CD4 | cOCRs | FANTOM enhancers | Lung | 6.91 | 6.03 | 7.91 | 9.24.E-172 |
| FraII (aTreg) | CD4 | CAMBUS control | FANTOM enhancers | Lung | 0.47 | 0.42 | 0.53 | 2.34.E-34 |
| FraIII | CD4 | cOCRs | Super enhancers | Blood | 3.95 | 3.69 | 4.24 | 0.00.E+00 |
| FraIII | CD4 | CAMBUS control | Super enhancers | Blood | 0.37 | 0.35 | 0.39 | 2.13.E-252 |
| FraIII | CD4 | cOCRs | Super enhancers | Brain | 2.67 | 2.50 | 2.86 | 1.45.E-176 |
| FraIII | CD4 | CAMBUS control | Super enhancers | Brain | 0.56 | 0.52 | 0.59 | 4.84.E-87 |
| FraIII | CD4 | cOCRs | Super enhancers | Kidney | 2.44 | 2.26 | 2.63 | 5.76.E-115 |
| FraIII | CD4 | CAMBUS control | Super enhancers | Kidney | 0.52 | 0.48 | 0.56 | 1.56.E-76 |
| FraIII | CD4 | cOCRs | Super enhancers | Lung | 2.69 | 2.51 | 2.88 | 6.23.E-179 |
| FraIII | CD4 | CAMBUS control | Super enhancers | Lung | 0.51 | 0.48 | 0.54 | 5.86.E-114 |
| FraIII | CD4 | cOCRs | FANTOM enhancers | Hemato | 6.10 | 5.62 | 6.63 | 0.00.E+00 |
| FraIII | CD4 | CAMBUS control | FANTOM enhancers | Hemato | 0.23 | 0.22 | 0.25 | 3.50.E-286 |
| FraIII | CD4 | cOCRs | FANTOM enhancers | Brain | 4.94 | 4.50 | 5.42 | 3.84.E-247 |
| FraIII | CD4 | CAMBUS control | FANTOM enhancers | Brain | 0.28 | 0.26 | 0.31 | 6.39.E-172 |
| FraIII | CD4 | cOCRs | FANTOM enhancers | Kidney | 5.28 | 4.70 | 5.93 | 1.57.E-174 |
| FraIII | CD4 | CAMBUS control | FANTOM enhancers | Kidney | 0.26 | 0.23 | 0.29 | 7.89.E-123 |
| FraIII | CD4 | cOCRs | FANTOM enhancers | Lung | 5.41 | 4.93 | 5.94 | 3.09.E-275 |
| FraIII | CD4 | CAMBUS control | FANTOM enhancers | Lung | 0.26 | 0.23 | 0.28 | 8.27.E-195 |

| | | | | | | | | |
|---|---|---|---|---|---|---|---|---|
| LAG3 Treg | CD4 | cOCRs | Super enhancers | Blood | 3.75 | 3.52 | 4.00 | 0.00.E+00 |
| LAG3 Treg | CD4 | CAMBUS control | Super enhancers | Blood | 0.38 | 0.36 | 0.40 | 1.22.E-279 |
| LAG3 Treg | CD4 | cOCRs | Super enhancers | Brain | 2.47 | 2.32 | 2.63 | 2.02.E-179 |
| LAG3 Treg | CD4 | CAMBUS control | Super enhancers | Brain | 0.57 | 0.54 | 0.60 | 1.71.E-94 |
| LAG3 Treg | CD4 | cOCRs | Super enhancers | Kidney | 2.31 | 2.15 | 2.48 | 1.02.E-118 |
| LAG3 Treg | CD4 | CAMBUS control | Super enhancers | Kidney | 0.53 | 0.50 | 0.57 | 4.30.E-86 |
| LAG3 Treg | CD4 | cOCRs | Super enhancers | Lung | 2.61 | 2.46 | 2.78 | 8.28.E-202 |
| LAG3 Treg | CD4 | CAMBUS control | Super enhancers | Lung | 0.52 | 0.49 | 0.55 | 3.94.E-131 |
| LAG3 Treg | CD4 | cOCRs | FANTOM enhancers | Hemato | 5.53 | 5.11 | 5.97 | 0.00.E+00 |
| LAG3 Treg | CD4 | CAMBUS control | FANTOM enhancers | Hemato | 0.26 | 0.24 | 0.28 | 3.85.E-280 |
| LAG3 Treg | CD4 | cOCRs | FANTOM enhancers | Brain | 4.52 | 4.14 | 4.93 | 1.80.E-248 |
| LAG3 Treg | CD4 | CAMBUS control | FANTOM enhancers | Brain | 0.31 | 0.28 | 0.33 | 2.38.E-168 |
| LAG3 Treg | CD4 | cOCRs | FANTOM enhancers | Kidney | 4.80 | 4.30 | 5.36 | 1.05.E-171 |
| LAG3 Treg | CD4 | CAMBUS control | FANTOM enhancers | Kidney | 0.28 | 0.26 | 0.32 | 1.06.E-119 |
| LAG3 Treg | CD4 | cOCRs | FANTOM enhancers | Lung | 4.95 | 4.53 | 5.40 | 3.61.E-278 |
| LAG3 Treg | CD4 | CAMBUS control | FANTOM enhancers | Lung | 0.28 | 0.26 | 0.31 | 1.32.E-189 |
| Memory CD4 | CD4 | cOCRs | Super enhancers | Blood | 5.10 | 4.70 | 5.53 | 0.00.E+00 |
| Memory CD4 | CD4 | CAMBUS control | Super enhancers | Blood | 0.34 | 0.32 | 0.36 | 2.63.E-233 |
| Memory CD4 | CD4 | cOCRs | Super enhancers | Brain | 3.33 | 3.08 | 3.61 | 1.03.E-192 |
| Memory CD4 | CD4 | CAMBUS control | Super enhancers | Brain | 0.51 | 0.48 | 0.54 | 6.28.E-93 |
| Memory CD4 | CD4 | cOCRs | Super enhancers | Kidney | 3.05 | 2.79 | 3.32 | 7.70.E-139 |
| Memory CD4 | CD4 | CAMBUS control | Super enhancers | Kidney | 0.45 | 0.42 | 0.49 | 4.21.E-90 |
| Memory CD4 | CD4 | cOCRs | Super enhancers | Lung | 3.22 | 2.98 | 3.49 | 1.56.E-184 |
| Memory CD4 | CD4 | CAMBUS control | Super enhancers | Lung | 0.49 | 0.46 | 0.53 | 2.36.E-100 |
| Memory CD4 | CD4 | cOCRs | FANTOM enhancers | Hemato | 7.22 | 6.55 | 7.97 | 0.00.E+00 |
| Memory CD4 | CD4 | CAMBUS control | FANTOM enhancers | Hemato | 0.22 | 0.20 | 0.24 | 2.39.E-220 |

| | | | | | | | | |
|---|---|---|---|---|---|---|---|---|
| Memory CD4 | CD4 | cOCRs | FANTOM enhancers | Brain | 6.07 | 5.45 | 6.77 | 3.39.E-232 |
| Memory CD4 | CD4 | CAMBUS control | FANTOM enhancers | Brain | 0.26 | 0.23 | 0.29 | 4.58.E-142 |
| Memory CD4 | CD4 | cOCRs | FANTOM enhancers | Kidney | 6.52 | 5.70 | 7.46 | 2.94.E-163 |
| Memory CD4 | CD4 | CAMBUS control | FANTOM enhancers | Kidney | 0.24 | 0.21 | 0.27 | 7.89.E-103 |
| Memory CD4 | CD4 | cOCRs | FANTOM enhancers | Lung | 6.58 | 5.90 | 7.34 | 3.57.E-251 |
| Memory CD4 | CD4 | CAMBUS control | FANTOM enhancers | Lung | 0.24 | 0.21 | 0.26 | 7.82.E-160 |
| Thx | CD4 | cOCRs | Super enhancers | Blood | 4.01 | 3.74 | 4.29 | 0.00.E+00 |
| Thx | CD4 | CAMBUS control | Super enhancers | Blood | 0.33 | 0.31 | 0.35 | 1.80.E-281 |
| Thx | CD4 | cOCRs | Super enhancers | Brain | 2.64 | 2.46 | 2.82 | 9.29.E-173 |
| Thx | CD4 | CAMBUS control | Super enhancers | Brain | 0.49 | 0.46 | 0.52 | 4.06.E-114 |
| Thx | CD4 | cOCRs | Super enhancers | Kidney | 2.50 | 2.32 | 2.70 | 3.40.E-122 |
| Thx | CD4 | CAMBUS control | Super enhancers | Kidney | 0.47 | 0.44 | 0.50 | 6.60.E-96 |
| Thx | CD4 | cOCRs | Super enhancers | Lung | 2.80 | 2.61 | 2.99 | 6.32.E-194 |
| Thx | CD4 | CAMBUS control | Super enhancers | Lung | 0.45 | 0.42 | 0.48 | 1.11.E-147 |
| Thx | CD4 | cOCRs | FANTOM enhancers | Hemato | 6.10 | 5.61 | 6.63 | 0.00.E+00 |
| Thx | CD4 | CAMBUS control | FANTOM enhancers | Hemato | 0.21 | 0.19 | 0.23 | 0.00.E+00 |
| Thx | CD4 | cOCRs | FANTOM enhancers | Brain | 5.24 | 4.77 | 5.75 | 2.74.E-265 |
| Thx | CD4 | CAMBUS control | FANTOM enhancers | Brain | 0.24 | 0.22 | 0.26 | 5.42.E-207 |
| Thx | CD4 | cOCRs | FANTOM enhancers | Kidney | 5.53 | 4.93 | 6.22 | 1.51.E-182 |
| Thx | CD4 | CAMBUS control | FANTOM enhancers | Kidney | 0.22 | 0.20 | 0.25 | 2.88.E-144 |
| Thx | CD4 | cOCRs | FANTOM enhancers | Lung | 5.63 | 5.13 | 6.19 | 6.24.E-288 |
| Thx | CD4 | CAMBUS control | FANTOM enhancers | Lung | 0.22 | 0.20 | 0.25 | 1.01.E-226 |
| Naive CD8 | CD8 | cOCRs | Super enhancers | Blood | 17.17 | 12.68 | 23.27 | 3.08.E-75 |
| Naive CD8 | CD8 | CAMBUS control | Super enhancers | Blood | 0.36 | 0.31 | 0.42 | 3.34.E-41 |
| Naive CD8 | CD8 | cOCRs | Super enhancers | Brain | 14.71 | 10.74 | 20.13 | 3.59.E-63 |
| Naive CD8 | CD8 | CAMBUS control | Super enhancers | Brain | 0.50 | 0.43 | 0.58 | 6.22.E-20 |

| | | | | | | | | |
|---|---|---|---|---|---|---|---|---|
| Naive CD8 | CD8 | cOCRs | Super enhancers | Kidney | 7.63 | 6.08 | 9.57 | 2.76.E-69 |
| Naive CD8 | CD8 | CAMBUS control | Super enhancers | Kidney | 0.38 | 0.32 | 0.45 | 3.56.E-28 |
| Naive CD8 | CD8 | cOCRs | Super enhancers | Lung | 5.02 | 3.97 | 6.34 | 1.63.E-41 |
| Naive CD8 | CD8 | CAMBUS control | Super enhancers | Lung | 0.59 | 0.50 | 0.69 | 1.32.E-11 |
| Naive CD8 | CD8 | cOCRs | FANTOM enhancers | Hemato | 6.01 | 4.32 | 8.36 | 1.71.E-26 |
| Naive CD8 | CD8 | CAMBUS control | FANTOM enhancers | Hemato | 0.42 | 0.31 | 0.58 | 1.05.E-07 |
| Naive CD8 | CD8 | cOCRs | FANTOM enhancers | Brain | 4.95 | 3.43 | 7.15 | 1.35.E-17 |
| Naive CD8 | CD8 | CAMBUS control | FANTOM enhancers | Brain | 0.49 | 0.34 | 0.69 | 6.02.E-05 |
| Naive CD8 | CD8 | cOCRs | FANTOM enhancers | Kidney | 6.42 | 4.19 | 9.84 | 1.22.E-17 |
| Naive CD8 | CD8 | CAMBUS control | FANTOM enhancers | Kidney | 0.38 | 0.25 | 0.58 | 5.74.E-06 |
| Naive CD8 | CD8 | cOCRs | FANTOM enhancers | Lung | 6.10 | 4.29 | 8.69 | 1.05.E-23 |
| Naive CD8 | CD8 | CAMBUS control | FANTOM enhancers | Lung | 0.41 | 0.29 | 0.58 | 3.37.E-07 |
| CD8+effector | CD8 | cOCRs | Super enhancers | Blood | 8.73 | 7.47 | 10.22 | 7.91.E-162 |
| CD8+effector | CD8 | CAMBUS control | Super enhancers | Blood | 0.27 | 0.24 | 0.30 | 6.74.E-115 |
| CD8+effector | CD8 | cOCRs | Super enhancers | Brain | 5.33 | 4.60 | 6.17 | 1.49.E-110 |
| CD8+effector | CD8 | CAMBUS control | Super enhancers | Brain | 0.41 | 0.37 | 0.46 | 2.80.E-54 |
| CD8+effector | CD8 | cOCRs | Super enhancers | Kidney | 4.26 | 3.69 | 4.92 | 7.77.E-87 |
| CD8+effector | CD8 | CAMBUS control | Super enhancers | Kidney | 0.36 | 0.32 | 0.41 | 4.20.E-55 |
| CD8+effector | CD8 | cOCRs | Super enhancers | Lung | 4.35 | 3.78 | 5.01 | 9.37.E-93 |
| CD8+effector | CD8 | CAMBUS control | Super enhancers | Lung | 0.43 | 0.38 | 0.48 | 4.93.E-50 |
| CD8+effector | CD8 | cOCRs | FANTOM enhancers | Hemato | 9.01 | 7.65 | 10.61 | 2.34.E-153 |
| CD8+effector | CD8 | CAMBUS control | FANTOM enhancers | Hemato | 0.19 | 0.16 | 0.22 | 4.75.E-101 |
| CD8+effector | CD8 | cOCRs | FANTOM enhancers | Brain | 8.13 | 6.81 | 9.70 | 2.21.E-119 |
| CD8+effector | CD8 | CAMBUS control | FANTOM enhancers | Brain | 0.20 | 0.17 | 0.24 | 3.45.E-75 |
| CD8+effector | CD8 | cOCRs | FANTOM enhancers | Kidney | 9.55 | 7.76 | 11.76 | 2.07.E-100 |
| CD8+effector | CD8 | CAMBUS control | FANTOM enhancers | Kidney | 0.17 | 0.14 | 0.21 | 2.11.E-66 |

| | | | | | | | | |
|---|---|---|---|---|---|---|---|---|
| CD8+effector | CD8 | cOCRs | FANTOM enhancers | Lung | 8.64 | 7.24 | 10.32 | 4.84.E-125 |
| CD8+effector | CD8 | CAMBUS control | FANTOM enhancers | Lung | 0.19 | 0.16 | 0.23 | 3.92.E-80 |
| CD8+central memory | CD8 | cOCRs | Super enhancers | Blood | 7.19 | 6.34 | 8.16 | 8.83.E-206 |
| CD8+central memory | CD8 | CAMBUS control | Super enhancers | Blood | 0.34 | 0.31 | 0.37 | 3.45.E-134 |
| CD8+central memory | CD8 | cOCRs | Super enhancers | Brain | 4.53 | 4.01 | 5.11 | 4.87.E-133 |
| CD8+central memory | CD8 | CAMBUS control | Super enhancers | Brain | 0.52 | 0.48 | 0.57 | 1.88.E-48 |
| CD8+central memory | CD8 | cOCRs | Super enhancers | Kidney | 4.07 | 3.60 | 4.59 | 2.50.E-113 |
| CD8+central memory | CD8 | CAMBUS control | Super enhancers | Kidney | 0.42 | 0.38 | 0.47 | 3.51.E-61 |
| CD8+central memory | CD8 | cOCRs | Super enhancers | Lung | 4.03 | 3.58 | 4.53 | 6.59.E-119 |
| CD8+central memory | CD8 | CAMBUS control | Super enhancers | Lung | 0.51 | 0.47 | 0.56 | 1.36.E-49 |
| CD8+central memory | CD8 | cOCRs | FANTOM enhancers | Hemato | 7.66 | 6.64 | 8.84 | 2.54.E-171 |
| CD8+central memory | CD8 | CAMBUS control | FANTOM enhancers | Hemato | 0.25 | 0.22 | 0.29 | 2.06.E-90 |
| CD8+central memory | CD8 | cOCRs | FANTOM enhancers | Brain | 7.01 | 6.00 | 8.18 | 2.33.E-134 |
| CD8+central memory | CD8 | CAMBUS control | FANTOM enhancers | Brain | 0.27 | 0.24 | 0.32 | 1.23.E-66 |
| CD8+central memory | CD8 | cOCRs | FANTOM enhancers | Kidney | 8.03 | 6.68 | 9.65 | 1.38.E-108 |
| CD8+central memory | CD8 | CAMBUS control | FANTOM enhancers | Kidney | 0.23 | 0.19 | 0.28 | 3.22.E-58 |
| CD8+central memory | CD8 | cOCRs | FANTOM enhancers | Lung | 7.46 | 6.39 | 8.72 | 1.03.E-141 |
| CD8+central memory | CD8 | CAMBUS control | FANTOM enhancers | Lung | 0.26 | 0.22 | 0.30 | 1.48.E-71 |
| CD8+effector memory | CD8 | cOCRs | Super enhancers | Blood | 8.41 | 7.14 | 9.92 | 4.51.E-142 |
| CD8+effector memory | CD8 | CAMBUS control | Super enhancers | Blood | 0.44 | 0.40 | 0.49 | 3.83.E-63 |
| CD8+effector memory | CD8 | cOCRs | Super enhancers | Brain | 5.48 | 4.69 | 6.41 | 1.10.E-100 |
| CD8+effector memory | CD8 | CAMBUS control | Super enhancers | Brain | 0.66 | 0.60 | 0.73 | 1.58.E-16 |
| CD8+effector memory | CD8 | cOCRs | Super enhancers | Kidney | 4.29 | 3.69 | 4.99 | 3.99.E-79 |
| CD8+effector memory | CD8 | CAMBUS control | Super enhancers | Kidney | 0.52 | 0.46 | 0.58 | 7.40.E-28 |
| CD8+effector memory | CD8 | cOCRs | Super enhancers | Lung | 4.35 | 3.74 | 5.05 | 7.62.E-83 |
| CD8+effector memory | CD8 | CAMBUS control | Super enhancers | Lung | 0.63 | 0.57 | 0.69 | 2.75.E-20 |
| CD8+effector memory | CD8 | cOCRs | FANTOM enhancers | Hemato | 8.17 | 6.84 | 9.75 | 1.48.E-118 |

| | | | | | | | | |
|---|---|---|---|---|---|---|---|---|
| CD8+effector memory | CD8 | CAMBUS control | FANTOM enhancers | Hemato | 0.33 | 0.28 | 0.39 | 1.02.E-40 |
| CD8+effector memory | CD8 | cOCRs | FANTOM enhancers | Brain | 7.30 | 6.02 | 8.86 | 2.29.E-90 |
| CD8+effector memory | CD8 | CAMBUS control | FANTOM enhancers | Brain | 0.36 | 0.30 | 0.43 | 3.82.E-29 |
| CD8+effector memory | CD8 | cOCRs | FANTOM enhancers | Kidney | 8.03 | 6.35 | 10.15 | 4.48.E-68 |
| CD8+effector memory | CD8 | CAMBUS control | FANTOM enhancers | Kidney | 0.31 | 0.25 | 0.39 | 7.99.E-25 |
| CD8+effector memory | CD8 | cOCRs | FANTOM enhancers | Lung | 7.79 | 6.42 | 9.46 | 1.28.E-95 |
| CD8+effector memory | CD8 | CAMBUS control | FANTOM enhancers | Lung | 0.33 | 0.28 | 0.40 | 1.48.E-32 |
| Naive B | B | cOCRs | Super enhancers | Blood | 21.23 | 15.01 | 30.04 | 9.14.E-67 |
| Naive B | B | CAMBUS control | Super enhancers | Blood | 0.40 | 0.34 | 0.47 | 6.39.E-31 |
| Naive B | B | cOCRs | Super enhancers | Brain | 14.44 | 10.39 | 20.06 | 6.86.E-57 |
| Naive B | B | CAMBUS control | Super enhancers | Brain | 0.54 | 0.46 | 0.63 | 3.22.E-15 |
| Naive B | B | cOCRs | Super enhancers | Kidney | 7.18 | 5.65 | 9.12 | 1.33.E-58 |
| Naive B | B | CAMBUS control | Super enhancers | Kidney | 0.46 | 0.38 | 0.55 | 3.83.E-16 |
| Naive B | B | cOCRs | Super enhancers | Lung | 5.62 | 4.37 | 7.23 | 2.22.E-41 |
| Naive B | B | CAMBUS control | Super enhancers | Lung | 0.60 | 0.51 | 0.70 | 2.74.E-10 |
| Naive B | B | cOCRs | FANTOM enhancers | Hemato | 4.74 | 3.26 | 6.89 | 4.13.E-16 |
| Naive B | B | CAMBUS control | FANTOM enhancers | Hemato | 0.56 | 0.39 | 0.80 | 1.55.E-03 |
| Naive B | B | cOCRs | FANTOM enhancers | Brain | 4.39 | 2.93 | 6.58 | 8.11.E-13 |
| Naive B | B | CAMBUS control | FANTOM enhancers | Brain | 0.59 | 0.40 | 0.88 | 9.47.E-03 |
| Naive B | B | cOCRs | FANTOM enhancers | Kidney | 5.23 | 3.20 | 8.55 | 4.39.E-11 |
| Naive B | B | CAMBUS control | FANTOM enhancers | Kidney | 0.49 | 0.30 | 0.79 | 3.58.E-03 |
| Naive B | B | cOCRs | FANTOM enhancers | Lung | 4.87 | 3.25 | 7.31 | 1.77.E-14 |
| Naive B | B | CAMBUS control | FANTOM enhancers | Lung | 0.54 | 0.36 | 0.79 | 1.82.E-03 |
| unsw memory B | B | cOCRs | Super enhancers | Blood | 8.73 | 7.46 | 10.22 | 2.49.E-160 |
| unsw memory B | B | CAMBUS control | Super enhancers | Blood | 0.20 | 0.18 | 0.23 | 1.94.E-133 |

| | | | | | | | | |
|---|---|---|---|---|---|---|---|---|
| unsw memory B | B | cOCRs | Super enhancers | Brain | 5.85 | 5.04 | 6.80 | 2.22.E-117 |
| unsw memory B | B | CAMBUS control | Super enhancers | Brain | 0.29 | 0.25 | 0.32 | 3.05.E-84 |
| unsw memory B | B | cOCRs | Super enhancers | Kidney | 4.65 | 4.02 | 5.38 | 2.71.E-95 |
| unsw memory B | B | CAMBUS control | Super enhancers | Kidney | 0.29 | 0.25 | 0.33 | 2.55.E-70 |
| unsw memory B | B | cOCRs | Super enhancers | Lung | 4.30 | 3.73 | 4.96 | 1.74.E-89 |
| unsw memory B | B | CAMBUS control | Super enhancers | Lung | 0.34 | 0.30 | 0.38 | 4.50.E-64 |
| unsw memory B | B | cOCRs | FANTOM enhancers | Hemato | 7.29 | 6.09 | 8.72 | 1.97.E-104 |
| unsw memory B | B | CAMBUS control | FANTOM enhancers | Hemato | 0.18 | 0.15 | 0.21 | 1.95.E-82 |
| unsw memory B | B | cOCRs | FANTOM enhancers | Brain | 6.68 | 5.50 | 8.13 | 7.19.E-81 |
| unsw memory B | B | CAMBUS control | FANTOM enhancers | Brain | 0.19 | 0.16 | 0.23 | 3.41.E-63 |
| unsw memory B | B | cOCRs | FANTOM enhancers | Kidney | 6.65 | 5.17 | 8.56 | 3.87.E-49 |
| unsw memory B | B | CAMBUS control | FANTOM enhancers | Kidney | 0.19 | 0.15 | 0.24 | 1.21.E-38 |
| unsw memory B | B | cOCRs | FANTOM enhancers | Lung | 7.43 | 6.12 | 9.03 | 1.94.E-90 |
| unsw memory B | B | CAMBUS control | FANTOM enhancers | Lung | 0.17 | 0.14 | 0.21 | 9.45.E-72 |
| sw memory B | B | cOCRs | Super enhancers | Blood | 7.73 | 6.77 | 8.83 | 2.13.E-199 |
| sw memory B | B | CAMBUS control | Super enhancers | Blood | 0.23 | 0.21 | 0.26 | 1.69.E-157 |
| sw memory B | B | cOCRs | Super enhancers | Brain | 5.34 | 4.70 | 6.07 | 1.82.E-144 |
| sw memory B | B | CAMBUS control | Super enhancers | Brain | 0.32 | 0.28 | 0.35 | 2.42.E-99 |
| sw memory B | B | cOCRs | Super enhancers | Kidney | 4.02 | 3.54 | 4.57 | 1.42.E-101 |
| sw memory B | B | CAMBUS control | Super enhancers | Kidney | 0.34 | 0.30 | 0.38 | 1.14.E-71 |
| sw memory B | B | cOCRs | Super enhancers | Lung | 4.09 | 3.62 | 4.63 | 2.37.E-112 |
| sw memory B | B | CAMBUS control | Super enhancers | Lung | 0.36 | 0.33 | 0.40 | 4.26.E-78 |
| sw memory B | B | cOCRs | FANTOM enhancers | Hemato | 7.55 | 6.48 | 8.79 | 1.05.E-148 |
| sw memory B | B | CAMBUS control | FANTOM enhancers | Hemato | 0.18 | 0.16 | 0.21 | 1.49.E-112 |
| sw memory B | B | cOCRs | FANTOM enhancers | Brain | 7.17 | 6.09 | 8.45 | 6.38.E-124 |
| sw memory B | B | CAMBUS control | FANTOM enhancers | Brain | 0.19 | 0.16 | 0.22 | 3.35.E-93 |

| | | | | | | | | |
|---|---|---|---|---|---|---|---|---|
| sw memory B | B | cOCRs | FANTOM enhancers | Kidney | 7.70 | 6.28 | 9.44 | 5.55.E-86 |
| sw memory B | B | CAMBUS control | FANTOM enhancers | Kidney | 0.17 | 0.14 | 0.21 | 8.68.E-66 |
| sw memory B | B | cOCRs | FANTOM enhancers | Lung | 7.46 | 6.32 | 8.81 | 1.87.E-123 |
| sw memory B | B | CAMBUS control | FANTOM enhancers | Lung | 0.18 | 0.15 | 0.21 | 1.54.E-93 |
| DN B | B | cOCRs | Super enhancers | Blood | 7.29 | 6.51 | 8.16 | 2.69.E-261 |
| DN B | B | CAMBUS control | Super enhancers | Blood | 0.18 | 0.17 | 0.20 | 6.97.E-234 |
| DN B | B | cOCRs | Super enhancers | Brain | 4.70 | 4.21 | 5.23 | 2.22.E-173 |
| DN B | B | CAMBUS control | Super enhancers | Brain | 0.27 | 0.25 | 0.30 | 6.83.E-144 |
| DN B | B | cOCRs | Super enhancers | Kidney | 3.66 | 3.27 | 4.09 | 7.25.E-114 |
| DN B | B | CAMBUS control | Super enhancers | Kidney | 0.32 | 0.28 | 0.35 | 2.01.E-95 |
| DN B | B | cOCRs | Super enhancers | Lung | 4.01 | 3.61 | 4.46 | 1.52.E-147 |
| DN B | B | CAMBUS control | Super enhancers | Lung | 0.31 | 0.28 | 0.34 | 6.01.E-121 |
| DN B | B | cOCRs | FANTOM enhancers | Hemato | 7.20 | 6.31 | 8.23 | 7.60.E-187 |
| DN B | B | CAMBUS control | FANTOM enhancers | Hemato | 0.16 | 0.14 | 0.18 | 5.14.E-165 |
| DN B | B | cOCRs | FANTOM enhancers | Brain | 6.89 | 5.97 | 7.95 | 7.22.E-154 |
| DN B | B | CAMBUS control | FANTOM enhancers | Brain | 0.17 | 0.14 | 0.19 | 1.53.E-135 |
| DN B | B | cOCRs | FANTOM enhancers | Kidney | 7.08 | 5.90 | 8.49 | 1.48.E-98 |
| DN B | B | CAMBUS control | FANTOM enhancers | Kidney | 0.16 | 0.13 | 0.19 | 3.60.E-87 |
| DN B | B | cOCRs | FANTOM enhancers | Lung | 7.13 | 6.16 | 8.25 | 1.87.E-153 |
| DN B | B | CAMBUS control | FANTOM enhancers | Lung | 0.16 | 0.14 | 0.19 | 1.50.E-135 |
| plasmablast | B | cOCRs | Super enhancers | Blood | 4.05 | 3.77 | 4.36 | 0.00.E+00 |
| plasmablast | B | CAMBUS control | Super enhancers | Blood | 0.42 | 0.40 | 0.45 | 1.96.E-203 |
| plasmablast | B | cOCRs | Super enhancers | Brain | 2.99 | 2.79 | 3.21 | 1.42.E-201 |
| plasmablast | B | CAMBUS control | Super enhancers | Brain | 0.58 | 0.54 | 0.61 | 1.12.E-82 |
| plasmablast | B | cOCRs | Super enhancers | Kidney | 2.42 | 2.23 | 2.62 | 1.86.E-104 |
| plasmablast | B | CAMBUS control | Super enhancers | Kidney | 0.56 | 0.53 | 0.60 | 2.03.E-60 |
| plasmablast | B | cOCRs | Super enhancers | Lung | 2.86 | 2.67 | 3.07 | 2.00.E-188 |
| plasmablast | B | CAMBUS control | Super enhancers | Lung | 0.55 | 0.52 | 0.58 | 1.50.E-97 |

| | | | | | | | | |
|---|---|---|---|---|---|---|---|---|
| plasmablast | B | cOCRs | FANTOM enhancers | Hemato | 5.36 | 4.89 | 5.88 | 1.49.E-276 |
| plasmablast | B | CAMBUS control | FANTOM enhancers | Hemato | 0.31 | 0.29 | 0.34 | 8.32.E-150 |
| plasmablast | B | cOCRs | FANTOM enhancers | Brain | 5.14 | 4.65 | 5.68 | 1.84.E-223 |
| plasmablast | B | CAMBUS control | FANTOM enhancers | Brain | 0.32 | 0.29 | 0.35 | 3.19.E-123 |
| plasmablast | B | cOCRs | FANTOM enhancers | Kidney | 5.15 | 4.52 | 5.85 | 3.90.E-137 |
| plasmablast | B | CAMBUS control | FANTOM enhancers | Kidney | 0.31 | 0.27 | 0.35 | 3.94.E-78 |
| plasmablast | B | cOCRs | FANTOM enhancers | Lung | 5.22 | 4.71 | 5.79 | 6.72.E-218 |
| plasmablast | B | CAMBUS control | FANTOM enhancers | Lung | 0.31 | 0.28 | 0.35 | 8.82.E-119 |
| plasmacytoid DC | DC | cOCRs | Super enhancers | Blood | 2.67 | 2.35 | 3.03 | 1.32.E-52 |
| plasmacytoid DC | DC | CAMBUS control | Super enhancers | Blood | 0.42 | 0.37 | 0.47 | 2.87.E-48 |
| plasmacytoid DC | DC | cOCRs | Super enhancers | Brain | 1.97 | 1.74 | 2.23 | 8.24.E-27 |
| plasmacytoid DC | DC | CAMBUS control | Super enhancers | Brain | 0.52 | 0.47 | 0.59 | 1.02.E-27 |
| plasmacytoid DC | DC | cOCRs | Super enhancers | Kidney | 1.79 | 1.55 | 2.06 | 2.39.E-15 |
| plasmacytoid DC | DC | CAMBUS control | Super enhancers | Kidney | 0.54 | 0.48 | 0.62 | 7.97.E-19 |
| plasmacytoid DC | DC | cOCRs | Super enhancers | Lung | 1.82 | 1.61 | 2.06 | 2.34.E-21 |
| plasmacytoid DC | DC | CAMBUS control | Super enhancers | Lung | 0.57 | 0.51 | 0.65 | 1.20.E-20 |
| plasmacytoid DC | DC | cOCRs | FANTOM enhancers | Hemato | 5.07 | 4.38 | 5.86 | 6.52.E-106 |
| plasmacytoid DC | DC | CAMBUS control | FANTOM enhancers | Hemato | 0.22 | 0.19 | 0.26 | 7.79.E-96 |
| plasmacytoid DC | DC | cOCRs | FANTOM enhancers | Brain | 4.61 | 3.93 | 5.40 | 1.18.E-78 |
| plasmacytoid DC | DC | CAMBUS control | FANTOM enhancers | Brain | 0.24 | 0.21 | 0.28 | 5.01.E-73 |
| plasmacytoid DC | DC | cOCRs | FANTOM enhancers | Kidney | 4.74 | 3.86 | 5.81 | 4.89.E-50 |
| plasmacytoid DC | DC | CAMBUS control | FANTOM enhancers | Kidney | 0.23 | 0.19 | 0.28 | 2.23.E-48 |
| plasmacytoid DC | DC | cOCRs | FANTOM enhancers | Lung | 4.77 | 4.06 | 5.62 | 1.45.E-78 |
| plasmacytoid DC | DC | CAMBUS control | FANTOM enhancers | Lung | 0.23 | 0.20 | 0.27 | 1.05.E-72 |
| myeloid DC | DC | cOCRs | Super enhancers | Blood | 3.24 | 2.87 | 3.67 | 9.42.E-78 |
| myeloid DC | DC | CAMBUS control | Super enhancers | Blood | 0.38 | 0.35 | 0.43 | 1.83.E-67 |

| | | | | | | | | |
|---|---|---|---|---|---|---|---|---|
| myeloid DC | DC | cOCRs | Super enhancers | Brain | 2.14 | 1.90 | 2.42 | 1.59.E-35 |
| myeloid DC | DC | CAMBUS control | Super enhancers | Brain | 0.51 | 0.45 | 0.56 | 9.23.E-36 |
| myeloid DC | DC | cOCRs | Super enhancers | Kidney | 1.87 | 1.63 | 2.15 | 1.22.E-18 |
| myeloid DC | DC | CAMBUS control | Super enhancers | Kidney | 0.55 | 0.48 | 0.62 | 2.29.E-21 |
| myeloid DC | DC | cOCRs | Super enhancers | Lung | 2.26 | 2.01 | 2.55 | 1.49.E-40 |
| myeloid DC | DC | CAMBUS control | Super enhancers | Lung | 0.50 | 0.45 | 0.56 | 2.22.E-36 |
| myeloid DC | DC | cOCRs | FANTOM enhancers | Hemato | 5.77 | 5.06 | 6.59 | 1.65.E-148 |
| myeloid DC | DC | CAMBUS control | FANTOM enhancers | Hemato | 0.22 | 0.20 | 0.25 | 1.64.E-121 |
| myeloid DC | DC | cOCRs | FANTOM enhancers | Brain | 4.82 | 4.15 | 5.61 | 6.39.E-93 |
| myeloid DC | DC | CAMBUS control | FANTOM enhancers | Brain | 0.25 | 0.22 | 0.29 | 1.59.E-81 |
| myeloid DC | DC | cOCRs | FANTOM enhancers | Kidney | 4.57 | 3.75 | 5.56 | 2.11.E-51 |
| myeloid DC | DC | CAMBUS control | FANTOM enhancers | Kidney | 0.25 | 0.21 | 0.30 | 1.87.E-48 |
| myeloid DC | DC | cOCRs | FANTOM enhancers | Lung | 5.05 | 4.34 | 5.88 | 6.77.E-97 |
| myeloid DC | DC | CAMBUS control | FANTOM enhancers | Lung | 0.24 | 0.21 | 0.28 | 3.14.E-83 |
| monocyte CD16+ | Mono | cOCRs | Super enhancers | Blood | 3.89 | 3.65 | 4.14 | 0.00.E+00 |
| monocyte CD16+ | Mono | CAMBUS control | Super enhancers | Blood | 0.30 | 0.28 | 0.32 | 0.00.E+00 |
| monocyte CD16+ | Mono | cOCRs | Super enhancers | Brain | 2.56 | 2.41 | 2.73 | 6.60.E-202 |
| monocyte CD16+ | Mono | CAMBUS control | Super enhancers | Brain | 0.43 | 0.41 | 0.46 | 5.97.E-172 |
| monocyte CD16+ | Mono | cOCRs | Super enhancers | Kidney | 2.40 | 2.24 | 2.57 | 1.69.E-135 |
| monocyte CD16+ | Mono | CAMBUS control | Super enhancers | Kidney | 0.45 | 0.42 | 0.48 | 9.92.E-116 |
| monocyte CD16+ | Mono | cOCRs | Super enhancers | Lung | 2.65 | 2.50 | 2.82 | 3.28.E-217 |
| monocyte CD16+ | Mono | CAMBUS control | Super enhancers | Lung | 0.42 | 0.40 | 0.45 | 5.80.E-186 |
| monocyte CD16+ | Mono | cOCRs | FANTOM enhancers | Hemato | 5.17 | 4.80 | 5.56 | 0.00.E+00 |
| monocyte CD16+ | Mono | CAMBUS control | FANTOM enhancers | Hemato | 0.21 | 0.20 | 0.23 | 0.00.E+00 |
| monocyte CD16+ | Mono | cOCRs | FANTOM enhancers | Brain | 4.79 | 4.40 | 5.21 | 2.17.E-291 |
| monocyte CD16+ | Mono | CAMBUS control | FANTOM enhancers | Brain | 0.23 | 0.21 | 0.25 | 1.52.E-263 |

| | | | | | | | | |
|---|---|---|---|---|---|---|---|---|
| monocyte CD16+ | Mono | cOCRs | FANTOM enhancers | Kidney | 4.68 | 4.18 | 5.23 | 5.94.E-162 |
| monocyte CD16+ | Mono | CAMBUS control | FANTOM enhancers | Kidney | 0.23 | 0.21 | 0.26 | 1.15.E-146 |
| monocyte CD16+ | Mono | cOCRs | FANTOM enhancers | Lung | 4.84 | 4.44 | 5.27 | 3.26.E-286 |
| monocyte CD16+ | Mono | CAMBUS control | FANTOM enhancers | Lung | 0.23 | 0.21 | 0.25 | 3.38.E-259 |
| monocyte CD16- | Mono | cOCRs | Super enhancers | Blood | 6.35 | 5.89 | 6.85 | 0.00.E+00 |
| monocyte CD16- | Mono | CAMBUS control | Super enhancers | Blood | 0.20 | 0.19 | 0.22 | 0.00.E+00 |
| monocyte CD16- | Mono | cOCRs | Super enhancers | Brain | 3.77 | 3.50 | 4.05 | 3.02.E-279 |
| monocyte CD16- | Mono | CAMBUS control | Super enhancers | Brain | 0.33 | 0.31 | 0.35 | 5.32.E-228 |
| monocyte CD16- | Mono | cOCRs | Super enhancers | Kidney | 3.11 | 2.87 | 3.37 | 1.57.E-171 |
| monocyte CD16- | Mono | CAMBUS control | Super enhancers | Kidney | 0.37 | 0.34 | 0.39 | 1.08.E-142 |
| monocyte CD16- | Mono | cOCRs | Super enhancers | Lung | 3.62 | 3.37 | 3.89 | 2.21.E-268 |
| monocyte CD16- | Mono | CAMBUS control | Super enhancers | Lung | 0.33 | 0.31 | 0.35 | 6.28.E-226 |
| monocyte CD16- | Mono | cOCRs | FANTOM enhancers | Hemato | 6.41 | 5.87 | 7.00 | 0.00.E+00 |
| monocyte CD16- | Mono | CAMBUS control | FANTOM enhancers | Hemato | 0.18 | 0.16 | 0.19 | 0.00.E+00 |
| monocyte CD16- | Mono | cOCRs | FANTOM enhancers | Brain | 6.22 | 5.63 | 6.86 | 4.70.E-289 |
| monocyte CD16- | Mono | CAMBUS control | FANTOM enhancers | Brain | 0.18 | 0.17 | 0.20 | 1.94.E-254 |
| monocyte CD16- | Mono | cOCRs | FANTOM enhancers | Kidney | 6.46 | 5.70 | 7.34 | 7.16.E-184 |
| monocyte CD16- | Mono | CAMBUS control | FANTOM enhancers | Kidney | 0.17 | 0.15 | 0.20 | 1.00.E-162 |
| monocyte CD16- | Mono | cOCRs | FANTOM enhancers | Lung | 6.10 | 5.51 | 6.75 | 3.65.E-268 |
| monocyte CD16- | Mono | CAMBUS control | FANTOM enhancers | Lung | 0.19 | 0.17 | 0.21 | 6.39.E-237 |
| nonclassical Monocyte | Mono | cOCRs | Super enhancers | Blood | 2.76 | 2.55 | 2.98 | 5.42.E-139 |
| nonclassical Monocyte | Mono | CAMBUS control | Super enhancers | Blood | 0.41 | 0.38 | 0.44 | 6.28.E-124 |
| nonclassical Monocyte | Mono | cOCRs | Super enhancers | Brain | 2.16 | 2.00 | 2.34 | 3.62.E-83 |
| nonclassical Monocyte | Mono | CAMBUS control | Super enhancers | Brain | 0.50 | 0.46 | 0.53 | 2.25.E-77 |
| nonclassical Monocyte | Mono | cOCRs | Super enhancers | Kidney | 1.95 | 1.78 | 2.13 | 6.34.E-48 |
| nonclassical Monocyte | Mono | CAMBUS control | Super enhancers | Kidney | 0.53 | 0.49 | 0.58 | 1.91.E-46 |
| nonclassical Monocyte | Mono | cOCRs | Super enhancers | Lung | 2.14 | 1.98 | 2.32 | 1.64.E-81 |
| nonclassical Monocyte | Mono | CAMBUS control | Super enhancers | Lung | 0.51 | 0.47 | 0.55 | 1.24.E-72 |

| | | | | | | | | |
|---|---|---|---|---|---|---|---|---|
| nonclassical Monocyte | Mono | cOCRs | FANTOM enhancers | Hemato | 4.58 | 4.18 | 5.02 | 1.59.E-233 |
| nonclassical Monocyte | Mono | CAMBUS control | FANTOM enhancers | Hemato | 0.25 | 0.23 | 0.27 | 7.79.E-209 |
| nonclassical Monocyte | Mono | cOCRs | FANTOM enhancers | Brain | 4.37 | 3.94 | 4.85 | 1.55.E-172 |
| nonclassical Monocyte | Mono | CAMBUS control | FANTOM enhancers | Brain | 0.25 | 0.23 | 0.28 | 7.71.E-161 |
| nonclassical Monocyte | Mono | cOCRs | FANTOM enhancers | Kidney | 4.27 | 3.73 | 4.88 | 5.50.E-98 |
| nonclassical Monocyte | Mono | CAMBUS control | FANTOM enhancers | Kidney | 0.25 | 0.22 | 0.29 | 8.37.E-93 |
| nonclassical Monocyte | Mono | cOCRs | FANTOM enhancers | Lung | 4.43 | 3.99 | 4.92 | 8.05.E-171 |
| nonclassical Monocyte | Mono | CAMBUS control | FANTOM enhancers | Lung | 0.25 | 0.22 | 0.27 | 2.71.E-158 |
| intermediate Monocyte | Mono | cOCRs | Super enhancers | Blood | 5.45 | 5.09 | 5.83 | 0.00.E+00 |
| intermediate Monocyte | Mono | CAMBUS control | Super enhancers | Blood | 0.22 | 0.21 | 0.24 | 0.00.E+00 |
| intermediate Monocyte | Mono | cOCRs | Super enhancers | Brain | 3.30 | 3.09 | 3.52 | 4.09.E-278 |
| intermediate Monocyte | Mono | CAMBUS control | Super enhancers | Brain | 0.36 | 0.34 | 0.38 | 7.78.E-231 |
| intermediate Monocyte | Mono | cOCRs | Super enhancers | Kidney | 2.91 | 2.70 | 3.13 | 4.25.E-180 |
| intermediate Monocyte | Mono | CAMBUS control | Super enhancers | Kidney | 0.38 | 0.35 | 0.41 | 2.47.E-154 |
| intermediate Monocyte | Mono | cOCRs | Super enhancers | Lung | 3.21 | 3.01 | 3.43 | 1.18.E-268 |
| intermediate Monocyte | Mono | CAMBUS control | Super enhancers | Lung | 0.36 | 0.34 | 0.38 | 2.49.E-231 |
| intermediate Monocyte | Mono | cOCRs | FANTOM enhancers | Hemato | 5.90 | 5.45 | 6.40 | 0.00.E+00 |
| intermediate Monocyte | Mono | CAMBUS control | FANTOM enhancers | Hemato | 0.19 | 0.18 | 0.21 | 0.00.E+00 |
| intermediate Monocyte | Mono | cOCRs | FANTOM enhancers | Brain | 5.73 | 5.24 | 6.27 | 0.00.E+00 |
| intermediate Monocyte | Mono | CAMBUS control | FANTOM enhancers | Brain | 0.20 | 0.18 | 0.21 | 1.38.E-282 |
| intermediate Monocyte | Mono | cOCRs | FANTOM enhancers | Kidney | 5.81 | 5.17 | 6.54 | 4.85.E-190 |
| intermediate Monocyte | Mono | CAMBUS control | FANTOM enhancers | Kidney | 0.19 | 0.17 | 0.21 | 3.85.E-170 |
| intermediate Monocyte | Mono | cOCRs | FANTOM enhancers | Lung | 5.77 | 5.26 | 6.32 | 3.70.E-307 |
| intermediate Monocyte | Mono | CAMBUS control | FANTOM enhancers | Lung | 0.19 | 0.18 | 0.21 | 2.79.E-275 |
| NK | NK | cOCRs | Super enhancers | Blood | 3.98 | 3.67 | 4.33 | 6.67.E-237 |
| NK | NK | CAMBUS control | Super enhancers | Blood | 0.27 | 0.25 | 0.29 | 5.41.E-229 |

| | | | | | | | | |
|---|---|---|---|---|---|---|---|---|
| NK | NK | cOCRs | Super enhancers | Brain | 2.62 | 2.42 | 2.84 | 1.81.E-124 |
| NK | NK | CAMBUS control | Super enhancers | Brain | 0.40 | 0.37 | 0.43 | 1.95.E-119 |
| NK | NK | cOCRs | Super enhancers | Kidney | 2.42 | 2.22 | 2.65 | 7.89.E-85 |
| NK | NK | CAMBUS control | Super enhancers | Kidney | 0.42 | 0.39 | 0.46 | 4.93.E-83 |
| NK | NK | cOCRs | Super enhancers | Lung | 2.74 | 2.53 | 2.96 | 1.53.E-135 |
| NK | NK | CAMBUS control | Super enhancers | Lung | 0.38 | 0.35 | 0.41 | 5.44.E-131 |
| NK | NK | cOCRs | FANTOM enhancers | Hemat o | 5.95 | 5.41 | 6.54 | 2.09.E-299 |
| NK | NK | CAMBUS control | FANTOM enhancers | Hemat o | 0.18 | 0.16 | 0.19 | 4.24.E-288 |
| NK | NK | cOCRs | FANTOM enhancers | Brain | 4.93 | 4.44 | 5.48 | 2.05.E-191 |
| NK | NK | CAMBUS control | FANTOM enhancers | Brain | 0.21 | 0.19 | 0.24 | 1.33.E-182 |
| NK | NK | cOCRs | FANTOM enhancers | Kidney | 5.22 | 4.58 | 5.96 | 1.73.E-133 |
| NK | NK | CAMBUS control | FANTOM enhancers | Kidney | 0.20 | 0.17 | 0.23 | 1.00.E-127 |
| NK | NK | cOCRs | FANTOM enhancers | Lung | 5.54 | 4.98 | 6.15 | 1.38.E-223 |
| NK | NK | CAMBUS control | FANTOM enhancers | Lung | 0.19 | 0.17 | 0.21 | 4.69.E-214 |
| PBMC | PBMC | cOCRs | Super enhancers | Blood | 4.59 | 4.40 | 4.79 | 0.00.E+00 |
| PBMC | PBMC | CAMBUS control | Super enhancers | Blood | 0.22 | 0.21 | 0.23 | 0.00.E+00 |
| PBMC | PBMC | cOCRs | Super enhancers | Brain | 3.12 | 2.99 | 3.25 | 0.00.E+00 |
| PBMC | PBMC | CAMBUS control | Super enhancers | Brain | 0.33 | 0.31 | 0.34 | 0.00.E+00 |
| PBMC | PBMC | cOCRs | Super enhancers | Kidney | 2.62 | 2.50 | 2.75 | 0.00.E+00 |
| PBMC | PBMC | CAMBUS control | Super enhancers | Kidney | 0.39 | 0.37 | 0.41 | 0.00.E+00 |
| PBMC | PBMC | cOCRs | Super enhancers | Lung | 2.93 | 2.81 | 3.05 | 0.00.E+00 |
| PBMC | PBMC | CAMBUS control | Super enhancers | Lung | 0.35 | 0.33 | 0.36 | 0.00.E+00 |
| PBMC | PBMC | cOCRs | FANTOM enhancers | Hemat o | 5.07 | 4.80 | 5.36 | 0.00.E+00 |
| PBMC | PBMC | CAMBUS control | FANTOM enhancers | Hemat o | 0.20 | 0.19 | 0.21 | 0.00.E+00 |
| PBMC | PBMC | cOCRs | FANTOM enhancers | Brain | 4.89 | 4.59 | 5.20 | 0.00.E+00 |
| PBMC | PBMC | CAMBUS control | FANTOM enhancers | Brain | 0.21 | 0.20 | 0.22 | 0.00.E+00 |

| | | | | | | | | |
|---|---|---|---|---|---|---|---|---|
| PBMC | PBMC | cOCRs | FANTOM enhancers | Kidney | 5.18 | 4.78 | 5.62 | 0.00.E+00 |
| PBMC | PBMC | CAMBUS control | FANTOM enhancers | Kidney | 0.20 | 0.18 | 0.21 | 0.00.E+00 |
| PBMC | PBMC | cOCRs | FANTOM enhancers | Lung | 5.02 | 4.71 | 5.35 | 0.00.E+00 |
| PBMC | PBMC | CAMBUS control | FANTOM enhancers | Lung | 0.20 | 0.19 | 0.22 | 0.00.E+00 |
| Neutrophil | Neutrophil | cOCRs | Super enhancers | Blood | 12.52 | 10.82 | 14.49 | 2.13.E-252 |
| Neutrophil | Neutrophil | CAMBUS control | Super enhancers | Blood | 0.08 | 0.07 | 0.09 | 1.14.E-253 |
| Neutrophil | Neutrophil | cOCRs | Super enhancers | Brain | 6.12 | 5.35 | 7.01 | 4.38.E-153 |
| Neutrophil | Neutrophil | CAMBUS control | Super enhancers | Brain | 0.16 | 0.14 | 0.19 | 3.28.E-153 |
| Neutrophil | Neutrophil | cOCRs | Super enhancers | Kidney | 4.44 | 3.89 | 5.07 | 8.50.E-109 |
| Neutrophil | Neutrophil | CAMBUS control | Super enhancers | Kidney | 0.23 | 0.20 | 0.26 | 1.89.E-108 |
| Neutrophil | Neutrophil | cOCRs | Super enhancers | Lung | 5.67 | 4.98 | 6.46 | 1.10.E-149 |
| Neutrophil | Neutrophil | CAMBUS control | Super enhancers | Lung | 0.18 | 0.16 | 0.20 | 1.93.E-150 |
| Neutrophil | Neutrophil | cOCRs | FANTOM enhancers | Hemato | 8.84 | 7.35 | 10.62 | 2.60.E-119 |
| Neutrophil | Neutrophil | CAMBUS control | FANTOM enhancers | Hemato | 0.11 | 0.09 | 0.14 | 1.47.E-118 |
| Neutrophil | Neutrophil | cOCRs | FANTOM enhancers | Brain | 7.99 | 6.56 | 9.73 | 7.39.E-95 |
| Neutrophil | Neutrophil | CAMBUS control | FANTOM enhancers | Brain | 0.13 | 0.10 | 0.15 | 3.15.E-94 |
| Neutrophil | Neutrophil | cOCRs | FANTOM enhancers | Kidney | 8.78 | 6.85 | 11.25 | 5.30.E-66 |

| | | | | | | | | |
|---|---|---|---|---|---|---|---|---|
| Neutrophil | Neutrophil | CAMBUS control | FANTOM enhancers | Kidney | 0.11 | 0.09 | 0.15 | 1.38.E-65 |
| Neutrophil | Neutrophil | cOCRs | FANTOM enhancers | Lung | 8.03 | 6.56 | 9.83 | 3.70.E-90 |
| Neutrophil | Neutrophil | CAMBUS control | FANTOM enhancers | Lung | 0.13 | 0.10 | 0.15 | 1.46.E-89 |

Supplementary Table 11 | meta analysis for tissue expressed super enhancers/FANTOM5 enhancers.
OR, odds ratio; CI_lo, lower 95% confidence interval; CI_hi, upper 95% confidence interval.

| OCR_type | enhancer_type | tissue | OR | CI_lo | CI_hi | P |
|---|---|---|---|---|---|---|
| cOCRs | Super_enhancers | Blood | 5.74 | 4.83 | 6.82 | 8.79.E-88 |
| CAMBUS_control | Super_enhancers | Blood | 0.32 | 0.27 | 0.36 | 7.42.E-59 |
| cOCRs | Super_enhancers | Brain | 3.79 | 3.21 | 4.49 | 2.00.E-54 |
| CAMBUS_control | Super_enhancers | Brain | 0.47 | 0.41 | 0.53 | 3.34.E-34 |
| cOCRs | Super_enhancers | Kidney | 3.15 | 2.79 | 3.57 | 2.94.E-74 |
| CAMBUS_control | Super_enhancers | Kidney | 0.45 | 0.41 | 0.49 | 1.26.E-77 |
| cOCRs | Super_enhancers | Lung | 3.32 | 3.02 | 3.66 | 3.35.E-131 |
| CAMBUS_control | Super_enhancers | Lung | 0.46 | 0.42 | 0.52 | 9.75.E-46 |
| cOCRs | FANTOM_enhancers | Hematopoietic | 6.51 | 6.09 | 6.95 | 5.79.E-673 |
| CAMBUS_control | FANTOM_enhancers | Hematopoietic | 0.24 | 0.21 | 0.26 | 5.29.E-145 |
| cOCRs | FANTOM_enhancers | Brain | 5.76 | 5.4 | 6.15 | 3.57.E-611 |
| CAMBUS_control | FANTOM_enhancers | Brain | 0.26 | 0.23 | 0.29 | 1.87.E-123 |
| cOCRs | FANTOM_enhancers | Kidney | 6.14 | 5.68 | 6.63 | 7.24.E-463 |
| CAMBUS_control | FANTOM_enhancers | Kidney | 0.24 | 0.22 | 0.26 | 2.58.E-170 |
| cOCRs | FANTOM_enhancers | Lung | 6.11 | 5.72 | 6.52 | 4.89.E-640 |
| CAMBUS_control | FANTOM_enhancers | Lung | 0.25 | 0.22 | 0.27 | 2.57.E-146 |

Supplementary Table 12 | meta analysis for tissue expressed super enhancers/FANTOM5 enhancers in relaxed cOCRs.
OR, odds ratio; CI_lo, lower 95% confidence interval; CI_hi, upper 95% confidence interval.

| OCR_type | enhancer_type | tissue | OR | CI_lo | CI_hi | P |
|---|---|---|---|---|---|---|
| cOCRs | Super_enhancers | Blood | 2.76 | 2.56 | 2.97 | 3.24E-161 |
| CAMBUS_control | Super_enhancers | Blood | 0.56 | 0.53 | 0.58 | 8.77E-126 |
| cOCRs | Super_enhancers | Brain | 1.76 | 1.69 | 1.82 | 1.01E-192 |
| CAMBUS_control | Super_enhancers | Brain | 0.77 | 0.75 | 0.78 | 1.64E-134 |
| cOCRs | Super_enhancers | Kidney | 1.66 | 1.59 | 1.73 | 2.96E-132 |
| CAMBUS_control | Super_enhancers | Kidney | 0.7 | 0.68 | 0.72 | 3.18E-115 |
| cOCRs | Super_enhancers | Lung | 1.85 | 1.76 | 1.94 | 4.93E-142 |
| CAMBUS_control | Super_enhancers | Lung | 0.69 | 0.67 | 0.71 | 9.08E-132 |
| cOCRs | FANTOM_enhancers | Brain | 3.48 | 3.29 | 3.69 | 3.78E-411 |
| CAMBUS_control | FANTOM_enhancers | Brain | 0.45 | 0.43 | 0.47 | 3.24E-233 |
| cOCRs | FANTOM_enhancers | Hemato | 4.52 | 4.2 | 4.86 | 6.75E-364 |
| CAMBUS_control | FANTOM_enhancers | Hemato | 0.39 | 0.37 | 0.42 | 6.45E-207 |
| cOCRs | FANTOM_enhancers | Kidney | 3.49 | 3.3 | 3.7 | 1.13E-395 |
| CAMBUS_control | FANTOM_enhancers | Kidney | 0.43 | 0.41 | 0.45 | 5.29E-213 |
| cOCRs | FANTOM_enhancers | Lung | 3.79 | 3.57 | 4.03 | 1.84E-395 |
| CAMBUS_control | FANTOM_enhancers | Lung | 0.42 | 0.4 | 0.45 | 4.52E-222 |

Supplementary Table 13 | phenotype list of S-LDSC

| Abbrevation | Trait Name | type | Reference | Sa |
|---|---|---|---|---|
| Celiac | Celiac Disease | Autoimmune | Dubois et al., 2010 Nat Genet | |
| CD | Crohn's Disease | Autoimmune | Jostins et al., 2012 Nature | |
| IBD | IBD | Autoimmune | Jostins et al., 2012 Nature | |
| SLE | Systemic Lupus Erythematosus | Autoimmune | Bentham et al., 2015 Nat Genet | |
| MS | Multiple Sclerosis | Autoimmune | IMS Genetics Consortium, 2011 Nature | |
| PBC | Primary Biliary Cirrhosis | Autoimmune | Cordell et al., 2015 Nat Commun | |
| RA | Rheumatoid Arthritis | Autoimmune | Okada et al., 2014 Nature | |
| T1D | Type 1 Diabetes | Autoimmune | Bradfield et al., 2011 Plos Genet | |
| UC | Ulcerative Colitis | Autoimmune | Jostins et al., 2012 Nature | |
| SLE (EAS) | Systemic Lupus Erythematosus | Autoimmune | Suetsugu et al., 2020 ARD | |
| AD (EAS) | Atopic | Autoimmune | Tanaka et al., 2022 JACI | |
| SSc (EAS) | Systemic Sclerosis | Autoimmune | Ishikawa et al., 2023 Nat Commun | |
| RA (EAS) | Rheumatoid Arthritis | Autoimmune | Ishigaki et al., 2022 Nat Genet | |
| TAK (EAS) | Takayasu Arteritis | Autoimmune | Terao et al., 2018 PNAS | |
| Alzheimer | Alzheimer's Disease | Psychiatric | Lambert et al., 2013 Nat Genet | |
| Anorexia | Anorexia | Psychiatric | Boraska et al., 2014 Mol Psych | |
| Autism | Autism Spectrum | Psychiatric | PGC Cross-Disorder Group, 2013 Lancet | |
| BP | Bipolar Disorder | Psychiatric | BIP Working Group of the PGC, 2011 Nat Genet | |
| DS | Depressive symptoms | Psychiatric | Okbay et al., 2016 Nat Genet | |
| Neuroticism | Neuroticism | Psychiatric | Okbay et al., 2016 Nat Genet | |
| SCZ | Schizophrenia | Psychiatric | SCZ Working Group of the PGC, 2014 Nature | |

| | | | |
|---|---|---|---|
| BMI1 | BMI (Speliotes 2010) | Metabolic | Speliotes et al., 2010 Nat Genet |
| CAD | Coronary Artery Disease | Metabolic | Schunkert et al., 2011 Nat Genet |
| FG | Fasting Glucose | Metabolic | Manning et. al., 2012 Nat Genet |
| HDL | HDL | Metabolic | Teslovich et al., 2010 Nature |
| LDL | LDL | Metabolic | Teslovich et al., 2010 Nature |
| TG | Triglycerides | Metabolic | Teslovich et al., 2010 Nature |
| T2D | Type 2 Diabetes | Metabolic | Morris et al., 2012 Nat Genet |
| Ever_Smoked | Ever Smoked | Others | TAG Consortium, 2010 Nat Genet |
| Height1 | Height (Lango Allen 2010) | Others | Lango Allen et al., 2010 Nature |
| SWB | Subject well being | Others | Okbay et al., 2016 Nat Genet |
| YED1 | Years of Education (Rietveld 2013) | Others | Rietveld et al., 2013 Science |
| YED2 | Years of Education (Okbay 2016) | Others | Okbay et al., 2016 Nature |

Supplementary Table 15 | fixed-effect meta analysis with five fold data sets to evaluate improving PRS model.

| Effect | StdErr | P-value | Direction |
|---|---|---|---|
| 0.00117 | 0.00025 | 3.2E-06 | +++++ |

Supplementary Table 16 | Pearson correlation of accessibility.

| name | subgroup | N | min | 1stQu | median | mean | 3rdQu | max | sd |
|---|---|---|---|---|---|---|---|---|---|
| Naive CD4 | CD4 | 2,704 | 0.2206 | 0.8480 | 0.9185 | 0.8570 | 0.9505 | 1 | 0.1578 |
| Th1 | CD4 | 2,809 | 0.4341 | 0.8993 | 0.9505 | 0.9157 | 0.9685 | 1 | 0.0900 |
| Th2 | CD4 | 2,809 | 0.1345 | 0.8427 | 0.9230 | 0.8575 | 0.9572 | 1 | 0.1694 |
| Th17 | CD4 | 2,916 | 0.3975 | 0.8960 | 0.9432 | 0.9060 | 0.9666 | 1 | 0.1124 |
| Tfh | CD4 | 2,916 | 0.2516 | 0.8671 | 0.9208 | 0.8771 | 0.9503 | 1 | 0.1358 |
| FraI | CD4 | 2,704 | 0.1788 | 0.8565 | 0.9268 | 0.8550 | 0.9536 | 1 | 0.1651 |
| FraII (aTreg) | CD4 | 2,809 | 0.7376 | 0.8866 | 0.9207 | 0.9137 | 0.9498 | 1 | 0.0465 |
| FraIII | CD4 | 2,809 | 0.3603 | 0.8936 | 0.9471 | 0.8896 | 0.9671 | 1 | 0.1314 |
| LAG3 Treg | CD4 | 2,704 | 0.2392 | 0.8927 | 0.9435 | 0.9008 | 0.9675 | 1 | 0.1373 |
| Memory CD4 | CD4 | 2,916 | 0.1570 | 0.8641 | 0.9298 | 0.8693 | 0.9587 | 1 | 0.1614 |
| Thx | CD4 | 2,916 | 0.1232 | 0.9043 | 0.9454 | 0.8796 | 0.9673 | 1 | 0.1766 |
| Naive CD8 | CD8 | 2,916 | 0.4804 | 0.8872 | 0.9369 | 0.8968 | 0.9566 | 1 | 0.0989 |
| CD8+effector | CD8 | 2,809 | 0.3238 | 0.8819 | 0.9324 | 0.8901 | 0.9582 | 1 | 0.1276 |
| CD8+central memory | CD8 | 2,704 | 0.1406 | 0.8611 | 0.9193 | 0.8558 | 0.9505 | 1 | 0.1673 |
| CD8+effector memory | CD8 | 2,916 | 0.1712 | 0.8323 | 0.9156 | 0.8585 | 0.9511 | 1 | 0.1517 |
| Naive B | B | 2,916 | 0.2738 | 0.7984 | 0.9044 | 0.8478 | 0.9452 | 1 | 0.1370 |
| unsw memory B | B | 2,809 | 0.2631 | 0.8232 | 0.9104 | 0.8605 | 0.9493 | 1 | 0.1324 |
| sw memory B | B | 2,704 | 0.3326 | 0.8400 | 0.9277 | 0.8758 | 0.9615 | 1 | 0.1304 |
| DN B | B | 2,704 | 0.3928 | 0.8747 | 0.9198 | 0.8899 | 0.9458 | 1 | 0.0937 |
| plasmablast | B | 2,704 | 0.4110 | 0.7420 | 0.8646 | 0.8322 | 0.9432 | 1 | 0.1321 |
| plasmacytoid DC | DC | 2,704 | 0.1365 | 0.8562 | 0.9457 | 0.8319 | 0.9687 | 1 | 0.2222 |
| myeloid DC | DC | 2,809 | 0.3243 | 0.9083 | 0.9648 | 0.9217 | 0.9788 | 1 | 0.1123 |
| monocyte CD16+ | Mono | 2,809 | 0.1176 | 0.8541 | 0.9463 | 0.8658 | 0.9685 | 1 | 0.1841 |
| monocyte CD16- | Mono | 2,809 | 0.1693 | 0.7920 | 0.8950 | 0.8305 | 0.9416 | 1 | 0.1652 |
| nonclassical Monocyte | Mono | 2,809 | 0.1600 | 0.9210 | 0.9592 | 0.8997 | 0.9778 | 1 | 0.1583 |

| | | | | | | | | | |
|---|---|---|---|---|---|---|---|---|---|
| intermediate Monocyte | Mono | 2,704 | 0.1050 | 0.7722 | 0.9093 | 0.8297 | 0.9505 | 1 | 0.1877 |
| NK | NK | 2,809 | 0.1854 | 0.9142 | 0.9564 | 0.8814 | 0.9760 | 1 | 0.1823 |
| PBMC | PBMC | 2,809 | 0.3494 | 0.8006 | 0.8938 | 0.8506 | 0.9435 | 1 | 0.1247 |
| Neutrophil | Neutrophil | 2,809 | 0.1745 | 0.8862 | 0.9171 | 0.8866 | 0.9366 | 1 | 0.1336 |

Supplementary Table 17 | Configuration of DeepSEA

| training | | |
|---|---|---|
| | sequence_length | 5000 |
| | center_bin_to_predict | 200 |
| | batch_size | 64 |
| | max_steps | 1000000 |
| | n_validation_samples | 32000 |
| | n_test_samples | 640000 |
| | blacklist | hg38 |
| | test_holdout | [chr8, chr9] |
| | validation_holdout | [chr6, chr7] |
| | feature_thresholds | 0.5 |
| | random_seed | 42 |

| *in-silico* mutagenesis | | |
|---|---|---|
| | sequence_length | 5000 |
| | batch_size | 64 |
| | blacklist | hg38 |
| | random_seed | 123 |

Supplementary Table 18 | genotype QC and imputation for identify caQTL

| | |
|---|---|
| Genotyping platform | Infinium ASA-24+v1.0 kit |
| SNP QC (exclusion criteria) | Call rate < 0.98<br>p-HWE < 1.0e-06<br>difference of allele frequency from reference panel > 0.03<br>monomorphic variants |
| Sample QC (exclusion criteria) | Call rate < 0.98<br>(For chrX nonPAR): Fraction of heterozygous genotype > 0.05 in male samples, < 0.05 in female samples |
| Imputation reference panel | 1KG phase3 v5 + Japanese WGS. 3,256 Japanese individuals in the BBJ and 2,504 individuals in the 1000 Genomes Proje[illegible] 2024) |

253

Supplementary Table 19 | The Number of cOCRs/relaxed cOCRs for each cell type and CAMBUS control.

| name | subgroup | cOCRs | CAMBUS_control | relaxed cOCRs | relaxed CAMBUS_control |
|---|---|---|---|---|---|
| Naive CD4 | CD4 | 1,266 | 251,807 | 12,736 | 229,229 |
| Th1 | CD4 | 2,435 | 216,873 | 11,033 | 198,589 |
| Th2 | CD4 | 2,301 | 247,926 | 12,632 | 227,366 |
| Th17 | CD4 | 3,857 | 225,990 | 11,558 | 208,027 |
| Tfh | CD4 | 1,584 | 248,749 | 12,585 | 226,514 |
| FraI | CD4 | 1,309 | 236,777 | 11,954 | 215,156 |
| FraII (aTreg) | CD4 | 1,526 | 234,176 | 11,952 | 215,123 |
| FraIII | CD4 | 3,416 | 197,572 | 10,114 | 182,043 |
| LAG3 Treg | CD4 | 4,094 | 193,214 | 9,949 | 179,079 |
| Memory CD4 | CD4 | 2,506 | 245,687 | 12,473 | 224,497 |
| Thx | CD4 | 3,423 | 207,262 | 10,575 | 190,350 |
| Naive CD8 | CD8 | 301 | 341,475 | 17,109 | 307,960 |
| CD8+effector | CD8 | 816 | 287,354 | 14,430 | 259,740 |
| CD8+central memory | CD8 | 1,156 | 264,327 | 13,320 | 239,747 |
| CD8+effector memory | CD8 | 729 | 284,215 | 14,298 | 257,346 |
| Naive B | B | 270 | 357,108 | 17,889 | 321,992 |
| unsw memory B | B | 788 | 325,980 | 16,349 | 294,263 |
| sw memory B | B | 1,062 | 328,042 | 16,471 | 296,474 |
| DN B | B | 1,432 | 322,364 | 16,199 | 291,576 |
| plasmablast | B | 3,174 | 220,609 | 11,286 | 203,139 |
| plasmacytoid DC | DC | 1,008 | 206,869 | 10,401 | 187,207 |
| myeloid DC | DC | 1,076 | 239,232 | 12,030 | 216,526 |
| monocyte CD16+ | Mono | 4,241 | 274,598 | 13,959 | 251,261 |
| monocyte CD16- | Mono | 3,035 | 344,692 | 17,404 | 313,269 |
| nonclassical Monocyte | Mono | 2,553 | 229,856 | 11,637 | 209,458 |

| | | | | | |
|---|---|---|---|---|---|
| intermediate Monocyte | Mono | 3,679 | 328,074 | 16,606 | 298,899 |
| NK | NK | 2,484 | 193,104 | 9,784 | 176,109 |
| PBMC | PBMC | 9,534 | 285,163 | 14,742 | 265,344 |
| Neutrophil | Neutrophil | 988 | 508,454 | 25,473 | 458,503 |

255

Supplementary Table 21 | The number of fine-mapped variant in each trait.

| cohort | hemato(leuko) | hemato(non_leuko) | non_hemato |
|---|---|---|---|
| EUR | 100,781 | 118,018 | 625,278 |
| EAS | 108,596 | 137,059 | 299,090 |

| cohort | Abbrevation | Phenotype | trait_type |
|---|---|---|---|
| EAS | APTT | Activated partial thromboplastin time | non_hemat |
| EAS | ALT | Alanine transaminase | non_hemat |
| EAS | ALB | Albumin | non_hemat |
| EAS | AG | Albumine globulin ratio | hemato(leuko) |
| EAS | ALP | Alkaline phosphatase | non_hemat |
| EAS | AST | Aspartate transaminase | non_hemat |
| EAS | BASO | Basophil count | hemato(leuko) |
| EAS | BS | Blood sugar | non_hemat |
| EAS | BUN | Blood urea nitrogen | non_hemat |
| EAS | BMI | Body mass index | non_hemat |
| EAS | CRP | C reactive protein | non_hemat |
| EAS | CL | Chloride | non_hemat |
| EAS | CK | Creatinine kinase | non_hemat |
| EAS | DBP | Diastolic blood pressure | non_hemat |
| EAS | EA | E/A ratio | non_hemat |
| EAS | EF | Ejection Fraction | non_hemat |
| EAS | EOSINO | Eosinophil count | hemato(leuko) |
| EAS | EGFR | Estimated glomerular filtration ratio | non_hemat |
| EAS | FBG | Fibrinogen | non_hemat |
| EAS | FS | Fractional shortening | non_hemat |
| EAS | GTP | Gamma glutamyl transpeptidase | non_hemat |

| | | | |
|---|---|---|---|
| EAS | HDLC | High density lipoprotein cholesterol | non_hemat |
| EAS | HT | Hematocrit | hemato(non_leuko) |
| EAS | HB | Hemoglobin | hemato(non_leuko) |
| EAS | HBA1C | Hemoglobin A1c | non_hemat |
| EAS | IVS | Intraventricular septum | non_hemat |
| EAS | LDH | Lactate dehydrogenase | non_hemat |
| EAS | LDL | Low-density lipoprotein cholesterol | non_hemat |
| EAS | LVDD | LV diastolic dimension | non_hemat |
| EAS | LVM | LV mass | non_hemat |
| EAS | LVMI | LV mass index | non_hemat |
| EAS | LVDS | LV systolic dimension | non_hemat |
| EAS | LYMPH | Lymphocyte count | hemato(leuko) |
| EAS | MG | Magnesium | non_hemat |
| EAS | MAP | Mean atrial pressure | non_hemat |
| EAS | MCH | Mean corpuscular hemoglobin | hemato(non_leuko) |
| EAS | MCHC | Mean corpuscular hemoglobin concentration | hemato(non_leuko) |
| EAS | MCV | Mean corpuscular volume | hemato(non_leuko) |
| EAS | MONO | Monocyte count | hemato(leuko) |
| EAS | NEUTRO | Neutrophil | hemato(leuko) |
| EAS | NAP | Non albmin protein | hemato(leuko) |
| EAS | P | Phosphate | non_hemat |
| EAS | PLT | Platelet count | hemato(non_leuko) |
| EAS | PW | Posterior wall thickness | non_hemat |
| EAS | K | Potassium | non_hemat |
| EAS | PT | Prothrombin time | non_hemat |
| EAS | PP | Pulse pressure | non_hemat |
| EAS | RBC | Red blood cell count | hemato(non_leuko) |
| EAS | RET | Reticulocite count | hemato(non_leuko) |
| EAS | SEG | Segment count | hemato(leuko) |
| EAS | CA | Serum calcium | non_hemat |

| | | | |
|---|---|---|---|
| EAS | SCR | Serum creatinine | non_hemat |
| EAS | NAT | Sodium | non_hemat |
| EAS | HEIGHT | Standing height | non_hemat |
| EAS | SBP | Systolic blood pressure | non_hemat |
| EAS | TBIL | Total billirubin | non_hemat |
| EAS | TC | Total cholesterol | non_hemat |
| EAS | TP | Total protein | non_hemat |
| EAS | TG | Triglyceride | non_hemat |
| EAS | UA | Uric acid | non_hemat |
| EAS | WBC | White blood cell count | hemato(leuko) |
| EAS | ZTT | Zinc sulfate turbidity test | non_hemat |
| EUR | AFib | Atrial fibrillation | non_hemat |
| EUR | AG | Albumin/Globulin ratio | hemato(leuko) |
| EUR | Age_at_Menarche | Age at menarche | non_hemat |
| EUR | Age_at_Menopause | Age at menopause | non_hemat |
| EUR | AID_Combined | Autoimmune disease (Phecode + Self-reported) | non_hemat |
| EUR | Alb | Albumin | non_hemat |
| EUR | ALP | Alkaline phosphatase | non_hemat |
| EUR | ALT | Alanine aminotransferase | non_hemat |
| EUR | Alzheimer_LTFH | Alzheimer disease (LTFH) | non_hemat |
| EUR | ApoA | Apolipoprotein A | non_hemat |
| EUR | ApoB | Apolipoprotein B | non_hemat |
| EUR | AST | Aspartate aminotransferase | non_hemat |
| EUR | Asthma | Asthma | non_hemat |
| EUR | Balding_Type4 | Balding Type 4 | non_hemat |
| EUR | Baso | Basophil count | hemato(leuko) |
| EUR | BFP | Body fat percentage | non_hemat |
| EUR | Blood_Clot_Lung | Blood clot in the lung | non_hemat |
| EUR | BMI | Body mass index | non_hemat |

| | | | |
|---|---|---|---|
| EUR | BrC | Breast cancer | non_hemat |
| EUR | BW | Body weight | non_hemat |
| EUR | Ca | Calcium | non_hemat |
| EUR | CAD | Coronary artery disease | non_hemat |
| EUR | Cholelithiasis | Cholelithiasis | non_hemat |
| EUR | CRC | Colorectal cancer | non_hemat |
| EUR | CRP | C-reactive protein | non_hemat |
| EUR | DBP | Diastolic blood pressure | non_hemat |
| EUR | Depression_GP | Seen doctor (GP) for nerves, anxiety, tension or depression | non_hemat |
| EUR | DVT | Blood clot in the leg | non_hemat |
| EUR | eBMD | Estimated heel bone mineral density | non_hemat |
| EUR | eGFR | Estimated glomerular filtration rate (serum creatinine) | non_hemat |
| EUR | eGFRcys | Estimated glomerular filtration rate (cystain C) | non_hemat |
| EUR | Eosino | Eosinophil count | hemato(leuko) |
| EUR | FedUp_Feelings | Fed-up feelings | non_hemat |
| EUR | FEV1FVC | FEV1/FVC ratio | non_hemat |
| EUR | Fibroblastic_Disorders | Fibroblastic disorders | non_hemat |
| EUR | GTP | Gamma-glutamyl transferase | non_hemat |
| EUR | Glaucoma_Combined | Glaucoma (Phecode + Self-reported) | non_hemat |
| EUR | Glucose | Glucose | non_hemat |
| EUR | Guilty_Feelings | Guilty feelings | non_hemat |
| EUR | Hb | Hemoglobin | hemato(non_leuko) |
| EUR | HbA1c | Hemoglobin A1c | non_hemat |
| EUR | HDLC | High density lipoprotein cholesterol | non_hemat |
| EUR | Height | Adult height | non_hemat |
| EUR | Ht | Hematocrit | hemato(non_leuko) |
| EUR | Hypothyroidism | Hypothyroidism | non_hemat |
| EUR | IBD | Inflammatory bowel disease | non_hemat |

| | | | |
|---|---|---|---|
| EUR | IGF1 | Insulin-like growth factor 1 | non_hemat |
| EUR | Inguinal_Hernia | Inguinal hernia | non_hemat |
| EUR | Insomnia | Insomnia | non_hemat |
| EUR | Irritability | Irritability | non_hemat |
| EUR | LDL | Low density lipoprotein cholesterol | non_hemat |
| EUR | LipoA | Lipoprotein A | non_hemat |
| EUR | Loneliness | Loneliness | non_hemat |
| EUR | LOY | Loss of Y | non_hemat |
| EUR | LYMPH | Lymphocyte count | hemato(leuko) |
| EUR | MAP | Mean arterial pressure | non_hemat |
| EUR | MCH | Mean corpuscular hemoglobin | hemato(non_leuko) |
| EUR | MCHC | Mean corpuscular hemoglobin concentration | hemato(non_leuko) |
| EUR | MCP | Multi-site chronic pain | non_hemat |
| EUR | MCV | Mean corpuscular volume | hemato(non_leuko) |
| EUR | Migraine_Self | Migraine (Self-reported) | non_hemat |
| EUR | Miserableness | Miserableness | non_hemat |
| EUR | Mono | Monocyte count | hemato(leuko) |
| EUR | Mood_Swings | Mood swings | non_hemat |
| EUR | Morning_Person | Morning person | non_hemat |
| EUR | Nervous_Feelings | Nervous feelings | non_hemat |
| EUR | Neuroticism | Neuroticism | non_hemat |
| EUR | Neutro | Neutrophil count | hemato(leuko) |
| EUR | Plt | Platelet count | hemato(non_leuko) |
| EUR | PP | Pluse pressure | non_hemat |
| EUR | PrC | Prostate cancer | non_hemat |
| EUR | RBC | Red blood cell count | hemato(non_leuko) |
| EUR | Risk_Taking | Risk taking | non_hemat |
| EUR | SBP | Systolic blood pressure | non_hemat |
| EUR | Sensitivity | Sensitivity | non_hemat |
| EUR | SHBG | Sex hormone binding globulin | non_hemat |

| | | | |
|---|---|---|---|
| EUR | Smoking_CPD | Smoking (cigarettes per day) | non_hemat |
| EUR | Smoking_Ever_Never | Smoking (ever vs never) | non_hemat |
| EUR | Suffer_from_Nerves | Suffer from nerves | non_hemat |
| EUR | T2D | Type 2 diabetes | non_hemat |
| EUR | T2D_BMI | Type 2 diabetes (adjusted by BMI) | non_hemat |
| EUR | TBil | Total bilirubin | non_hemat |
| EUR | TC | Total cholesterol | non_hemat |
| EUR | Tense | Tense | non_hemat |
| EUR | Testosterone | Testosterone | non_hemat |
| EUR | TG | Triglyceride | non_hemat |
| EUR | TP | Total protein | non_hemat |
| EUR | UA | Uric acid | non_hemat |
| EUR | Urea | Urea | non_hemat |
| EUR | VitD | Vitamin D | non_hemat |
| EUR | WBC | White blood cell count | hemato(leuko) |
| EUR | WHRadjBMI | Waist-to-hip ratio (adjusted by BMI) | non_hemat |
| EUR | Worrier | Worrier | non_hemat |
| EUR | Worry_Too_Long | Worry too long after embarrassment | non_hemat |

257

Supplementary Table 22 | The antibodies for FACS sorting

| Panel | marker | conjugate | clone | isotype | vender | Cat# | volume (ul) |
|---|---|---|---|---|---|---|---|
| CD4 | CD25 | AF488 | BC96 | Mouse IgG1, κ | Biolegend | 302616 | 5 |
| | LAG3 | PE | Polyclonal | Goat IgG | R&D | FAB2319P | 20 |
| | CD185(CXCR5) | PerCP-Cy5.5 | RF8B2 | Rat IgG2b, κ | BD | 562781 | 5 |
| | CD3 | PE-Cy7 | UCHT1 | Mouse IgG1, κ | Biolegend | 300420 | 0.25 |
| | CD196(CCR6) | APC | 11A9 | Mouse IgG1, κ | BD | 560619 | 2.5 |
| | CD45RA | APC-Cy7 | HI100 | Mouse IgG2b, κ | Biolegend | 304128 | 1 |
| | CD183(CXCR3) | BV421 | 1C6/CXCR3 | Mouse IgG1, κ | BD | 562558 | 5 |
| | CD4 | BV510 | RPA-T4 | Mouse IgG1, κ | BD | 740161 | 1 |
| | CCR4(CD194) | BV605 | L291H4 | Mouse IgG1, κ | Biolegend | 359418 | 5 |
| | CCR7(CD197) | BV711 | G043H7 | Mouse IgG2a, κ | Biolegend | 353228 | 7.5 |
| | CD8 | PE-CF594 | RPA-T8 | Mouse IgG1, κ | BD | 562282 | 0.5 |
| Mono/NK/DC | CD14 | FITC | M5E2 | Mouse IgG2a, κ | Biolegend | 301804 | 5 |
| | HLA-DR | PE | L243 | Mouse IgG2a, κ | Biolegend | 307606 | 1 |
| | CD16 | PerCP-Cy5.5 | 3G8 | Mouse IgG1, κ | Biolegend | 302028 | 3 |
| | CD3 | PE-Cy7 | UCHT1 | Mouse IgG1, κ | Biolegend | 300420 | 0.25 |
| | CD123 | APC | AC145 | Mouse IgG2aκ | Miltenyi Biotec | 130-113-322 | 10 |
| | CD56(NCAM) | APC-Cy7 | HCD56 | Mouse IgG1, κ | Biolegend | 318332 | 5 |
| | CD11c | BV421 | B-ly6 | Mouse IgG1, κ | BD | 562561 | 2.5 |
| | CD19 | BV510 | HIB19 | Mouse IgG1, κ | BD | 740164 | 5 |
| B/CD8 | CD27 | FITC | O323 | Mouse IgG1, κ | Biolegend | 302806 | 2 |
| | CD19 | PE | HIB19 | Mouse IgG1, κ | Biolegend | 302208 | 2 |
| | CD38 | PerCP-Cy5.5 | HIT2 | Mouse IgG1, κ | Biolegend | 303522 | 2 |

| | | | | | | | |
|---|---|---|---|---|---|---|---|
| | CD3 | PE-Cy7 | UCHT1 | Mouse IgG1, κ | Biolegend | 300420 | 0.25 |
| | CD45RA | APC | HI100 | Mouse IgG2b, κ | Biolegend | 304112 | 5 |
| | CD4 | APC-Cy7 | RPA-T4 | Mouse IgG1, κ | Biolegend | 300518 | 2 |
| | Human IgD | BV421 | IA6-2 | Mouse IgG2a, κ | BD | 562518 | 3 |
| | CD8 | BV510 | RPA-T8 | Mouse IgG1, κ | BD | 563256 | 5 |
| | CCR7 | BV711 | G043H7 | Mouse IgG2a, κ | Biolegend | 353228 | 7.5 |

Supplementary Table 23 | Configuration of DeepSEA and ChromBPNet

DeepSEA training

| | |
|---|---|
| sequence_length | 5000 |
| center_bin_to_predict | 200 |
| batch_size | 64 |
| max_steps | 1000000 |
| n_validation_samples | 32000 |
| n_test_samples | 640000 |
| blacklist | hg38 |
| test_holdout | [chr1, chr3, chr6] |
| validation_holdout | [chr8, chr20] |
| feature_thresholds | 0.5 |
| random_seed | 42 |
| stopping_criteria | ["roc_auc", 5] |

ChromBPNet training

| | |
|---|---|
| input_length | 2114 |
| output_length | 1000 |
| batch_size | 64 |
| max_epochs | 50 |
| early_stopping_patience | 5 |
| learning_rate | 0.001 |
| filters | 512 |
| n_dilation_layers | 8 |
| negative_sampling_ratio | 0.1 |
| max_jitter | 500 |
| outlier_threshold | 0.99 |
| test_holdout | [chr1, chr3, chr6] |
| validation_holdout | [chr8, chr20] |
| random_seed | 1234 |

DeepSEA *in-silico* mutagenesis

| | |
|---|---|
| sequence_length | 5000 |
| batch_size | 64 |
| blacklist | hg38 |
| random_seed | 123 |

ChromBPNet *in-silico* mutagenesis

| | |
|---|---|
| sequence_length | 2114 |
| batch_size | 512 |
| variant_schema | chrombpnet |
| mutation_effect | logFC |
| forward_only | FALSE |
| random_seed | 1234 |
